# Development of supramolecular anticoagulants with on-demand reversibility

**DOI:** 10.1101/2023.11.12.566735

**Authors:** Millicent Dockerill, Daniel J. Ford, Simona Angerani, Imala Alwis, Jorge Ripoll-Rozada, Rhyll E. Smythe, Joanna S.T. Liu, Pedro José Barbosa Pereira, Shaun P. Jackson, Richard J. Payne, Nicolas Winssinger

## Abstract

Drugs are administered at a dosing schedule set by their therapeutic index and termination of action is achieved by clearance and metabolism of the drug (hours to days for small molecules, weeks to months for biologics). In some cases, it is important to achieve a fast reversal of the drug’s action to overcome dangerous side effects or in response to unforeseen events. A case in point is for anticoagulant drugs. Here we report a general strategy to achieve on-demand reversibility by leveraging supramolecular assembly of drug fragments and showcase the approach with thrombin-inhibiting anticoagulants. In our supramolecular drug design, the action of the drug is reinforced by a dynamic hybridisation of peptide nucleic acids (PNAs) between drug fragments. We show that this design enables the generation of very potent bivalent direct thrombin inhibitors (*K*_i_ 74 pM) and this inhibition can be reversed through the use of a PNA antidote. We demonstrate that these supramolecular inhibitors exhibit potent anticoagulant activity *in vitro* and *in vivo* and that this activity can also be reversed on demand.

---

Anticoagulants are critically important therapies for the prevention or reversal of thrombotic events in patients and achieve their effect by reducing fibrin deposition by inhibiting fibrinogen proteolysis and/or platelet activation.^1^ One of the key targets of anticoagulant therapy is the protease thrombin (Factor IIa/FIIa), however, blockade of thrombin with direct (e.g., hirudin and argatroban) or indirect (heparins and warfarin) thrombin inhibitor therapy is contraindicated for several thrombotic disorders, e.g. stroke, due to the high risk of bleeding side effects that can be lethal. Indeed, many anticoagulants (particularly heparin and warfarin)^2^ require close clinical monitoring to prevent life threatening bleeding side effects. Despite this, it has been estimated that anticoagulant-related bleeding is responsible for 15% of all emergency hospital visits.^3^ Life threatening bleeding is the most concerning complication of anticoagulant therapy and strategies for the reversal of anticoagulation are therefore essential.^4^ A common strategy to reverse the effects of anticoagulants is the administration of non-specific reversal agents that involves the infusion of coagulation factors designed to overwhelm the effects of circulating anticoagulants.^5^ More recently monoclonal antibodies and recombinant FXa have been developed, which bind to a specific small molecule anticoagulant with high affinity (idarucizumab for dabigatran and andexanet alfa for apixaban, edoxaban, and rivaroxaban), thus reversing the inhibition of factor Xa (FXa) or thrombin (FIIa).^6,7^ Whilst both of these approaches are effective, there are limitations for their use and are associated with high cost.

Herein, we present a novel means to generate potent thrombin-inhibiting anticoagulants with on-demand reversibility through programmed supramolecular molecular assembly.^8^ Supramolecular entities rely on labile non-covalent interactions, and by their very nature are dynamic and reversible in response to specific environmental cues or stimuli by shifting equilibria in the system.^9^ These features of supramolecular systems have been elegantly applied to molecular recognition, catalysis, molecular motors, stimuli responsive polymers and drug discovery and delivery, but to our knowledge has not been realised for applications in medicinal chemistry and pharmacology (refs^10-14^). The strategy presented here is based on the ability to link two fragments by a reversible supramolecular interaction that are able to interact cooperatively with the target at two distinct sites (Fig. 1), with the formation of the active inhibitor instructed by the target. Disruption of the supramolecular interaction linking the two fragments results in a loss of cooperativity yielding a loss of inhibitory activity. Our design of supramolecular thrombin inhibitors made use of binary interactions directed to the active site of thrombin and to exosite II (the so-called heparin binding site), joined together by hybridized peptide nucleic acid (PNA) molecules.

**Figure 1.**
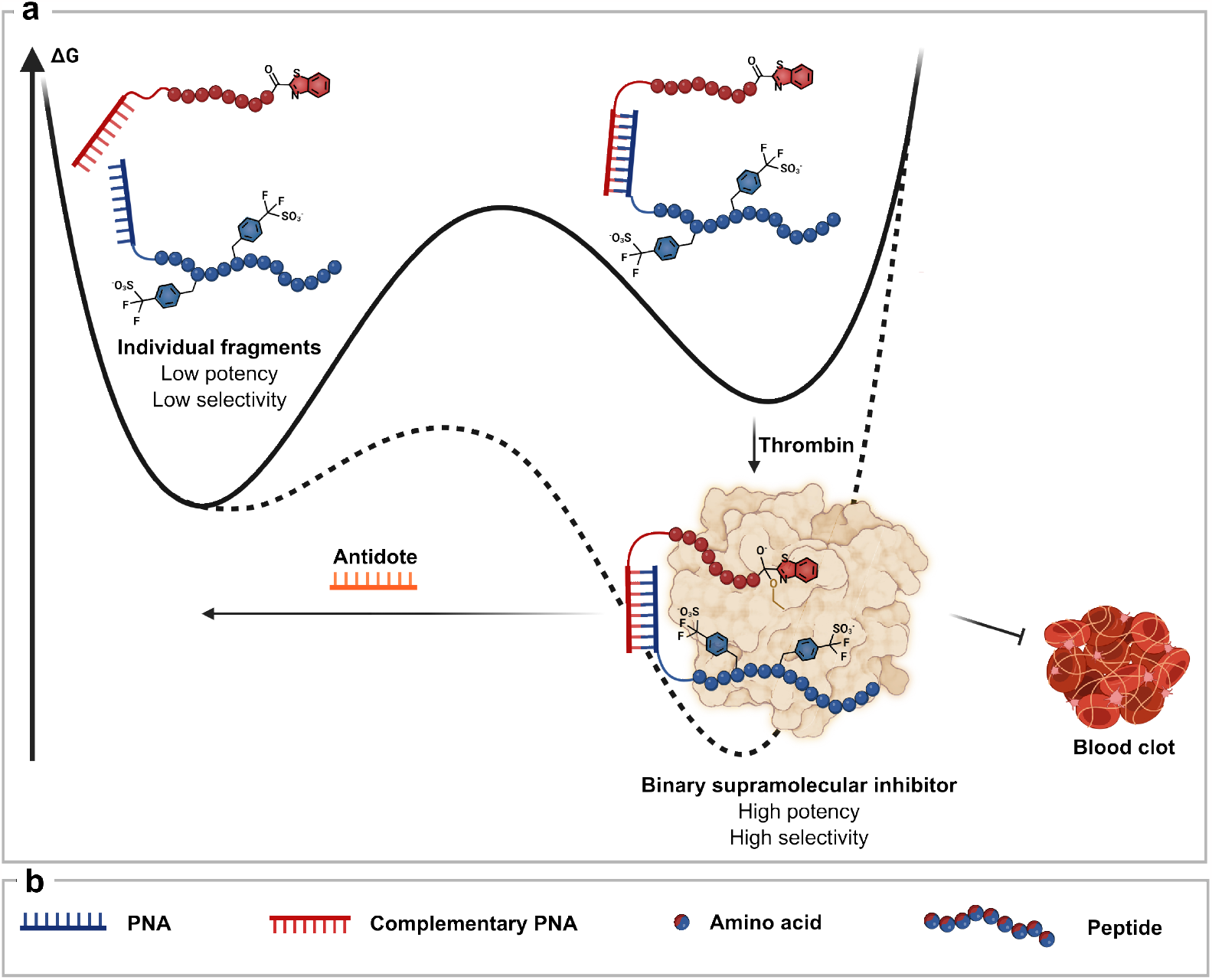
Supramolecular drug with on-demand reversibility. **a**. The assembly of the supramolecular drug is catalysed by the binding to thrombin which creates a highly potent and highly selective inhibitor from two compounds with low potency and selectivity. The inhibition of thrombin can be rapidly reversed by addition of an antidote. **b**. Legend of components represented in a.

The designed supramolecular assembly was inspired by thrombin inhibitors produced naturally by blood feeding (hematophagous) organisms such as leeches, ticks, mosquitoes and flies that secrete small protein thrombin inhibitors in their salivary glands to facilitate acquisition and digestion of a bloodmeal. These salivary proteins exhibit potent thrombin inhibition by interacting with two distinct binding sites on thrombin but this activity cannot be easily reversed due to their extremely high affinity for thrombin. At the outset we focused on hyalomin 1 (Hya1), a 59-residue sulfated protein secreted by the tick *Hyalomma marginatum rufipes* that shares sequence similarity to other tick anticoagulants proteins, but is the most potent thrombin inhibitor in the family (*K*_i_ = 5.4 pM). Analysis of the X-ray crystal structures of several of these proteins complexed with thrombin (e.g., tick-derived madanin-1 (PDB 5L6N)^15^ and TTI from the tsetse fly (PDB 6TKG))^16^ together with the thrombin inhibitory data suggested that the potent inhibition exhibited by these molecules was derived from interactions at two loci of thrombin, the active site and exosite II, separated by 20-30 Å (Extended Data, Fig. 1b). We reasoned that we could leverage an established ketobenzothiazole-containing mechanism-based pan-serine protease inhibitor for active site targeting that forms reversible covalent (hemiketal) intermediates with serine proteases, but is not selective for thrombin.^17^ For the peptide targeting exosite II, we investigated sequences from several salivary proteins from hematophagous organisms that possess sulfotyrosine residues as a common post-translational modification (PTM) that has been shown to enhance activity (Extended Data, Fig. 1c).^15,16,18^ Considering the reported lability of the tyrosine sulfate PTM, we opted for a synthetic analogue of the natural modification, namely (sulfono(difluoro)methyl-phenylalanine: F2Smp)).^19^ For the link between the two binding motifs in our supramolecular anticoagulant, we chose to employ the synthetic DNA mimetic PNA,^20^ based on the tunability of the hybridisation dynamics of this molecular class to provide anticoagulant reversibility, its metabolic stability, and the compatibility of its chemistry with peptide synthesis.^21^

We first used solid-phase synthesis to prepare the mechanism-based active site targeting peptide fragment **A** (A1: derived from Hya1 fused to a ketobenzothiazole warhead) linked to an 8-mer PNA sequence and fragment **E** (E1: derived from the exosite II binding region of TTI) linked to the complementary 8-mer PNA (Fig. 2a, see Extended Data, Fig. 2 for detailed chemical structure of the main compounds used in the study). Given the known importance of two native negatively charged sulfotyrosine residues for interaction with the heparin-binding exosite II in TTI, we incorporated two difluorosulfonomethylphenylalanine (F2Smp) residues as stable mimics in fragment **E1**. Fragment **A1** showed moderate inhibitory activity against thrombin (*K*_i_ 58.7 nM) in a fluorogenic thrombin-activity assay, while E1 alone possessed no inhibitory activity (Fig. 2b). However, an 800-fold enhancement of activity was observed when both components were mixed together using the 8-mer PNA supramolecular connection, with **A1-E1** exhibiting a *K*_i_ of 74 pM (Fig. 2b). Pleasingly, this supramolecular inhibitor also gained selectivity for thrombin when tested against a panel of proteases present in the coagulation pathway including FXa, FXIa, FXIIa and PK (> 1000-fold, Fig. 2c). It is noteworthy that, like thrombin, the substrate specificity for factor FXa and FXIa also strongly favour Arg at P1^22,23^ but only thrombin benefits from the binary interaction of the supramolecular drug, resulting in

**Figure 2.**
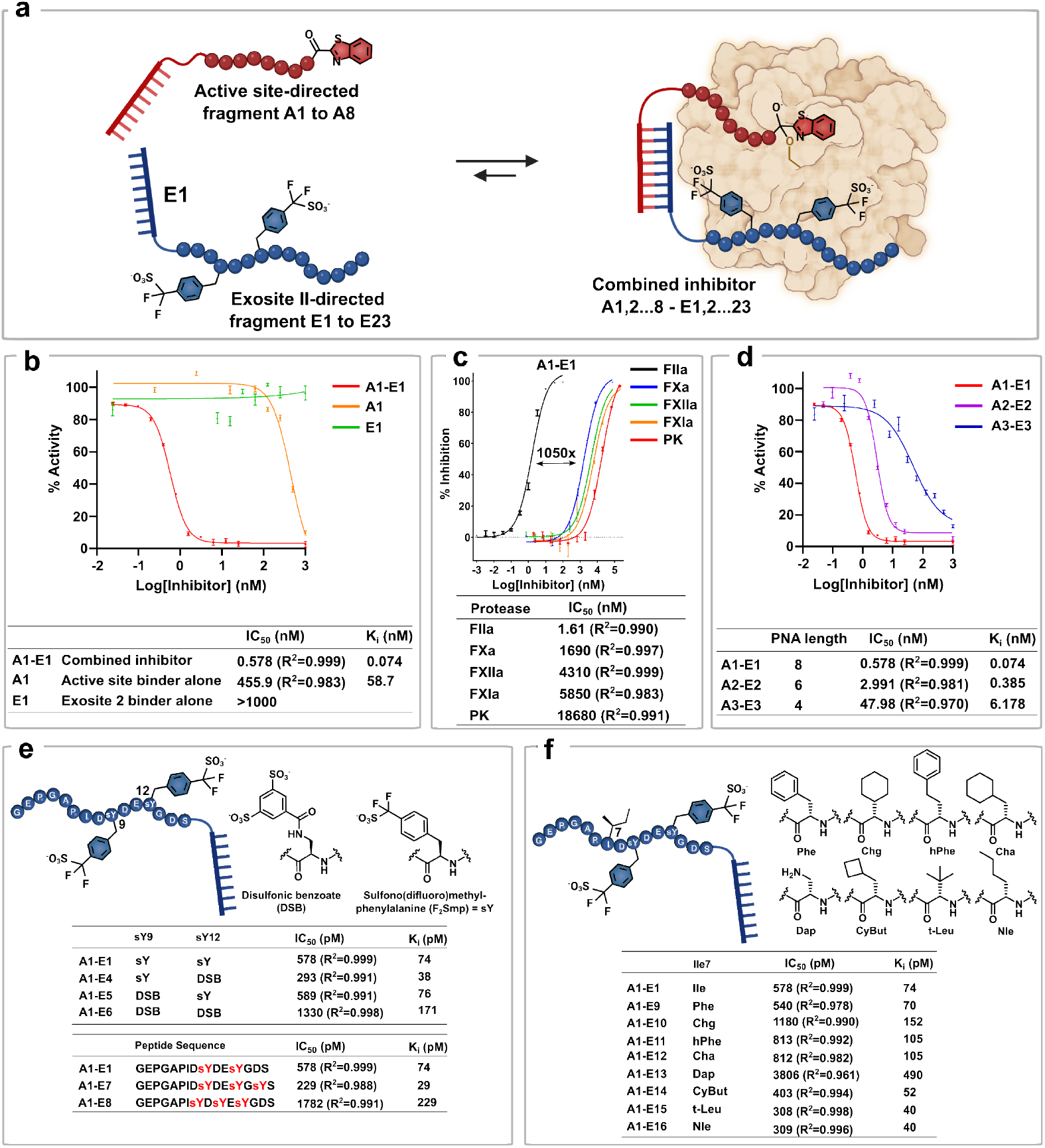
Inhibitor development. **a**. Schematic representation of the cooperative dynamic drug assembly. The inhibitors disclosed in this study are comprised of two fragments: the active site-directed fragments numbered A1 to A8 and the exosite II-directed fragment which are numbered E1 to E23. The combination of the two fragments yields a potent inhibitor named as the combination of the two assembled fragments (e.g., A1-E1 is the combination of active site fragment A1 and exosite II fragment E1). **b**. Thrombin inhibition data for the combined inhibitor versus the two fragments alone. **c**. Selectivity data for **A1-E1** against a panel of common proteases. **d**. Effect of PNA length on inhibition. **e**. SAR data of the exosite II binder by varying charge. **f**. SAR data of the exosite II binder by varying hydrophobic amino acids instead of isoleucine.

>1000-fold selectivity. To further investigate the supramolecular connectivity between the two fragments, we reduced the length of PNA from 8-mer to 6-mer or 4-mer, while keeping the overall distance equal. This led to a progressive loss of activity (Fig. 2d). However, the assembly composed of the shortest supramolecular linker (4-mer: **A3-E3**) was still 10-fold more potent than the active site inhibitor alone (**A1**). Taken together, these data support a cooperative interplay between the supramolecular interaction of the PNA and the inhibition of thrombin through engagement with both the active site and exosite II. The hybridisation K_D_ of the 4mer PNA was measured by SPR to be 4.14 *µ*M at 25 °C (Extended Data, Fig. 3), yet the supramolecular tether still yields a benefit at concentrations well below its K_D_. A cooperativity in the inhibition is observed if the equilibrium re-binding of the active site ligand is faster in the supramolecular assembly-enzyme complex than the dissociation of the supramolecular tether. It stands to reason that the longer PNA with slower *k*_off_ yields better cooperativity.

**Figure 3.**
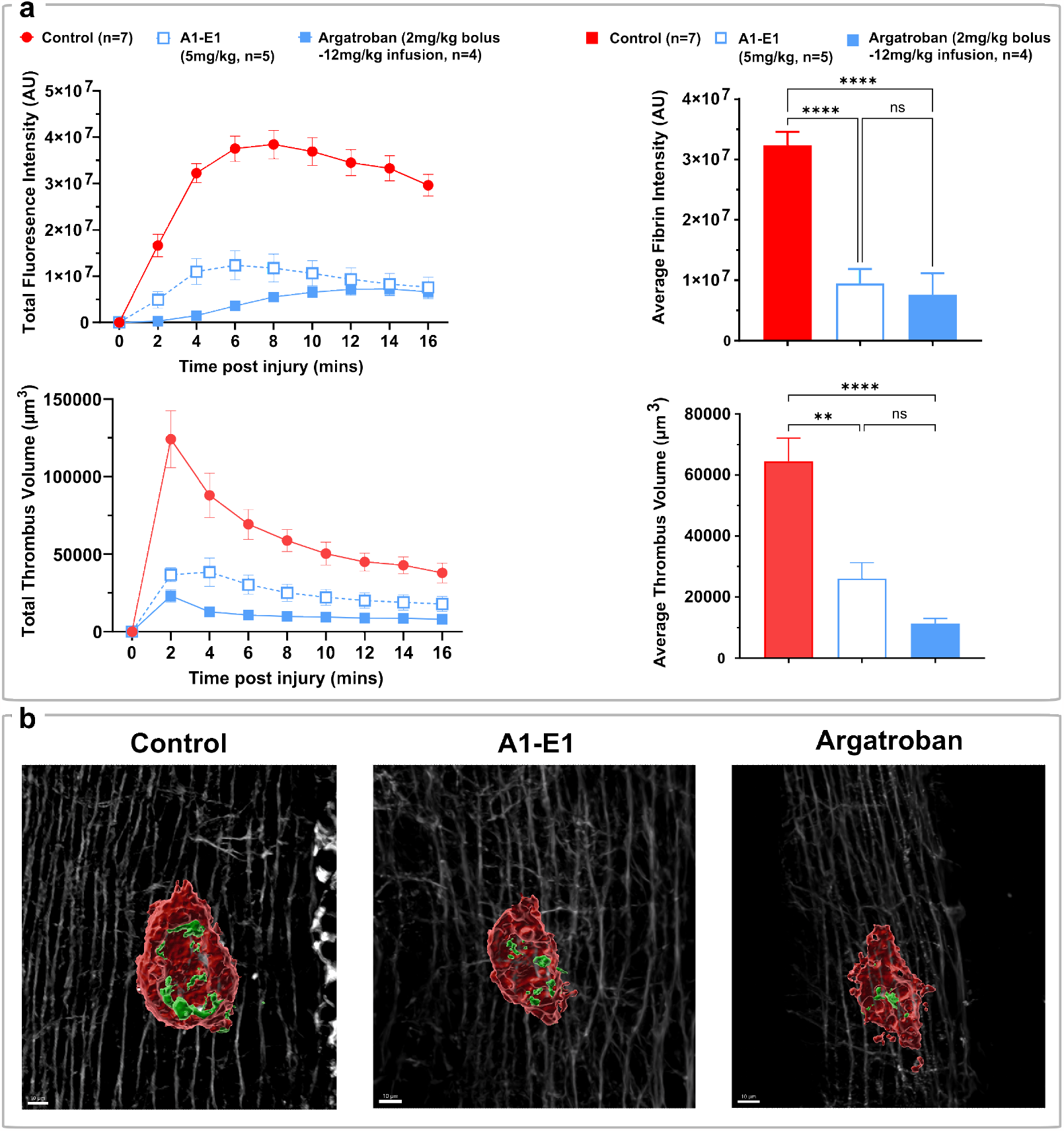
*In-vivo* inhibition of thrombin. **a**. Time course of fibrin fluorescence intensity and total thrombus volume (left), and average fibrin intensity and average thrombus volume for control group (n=7), argatroban treated cohort (n=4, 2mg/mL bolus followed by 12 mg/kg infusion) and **A1-E1** treated cohort (n=5, 5 mg/kg bolus). **b**. Exemplar image of thrombus 15 minutes after needle injury without inhibitor (left), with argatroban (centre) and with **A1-E1** (right). Platelets are shown in red, fibrin in green and collagen in the background in white, scale bar is 10 μm.

The use of the PNA as a supramolecular tether also provides the opportunity to quickly assemble analogues and perform structure-activity studies since new combinations can be generated simply by mixing the binary ligands. We first explored other stable sulfotyrosine mimics in the exosite II binding fragment by incorporating disulfonic benzoate (DSB) in lieu to F2Smp into the **E** fragment to generate **E4** (sY12→DSB), **E5** (sY9→DSB) and **E6** (sY9,12→DSB) that could be used form supramolecular assemblies with active site binding fragment **A1** by simple mixing (Fig 2e).^16^ Inclusion of DSB in place of F2Smp at position 12 led to a two-fold gain of activity (**A1-E4**, Fig. 2e) but replacement of both F2Smp residues with DSB moieties led to a decrease of inhibition (A1-E6, Fig. 2e). The position and number of sulfotyrosine mimics also had a strong impact (A1-E1 vs A1-E7,8, Fig. 2e). We next performed an alanine scan of the peptide sequence of **E1** that targeted exosite II. This revealed an isoleucine residue at position 7 (Ile7) as a hot spot (Extended Data, Fig. 4a), an observation consistent with the structure of the TTI-thrombin complex (PDB 6TKG, Extended Data, Fig. 4b)^16^, which shows this isoleucine filling a hydrophobic pocket. A moderate (ca. 2-fold) gain in activity could be achieved with substitution for hydrophobic non-proteinogenic amino acids (e.g., t-Leu or Nle in A1-E15 or A1-E16, respectively).

**Figure 4.**
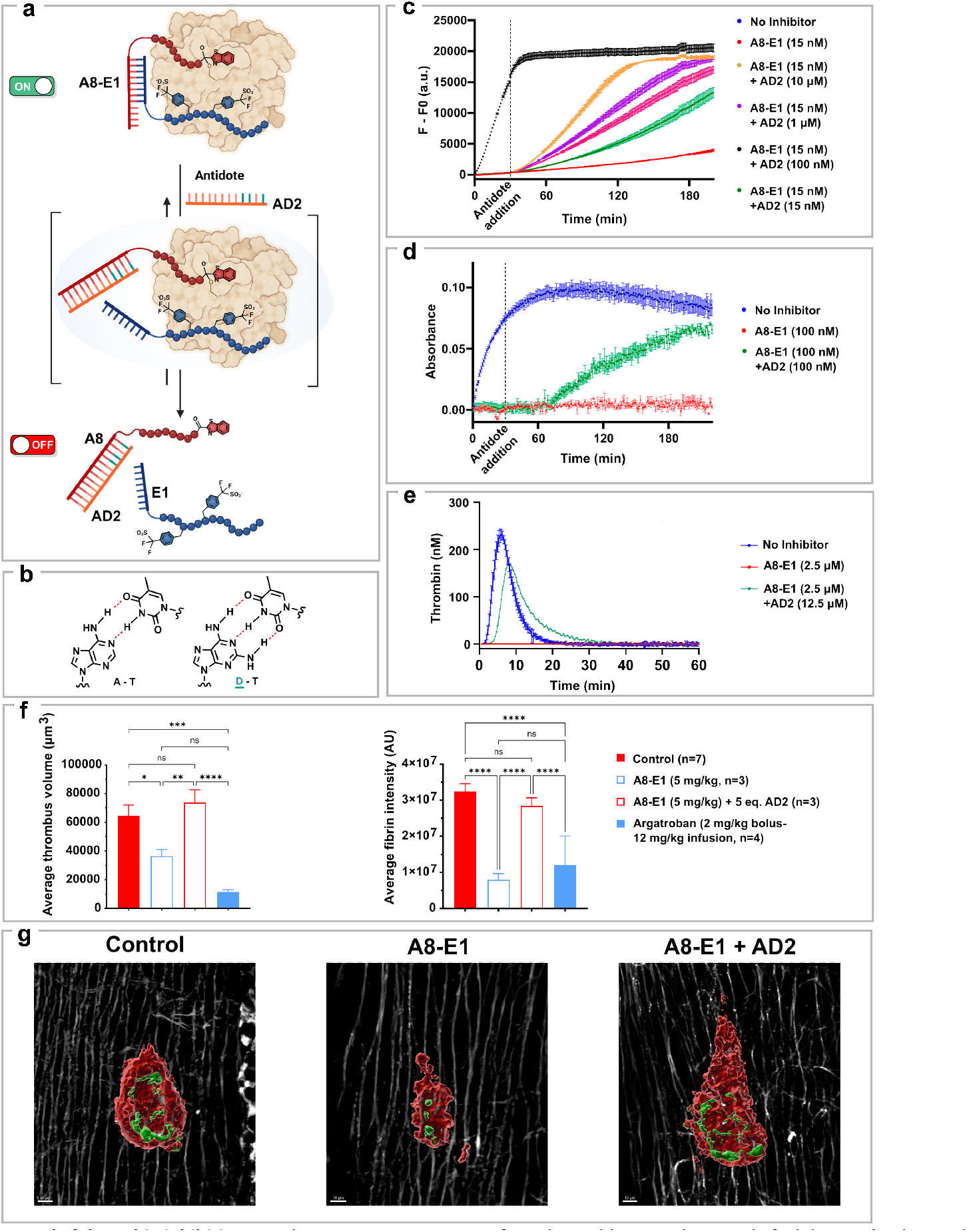
Reversal of thrombin inhibition. **a**. Schematic representation of antidote addition and reversal of inhibition. **b**. Chemical structures of adenine and diaminopurine forming hydrogen bonds with thymine. **c**. Fluorogenic assay data showing the reversal of thrombin inhibition by addition of different concentrations of antidote after 30 minutes of inhibition. **d**. Fibrinogen assay data showing the reversal of thrombin inhibition by addition of antidote (1eq.) after 30 minutes of inhibition. **e**. Calibrated Automated Thrombogram (CAT) of **A8-E1** with and without antidote. **f**. Average fibrin intensity and average thrombus volume for control group (n=7), argatroban treated cohort (n=4, 2mg/mL bolus followed by 12 mg/kg infusion), **A8-E1** treated cohort (n=3, 5 mg/kg bolus) and **A8-E1+AD1** treated cohort (n=3, 5mg/kg + 5 molar eq. antidote). **g**. Exemplar image of thrombus 15 minutes after needle injury without inhibitor (left), with **A8-E1** (centre) and with **A8-E1+AD1** (right). Platelets are shown in red, fibrin in green and collagen in white, scale bar is 10 μm.

Having established the feasibility of the supramolecular inhibitor concept, we selected **A1-E1** as a lead to profile in subsequent biochemical assays and for anticoagulant activity *in vitro*. Towards this end, we first investigated the inhibition of fibrinogen proteolysis, whereby **A1-E1** exhibited complete inhibition at 100 nM (Extended Data, Fig. 5a) whilst **A1** or **E1** alone were comparable to no inhibitor. Having demonstrated **A1-E1** was able to prevent fibrinogen proteolysis *in vitro* we next turned our attention to an activated partial thromboplastin time (aPTT) assay in both human and mouse plasma. aPTT assays are routine tests carried out by physicians and is an indicator of the function of coagulation factors in the intrinsic and common pathways, and effective inhibition of thrombin is expected to lengthen the time plasma takes to clot. A clinically significant increase in clotting is said to be 2-fold. Pleasingly **A1-E1** exhibited a therapeutically significant prolongation of clotting time in both human and mouse plasma at a concentration as low as 250 nM (Extended Data, Fig. 5b). We next investigated the effects of **A1-E1** on thrombin generation in a calibrated automated thrombogram (CAT). The CAT employs a fluorogenic thrombin substrate, thus allowing measurement of thrombin formation in plasma in real time. This is of particular importance since thrombin generation is a dynamic process, the coagulation cascade has many feedback loops and inhibitory pathways that are all directly or indirectly influenced by the developing thrombin concentration and thrombin plays a central and pivotal role throughout the whole process. Additionally, and in contrast to aPTT assays, the CAT allows for a large variation in the concentration and character of the trigger used and can therefore be implemented to detect subtle differences between thrombin inhibitors. **A1-E1** potently inhibited thrombin activity in both the initiation phase and propagation phase of coagulation and was able to completely inhibit thrombin activity at 2.5 *µ*M (Extended Data, Fig. 6).

Having determined that our supramolecular anticoagulant potently inhibited thrombin activity and possessed anticoagulant activity *in vitro*, we next investigated whether **A1-E1** would be effective at inhibiting thrombus formation *in vivo*. To determine a suitable dose for our *in vivo* efficacy study we utilised an *ex vivo* aPTT assay. Briefly, **A1-E1** was administered intravenously to mice at 2.5 or 5 mg/kg and blood samples were collected at 5, 15 and 45 mins respectively. Clotting times were then measured using a standard aPTT protocol and showed that a single 5 mg/kg bolus was effective at prolonging the aPTT >2 fold for 30 mins (Extended Data, Fig. 5c). We next assessed the *in vivo* efficacy of the supramolecular anticoagulant **A1-E1** compared to standard of care argatroban in a localised needle injury model.^24^ The injury leads to both fibrin formation and platelet aggregation in thrombus formation, which were visualized by Alexa-647 α-fibrin and Dylight-488 αGBP1bβ, respectively.^25^ Owing to its short half-life *in vivo*, argatroban was dosed at 3.9 μmol/kg (2 mg/kg) IV bolus followed by an infusion at 24 μmol/kg over 60 minutes (12 mg/kg, total dose: 27.9 μmol/kg). **A1-E1** was dosed twice at 0.63 μmol/kg (5mg/kg) IV bolus 30 minutes apart (total dose: 1.3 μmol/kg). Both **A1-E1** and argatroban showed significant decrease in both fibrin formation and thrombus size (Fig. 3). Following treatment with the supramolecular anticoagulant **A1-E1**, followed by injury, we observed near complete inhibition of fibrin deposition at the site of injury when compared to control injuries (Fig. 3). Pleasingly, we also observed that **A1-E1** achieved a similar level of anticoagulation to a bolus infusion of argatroban at the 5 mg/kg dosing regimen (Fig. 3). On a molarity basis, **A1-E1** yielded comparable results to the standard of care (argatroban) at 24-fold lower drug loading indicating that the potent inhibitory activity observed *in vitro* translates *in vivo*.

Having established promising *in vivo* efficacy for our supramolecular inhibitor, we turned our attention to investigate the ability to reverse the anticoagulant activity with an antidote. Given the non-covalent nature of the supramolecular linker between the active site and exosite II binding entities, we rationalized that the inhibition could be disrupted by competing for the hybridization. To favour the equilibrium towards the dissociation of the binary fragments, the competitor PNA was designed to incorporate diaminoapurines (D) instead of adenine (A), since oligomers containing D form more stable duplexes with their complementary strand than oligomers containing A.^26^ While this competitor **(AD1)** functioned as an effective antidote by reversing inhibition, the kinetics of the antidote were deemed too slow at low concentrations (1-10 μM, Extended Data, Fig. 7). Mindful of the observed cooperativity between target interaction and hybridization, we introduced a toehold sequence^27^ on the supramolecular connector **(A8-E1)** to achieve a larger equilibrium shift in the hybridization with **AD2**, a 12-mer PNA (Fig. 4a-b). Following the kinetic progress of the reaction in real time with a fluorogenic substrate, we observed the ability to switch from complete inhibition (15 nM of binary inhibitor) to ca. 40% of the uninhibited thrombin activity within 30 min using 10 μM of antidote (Fig. 4c). Using lower concentration of antidote resulted in more progressive restoration of thrombin’s activity. Using just 1 equivalent of antidote was sufficient to restore *ca*. 20% of thrombin catalytic activity within 90 minutes. These observations were also validated in the fibrinogen clotting and CAT assays described above, with clotting restored using 5 equivalents of antidote relative to the supramolecular inhibitor (Fig. 4d-e, **A8-E1 + AD2**). Based on these promising *in vitro* data, we assessed the ability of our designed antidote to reverse anticoagulation in the *in vivo* thrombosis model. In this experiment we first treated with 5mg/kg of our supramolecular construct (that provided effective anticoagulation in the needle injury thrombosis model) followed by administration of 5 molar equivalents of the 12-mer PNA antidote (9.4 mg/kg). Following addition of the antidote, anticoagulation was effectively reversed as determined by the amount of fibrin deposition and thrombus volume compared to control injuries lacking the antidote (Fig. 4 f-g). These data support the potential of supramolecular inhibitors as bona fide therapeutic leads and lays the foundation for targeting a range of therapeutic targets with this approach in the future.

## Conclusion

In summary, we have designed highly potent direct bivalent thrombin inhibitors that display 800-fold gain in activity relative to individual fragments by leveraging the constitutional dynamic properties of supramolecular binary fragments. The supramolecular pairing was achieved with PNA allowing simple tuning of the dissociation kinetics of the supramolecular complex. An important point of difference of these molecules compared to classical inhibitors is that the dynamic equilibrium can be modulated by external factors, yielding a simple strategy for reversing inhibition. This feature is highly pertinent for direct thrombin inhibition due to the risk of adverse bleeding side effects in anticoagulation therapy. These known side effects has stimulated the development of a number of antidotes to clinically approved anticoagulant drugs^28^ that centre on the use of expensive monoclonal antibodies and cocktails of competitive coagulation proteins. Our designed supramolecular anticoagulants showed potent thrombin inhibition and anticoagulation activities *in vitro* that could be rapidly reversed using small PNA-based antidotes. Importantly, this potent anticoagulant activity with on-demand reversibility was also demonstrated in an *in vivo* thrombosis model thus providing a starting point for the future use of this new therapeutic modality for bona fide anticoagulant drug candidates.

Importantly, the strategy adopted here offers a general mechanism to turn therapeutic activity on or off rapidly and is therefore not limited to applications in thrombosis. For example, the supramolecular concept would be amenable to the emerging area of immunotherapy where an antidote to a CAR-T response is highly desirable, or to immunomodulators where reversal of action is important in case of severe infection. The fact that assembly can be encoded by different sequences of low cost PNA should make it possible to multiplex programmable supramolecular drug candidates in the future. The strategy requires the identification of two fragments that bind synergistically to a protein of interest. DNA-encoded libraries making use of dual-display are poised to deliver such fragments for new targets lacking prior information.^29^ The same approach can also be considered with Fab fragments of antibodies.^30^

## Acknowledgements

This work was supported in part by the Swiss National Science Foundation (188406), NCCR Chemical Biology (185898), the Portuguese national funds via FCT - Fundação para a Ciência e a Tecnologia through project PTDC/BIA-BQM/2494/2020. J. R.-R. also acknowledges the support of RYC2021-033063-I funded by MCIN/AEI/10.13039/501100011033 and the European Union «NextGenerationEU»/PRTR.”

## Author contribution

M.D. conceived and performed the synthesis of all compounds reported and biochemical experiments. D.J.F., I.A., J.S.T.L and R.E.S. designed and performed *in vitro* clotting assays, Thrombinoscope assays and all animal experiments. S.P.J. supervised the animal experiments. S.A. conceived and synthesized preliminary compounds. J.R.R. and P.J.B.P. performed and analysed enzyme selectivity experiments. R.P. and N.W. conceived and supervised the work. The manuscript was written by M.D., D.J.F., R.P. and N.W.

## Extended Data

**Extended Data Figure 1.**
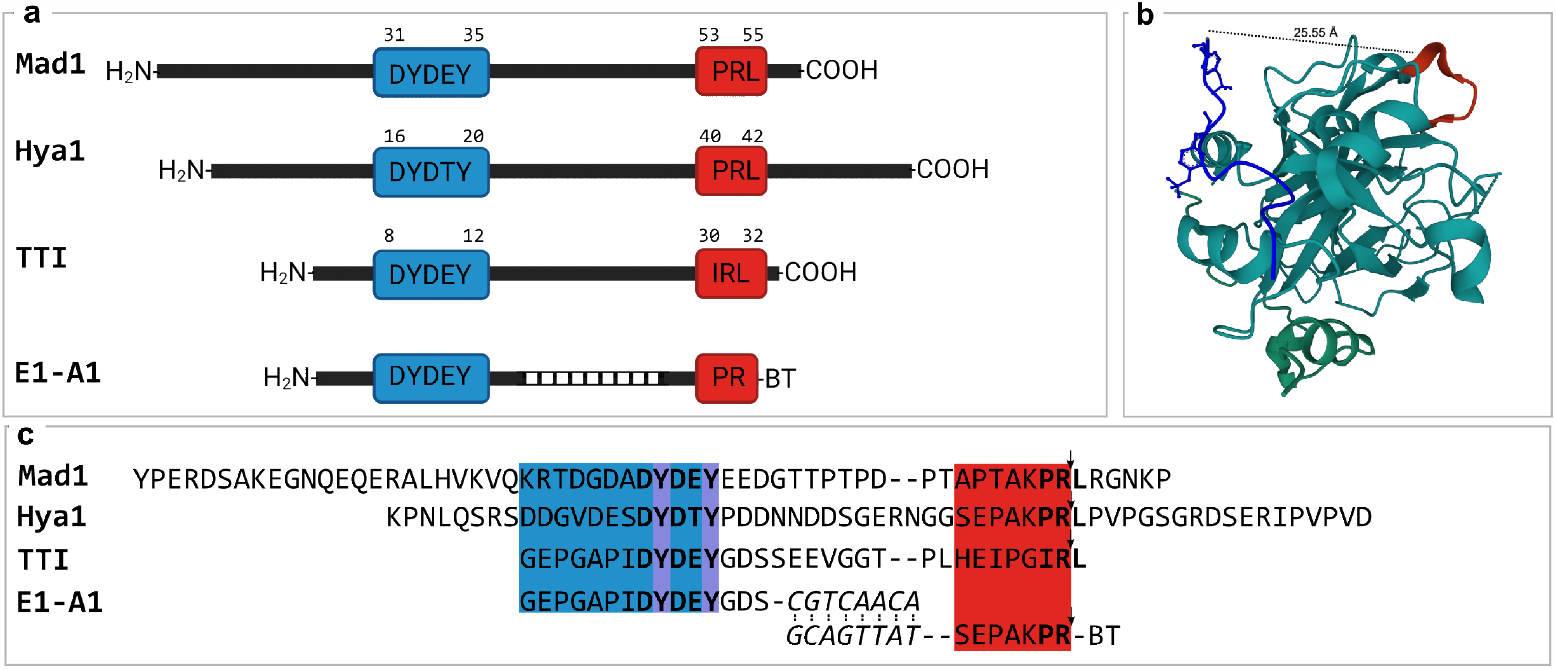
Sequence alignment of inhibitor from a range of blood-feeding insects alongside the supramolecular drug. **a**. Schematic representation of sequences of Madanin1, Hyalomin1 and Tsetse fly thrombin inhibitors alongside **A1-E1. b**. Crystal structure of the tsetse thrombin inhibitor (PDB 6TKG) showing the distance between the active site and exosite II binding components. **c**. Full sequences aligned. The exosite II sequence alignment is shown in blue with the sulphated tyrosine residues highlighted in purple. The active site sequence alignment is shown in red with the scissile bond represented by an arrow. The PNA is written in italics and the benzothiazole reversible covalent warhead is shorted to ‘BT’.

**Extended Data Figure 2.**
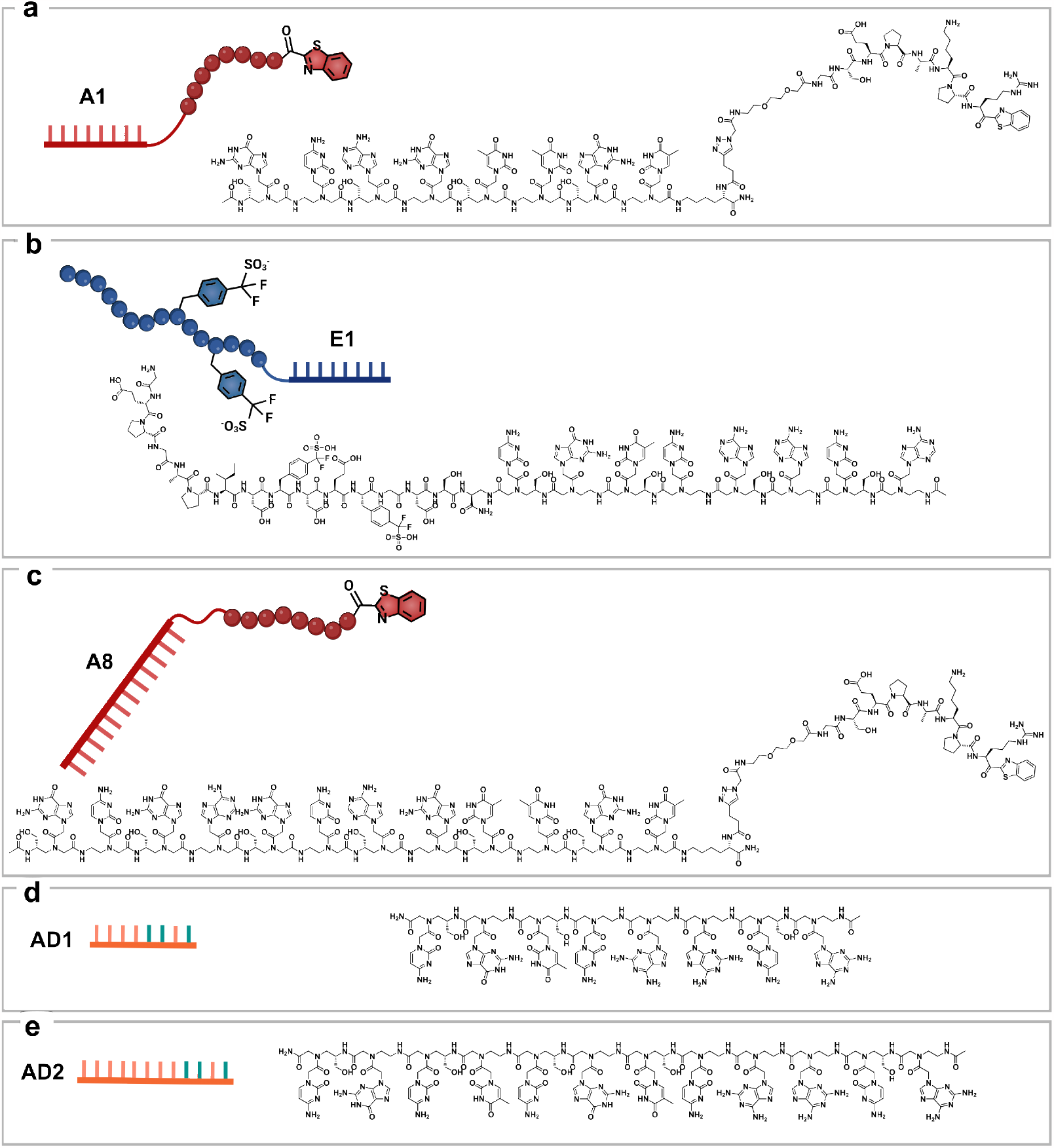
Chemical Structures of main compounds of the study. **a**. Active site-directed inhibitor **A1, b**. Exosite II-directed inhibitor **E1, c**. Active site-directed inhibitor with 4-mer toehold PNA **A8, d**. 8-mer antidote **AD1** and **e**. 12-mer antidote **AD2**.

**Extended Data Figure 3.**
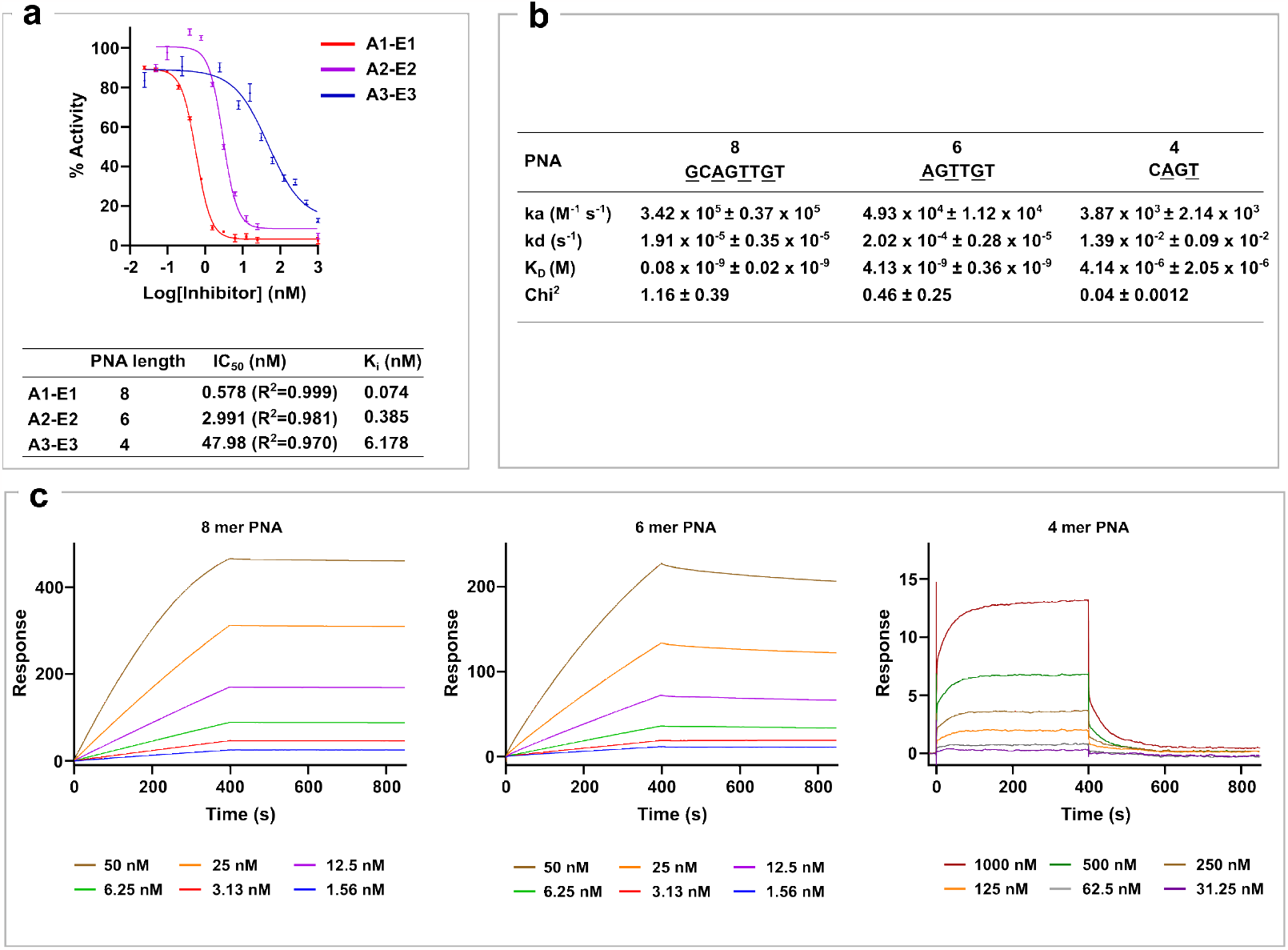
K_D_ of binding between PNAs of different lengths. **a**. Inhibition data for inhibitors with different length of PNA. **b**. SPR data for the binding between 4-, 6-, and 8-mer PNA strands with **A<u>C</u>A<u>A</u>C<u>T</u>G<u>C</u>** immobilised via a biotin on a streptavidin coated SPR chip. The PNA sequences are written N to C, with serine-modified monomers underlined. **c**. SPR kinetic curves.

**Extended Data Figure 4.**
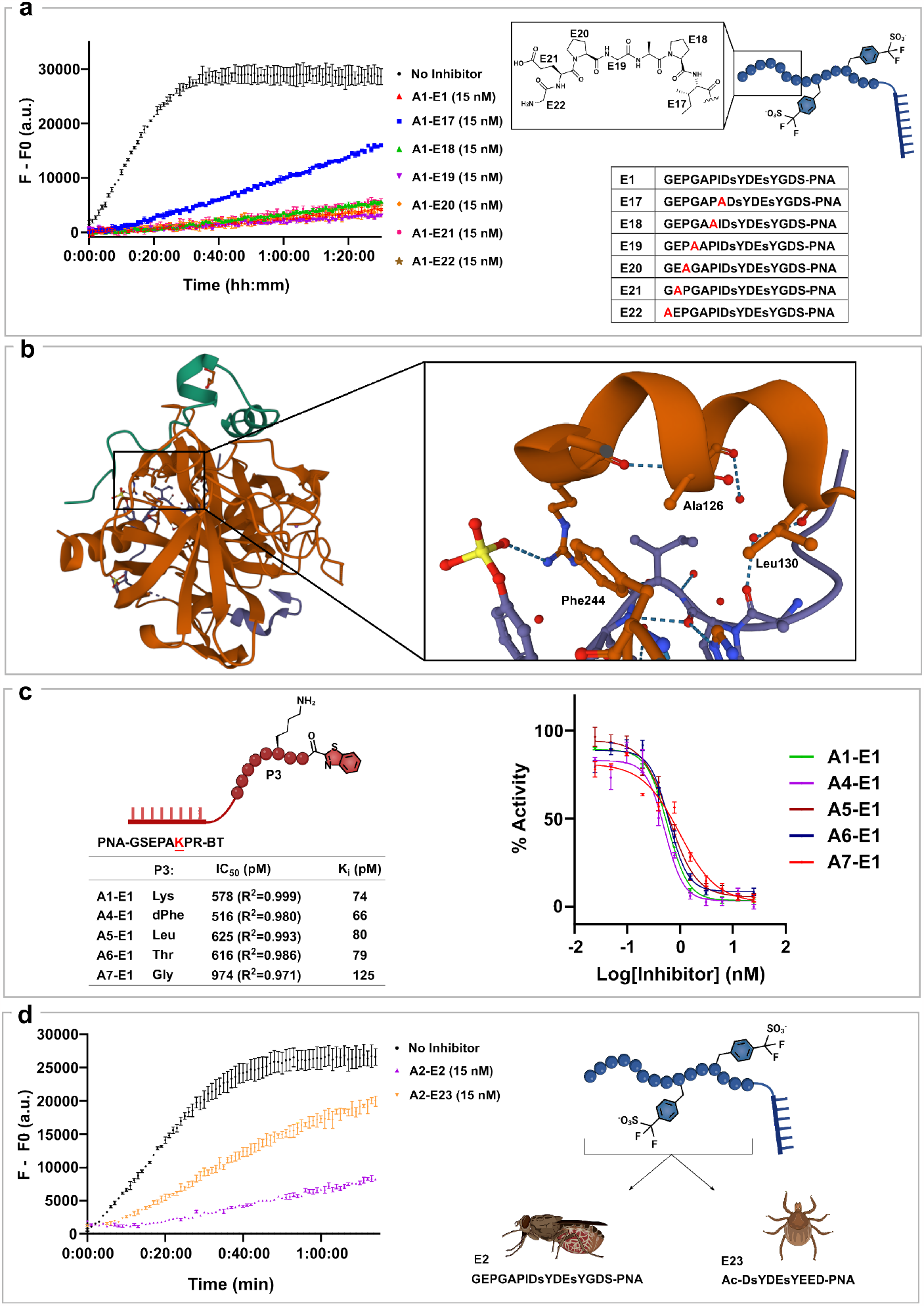
Additional SAR. **a**. Exosite II ala scan - fluorogenic inhibition assay data for **A1** combined with **E17** to **E22. b**. Crystal structure (PDB 6TKG) of the Tsetse thrombin inhibitor (TTI) complexed with thrombin. The zoom shows the hydrophobic pocket in which isoleucine is situated. **c**. IC_50_ values for active site binders with modifications in the P3 positions. **d**. Comparison between inhibitors inspired from the exosite II sequence of the Tsetse Thrombin inhibitor (TTI) and Madanin1 (Mad1, from the *Haemaphysalis longicornis species)* at 15 nM.

**Extended Data Figure 5.**
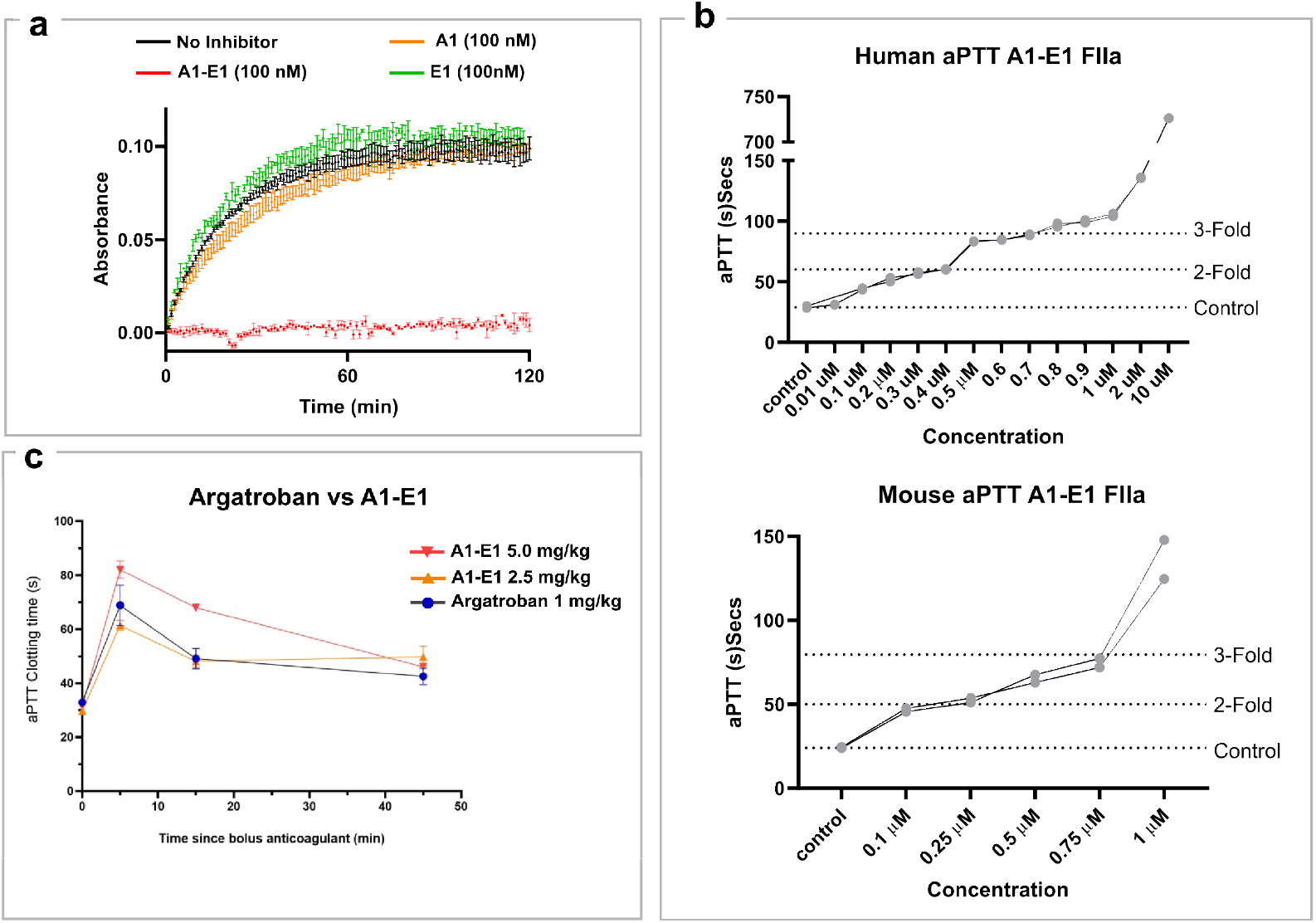
Additional Assays. **a**. Fibrinogen inhibition assay of compounds **A1-E1** and **A1** and **E1** alone. **b**. *In vitro* aPTT of **A1-E1** in human (top) and mouse (bottom) plasma. **c**. *Ex-vivo* aPTT of **A1-E1** at 0.314 μmol/kg (2.5 mg/kg) and 0.627 μmol/kg (5 mg/kg) versus Argatroban at 1.966 μmol/kg (1 mg/kg).

**Extended Data Figure 6.**
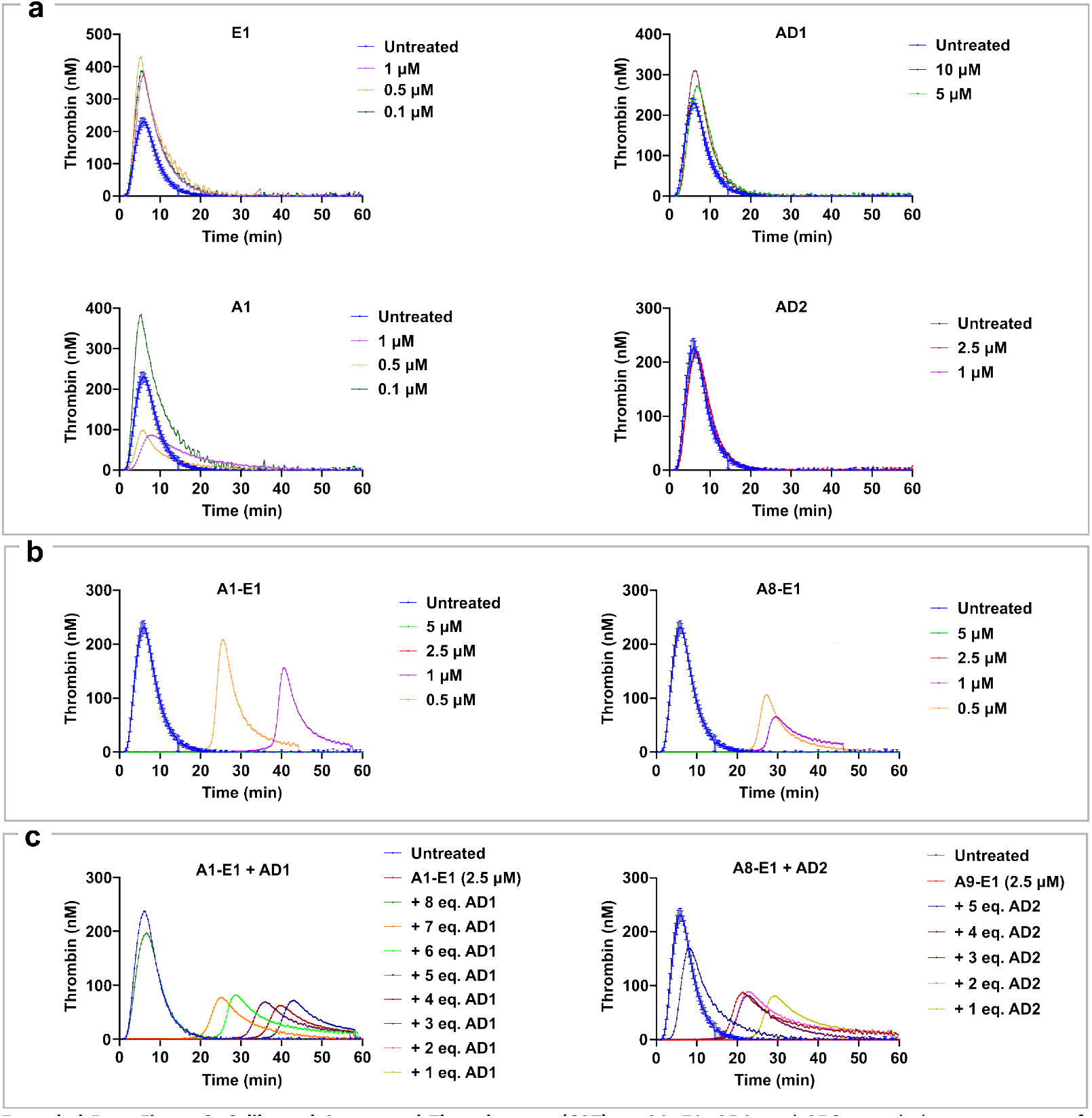
Calibrated Automated Thrombogram (CAT). **a. A1, E1, AD1**, and **AD2** tested alone at a range of concentrations. **b**. Combined inhibitors **A1-E1** and **A8-E1** tested at a range of concentrations. **c**. Combined inhibitors with antidote **A1-E1+AD1** and **A8-E1+AD2** tested at 2.5 μM with a range of antidote concentrations.

**Extended Data Figure 7.**
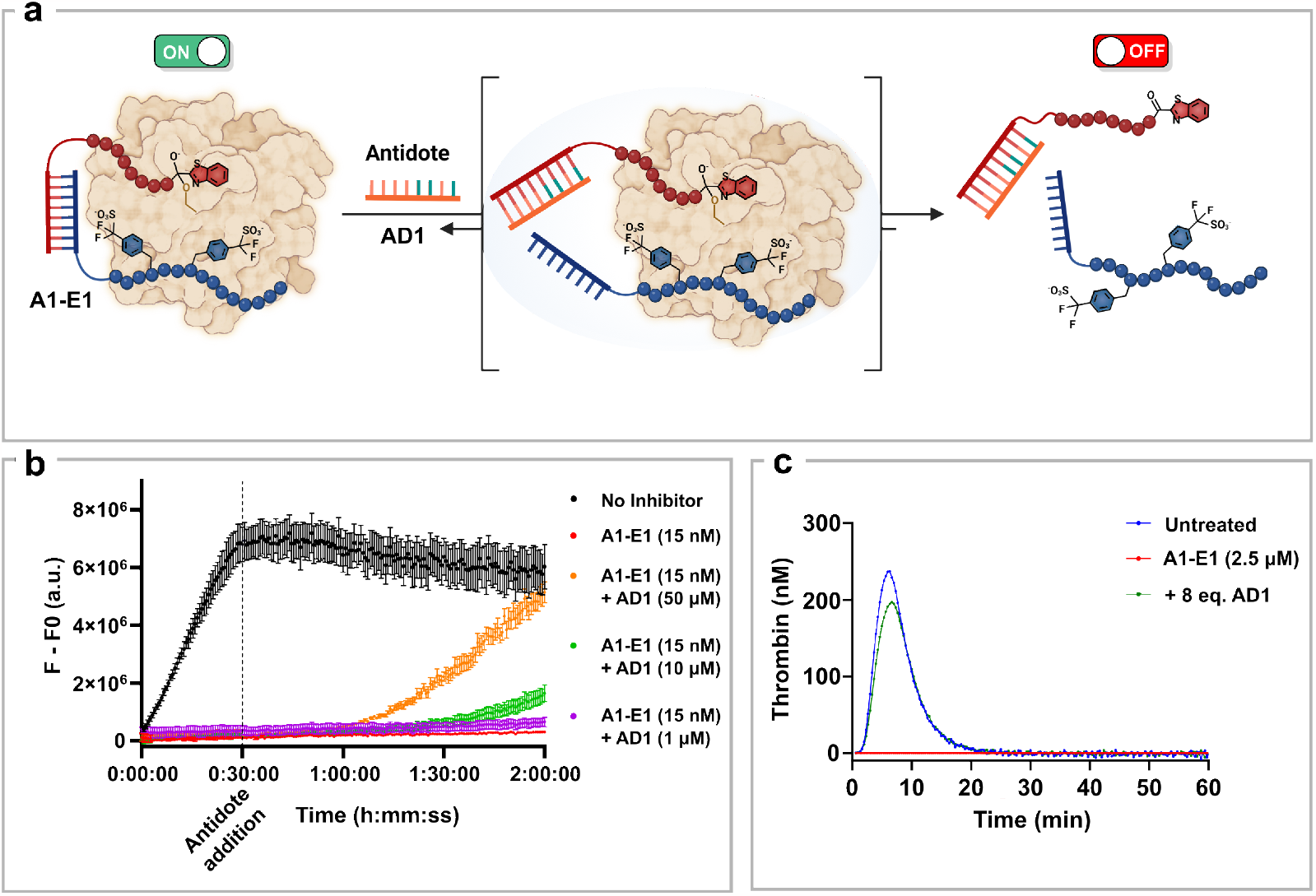
Reversal of thrombin inhibition with an 8 mer PNA antidote. **a**. Schematic representation of antidote addition and reversal of inhibition. **b**. Fluorogenic assay data showing the reversal of thrombin inhibition by addition of different concentrations of antidote (**AD1**) after 30 minutes of inhibition. **c**. Calibrated Automated Thrombogram (CAT) of **A1-E1** with and without antidote (**AD1**).

