## Supplementary Information for "Development of supramolecular anticoagulants with on-demand reversibility"

### 1. **Acronyms**

CuAAC	Copper(I)-catalyzed azide-alkyne cycloaddition

DAP	Diaminopropionic acid

DCE	1,2-dichloroethane

DCM	Dichloromethane

DHB	2,5-Dihydroxybenzoic acid

DIPEA	N,N-Diisopropylethylamine

DMF	Dimethylformamide

DMSO	Dimethyl sulfoxide

EDC	1-Ethyl-3-(3-dimethylaminopropyl)carbodiimide

ESI	Electrospray ionization

Fmoc	Fluorenylmethoxycarbonyl

HATU	Hexafluorophosphate Azabenzotriazole Tetramethyl Uronium

HIFP	Hexafluoroisopropanol

HOBt	Hydroxybenzotriazole

HPLC	High performance liquid chromatography

HRMS	High-resolution mass spectra

LCMS	Liquid chromatography-mass spectrometry

MALDI	Matrix-assisted laser desorption/ionization

MS	Mass spectrometry

NaAsc	Sodium ascorbate

NMP	N-Methyl-2-pyrrolidone

PEG	Polyethylene glycol

PNA	Peptide Nucleic Acid

TBTA	Tris((1-benzyl-4-triazolyl)methyl)amine

TFA	Trifluoroacetic acid

UHPLC	Ultra High performance liquid chromatography

### 2. **General methods**

All reagents and solvents for the organic synthesis were purchased from commercial sources and were used without further purification. HPLC purification was performed with an Agilent Technologies 1260 infinity HPLC using a ZORBAX 300SB-C18 column (9.4 x 250 mm). LC-MS spectra were recorded on a DIONEX Ultimate 3000 UHPLC with a Thermo LCQ Fleet Mass Spectrometer System using PINNACLE DB C18 column (1.9 µm, 50 x 2.1 mm) operated in positive mode. All the LC-MS spectra were measured by ESI. MALDI-TOF Mass spectra were measured using a Bruker Daltonics Autoflex spectrometer operated in positive mode. High-resolution mass spectra (HRMS) were obtained on a Xevo G2 Tof spectrometer (Ionization mode: ESI positive polarity; Mobile phase: MeOH 100 µl/min). Automated solid-phase synthesis was carried out on an Intavis AG Multipep RS instrument.

### 3. **Synthesis of compounds**

5.0 mg of resin were swollen in DCM for 10 minutes and washed twice with DMF. Iterative cycles of amide coupling (**Procedure 1**), capping of the resin (**Procedure 4**), and deprotection of the protecting group (**Procedure 2** **or 3**) were done to synthesize the PNA probes. The compounds were deprotected and cleaved from the resin using **Procedure 5** and finally purified using HPLC.

**2-Chlorotrityl Chloride Resin Loading**

2-Chlorotrityl chloride resin (1.46 mmol/g loading) was swollen in dry DCM for 30 min, followed by washing with DCM+1% DIPEA (1 x 3 mL) and DCM (10x 3 mL). A solution of Fmoc-Xaa-OH (0.7 mmol/g resin) and DIPEA (4 eq. relative to resin functionalization) in DCM (final concentration 0.125 M of amino acid) was added to the resin, which was shaken at room temperature for 16 h. The resin was then washed with DCM (5x 3 mL), DMF (5x 3 mL) and DCM (5x 3 mL). The resin was then capped via treatment with 17:2:1 v/v/v DCM:MeOH:DIPEA (5 mL) for 40 mins at room temperature. The resin was then washed again with DCM (5x 3 mL), DMF (5x 3 mL) and DCM (5x 3 mL) prior to further use.

**Rink Amide Resin Loading**

Nova PEG® Rink amide resin (0.44mmol/g, NovaBiochem) was swollen in DCM for 10 minutes and washed twice with DMF. Standard amide coupling (**Procedure 1**) was performed, followed by capping of the resin (**Procedure 4**). The resin was then washed again with DCM (5x 3 mL), DMF (5x 3 mL) and DCM (5x 3 mL) prior to further use.

**Procedure 1 (P1): Amide coupling.**

The corresponding Fmoc protected PNA monomer, or amino acid (4.0 equiv., 0.2M in NMP) was incubated for 5 minutes with HATU (3.5 equiv., 0.5M in NMP) and base solution [DIPEA, 1.2M (4.0 equiv.) and 2,6-lutidine 1.8M (6.0 equiv.) in NMP]. The mixture was then added to the corresponding resin. After 20 minutes, the mixture was filtered, the resin was washed with DMF, and a new premixed reaction solution was added to the resin and let react for another 20 minutes. Finally, the resin was washed with 2x DMF, 2x DCM, and 2x DMF.

**Procedure 2 (P2): Fmoc deprotection.**

A solution of 20% (v/v) piperidine in DMF was added to the resin and allowed to react for 5 minutes. The mixture was then filtered, the resin washed with DMF, and the sequence repeated for another 5 minutes. Finally, the resin was washed with 2x DMF, 2x DCM, and 2x DMF.

**Procedure 3 (P3): Mtt deprotection.**

A solution (made from 244 mg of HOBt in 10 mL HFIP and 10 mL DCE) was added to the prewashed resin to reach a volume of 10 mL/g of resin and allowed to react for 5 minutes. The solution was flushed, the resin washed with DCM_,_ and the sequence repeated for another 5 minutes. Finally, the resin was washed with 2x DCM and 2x DMF.

**Procedure 4 (P4): Capping.**

The resin was treated with a capping mixture (0.92 mL acetic anhydride and 1.3 mL 2,6 lutidine in 18 mL DMF: 10 mL of solution/g of resin) for 5 minutes. After flushing the solution, the resin was washed with 2x DMF, 2x DCM, and 2x DMF.

**Procedure 5 (P5): Cleavage from the resin and final deprotection.**

Resin (5.0 mg, 1.0 μmol) was treated with 125 μL of a mixture of TFA and scavengers (440 µL TFA + 25 mg phenol + 25 µL water + 10 µL triisopropylsilane) for 2 hours. The resin was filtered, washed with TFA (50 μL), and the collected fractions of cleavage product precipitated in cold ether (1.5 mL). After centrifugation, the pellet was vortexed again with cold Et_2_O (1.5 mL) and centrifuged (14k rpm). The pellet was dissolved in H_2_O/CH_3_CN (3:1, 1.5 mL) and lyophilized to obtain a white powder.

**Procedure 6 (P6): Microcleavage for quality control.**

The minimum number of beads were picked up with a pipette plastic tip and transferred to 50 µL of TFA. The solution was left for 1 hour and transferred to 1.0 mL of ether. The ether solution was kept for 5 minutes at -20 °C and then centrifuged for 5 minutes at 14k rpm. The ether supernatant was removed, and the pellet dissolved in 20 µL 1:1 acetonitrile/water, which was then used to analyze by MALDI and/or LC-MS.

**Procedure 7 (P7): On-resin Copper(I)-catalyzed azide-alkyne cycloaddition (CuAAC).**

To TBTA (2.0 mg) in 20 µL DMF was added 15 µL CuSO_4_ (64 mg/mL in H_2_O) followed by 50 µL of NaAsc (396 mg/mL in H_2_O). Azide-containing peptide (2 eq. in 60 µL DMF) was added to the mixture which was mixed prior to the addition to 5.0 mg alkyne derivatized rink amide resin (0.0022 mmol). After 16 hours of shaking, the mixture was filtered and the resin was washed with 6x 250 µL of sodium diethyl dithiocarbamate 0.02 M in DMF, 6x 250 µL of DMF, 6x MeOH and 6x DCM.

**Procedure 8 (P8): Coupling of Fmoc-l-F_2_Smp(nP)-OH**

Fmoc-l-F_2_Smp(nP)-OH was prepared following the procedure described by Dowman *et al*.^1^ A mixture of Fmoc-l-F_2_Smp(nP)-OH (0.003 mmol, 1.5 equiv.), HOBt (0.003 mmol, 1.5 equiv.) and DIC (0.003 mmol, 1.5 equiv.) were added to the corresponding resin and shaken overnight. The mixture was filtered and the resin was washed with 2x DMF, 2x DCM, and 2x DMF.

**Procedure 9 (P9): Coupling of Arg(Pbf)-Benzothiazole**

To 5.0 mg of resin (0.0022 mmol), Arg(Pbf)-Benzothiazole (0.0044 mmol, 2 equiv.) and HATU (0.0034 mmol, 1.5 equiv.) in NMP (100 μL) were added followed by DIPEA (0.012 mmol, 6 equiv.). The reaction was shaken for 2 hours. The mixture was filtered, and the resin was washed with 2x DMF, 2x DCM, and 2x DMF.

**Procedure 10 (P10): Neopentyl deprotection**

The precipitate collected after cleavage and ether precipitation was lyophilized. The remaining solid was dissolved in a solution 1M NH_4_Ac and 6M GnHCl and shaken at 37 °C for 2 hours. The solution was then diluted with H_2_O/ACN (50:50) and purified by HPLC.

**Characterization of PNA-peptide conjugates.**

Characterization of the PNA-peptide conjugates was done by MALDI (Bruker Daltonics Autoflex spectrometer with Flex control 3.4 software and analysis with FlexAnalysis 3.4) and/or LC-MS (DIONEX Ultimate 3000 UHPLC with a Thermo LCQ Fleet Mass Spectrometer System using PINNACLE DB C18 column (1.9 µm, 50 x 2.1 mm) with Thermo Xcalibur 2.2.SP1.48 software and analysis with Thermo Xcalibur Qual Browser 2.2.Sp1.48). For MALDI analysis, 1.0 µL of the sample (in either water or water/acetonitrile 1:1) was mixed with 1.0 µL of DHB matrix solution (30 mg of DHB in 1.0 mL of 70:30:0.01 water/acetonitrile/TFA), and the mixture spotted on a MALDI plate. The measurements were done in a positive linear mode. For LC-MS analysis, 20 µL of sample in water or water/acetonitrile 1:1 was injected on the LC and further analyzed by MS on a positive mode. Compounds containing the benzene disulfonic acid motif could only be analyzed by LC-MS due to a fragmentation when analyzed by MALDI.

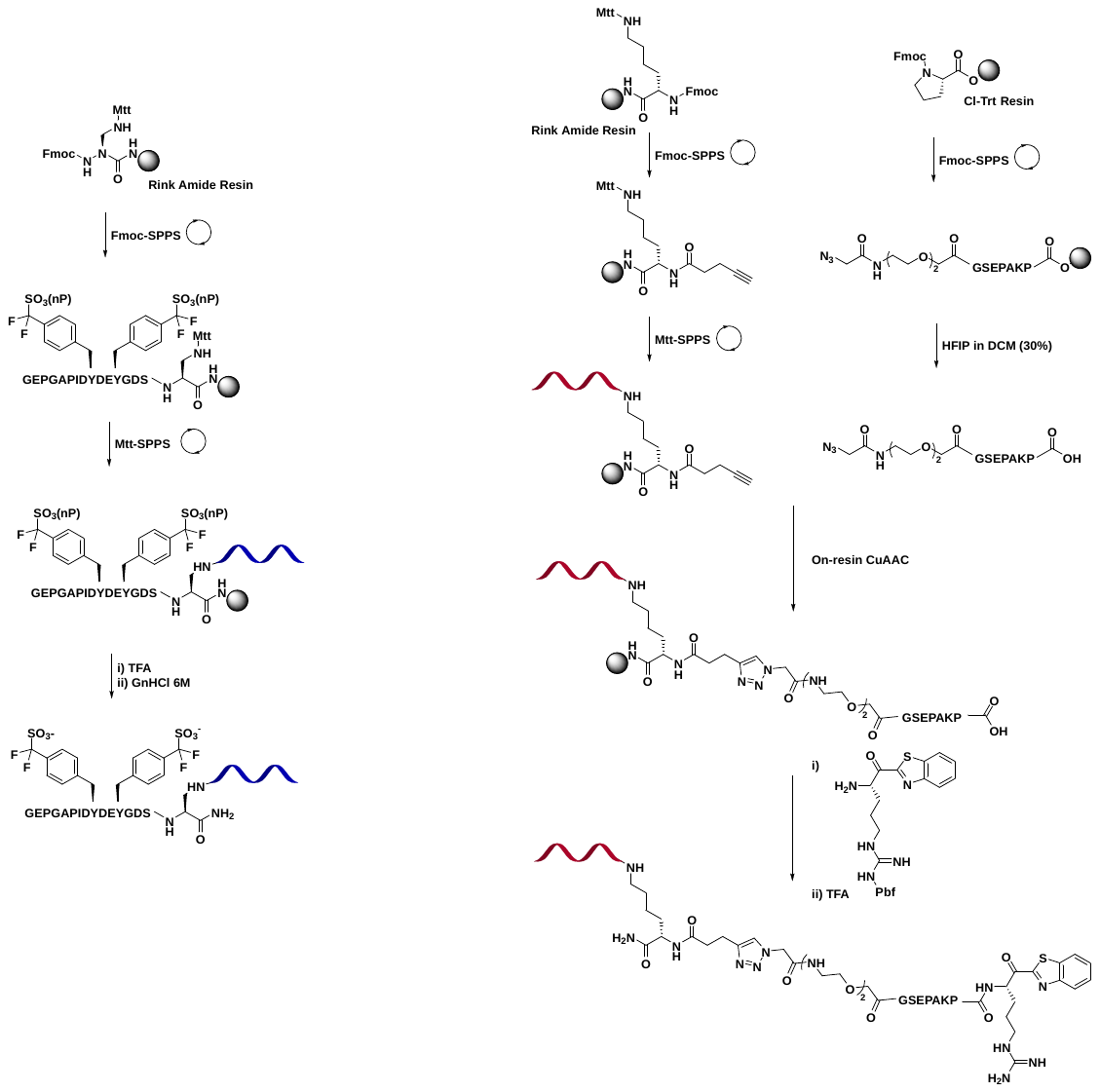
**General Scheme for the synthesis of the compounds**

**Tsetse thrombin inhibitor (TTI) exosite II binder (E1)**

**Hyalomin 1 (Hya1) active site binder (A1)**

### 4. **Synthesis of Arg(Pbf)-Benzothiazole**

**Tert-butyl (S,Z)-(1-(methoxy(methyl)amino)-1-oxo-5-(2-((2,2,4,6,7-pentamethyl-2,3-dihydrobenzofuran-5-yl)sulfonyl)guanidino)pentan-2-yl)carbamate**

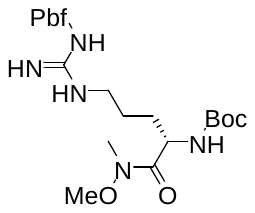

**Chemical Formula**: C_26_H_43_N_5_O_7_S, **Exact Mass**: 569.29, **Molecular Weight**: 569.72.

Boc-Arg(Pbf)-OH was prepared as previously described by Jakobsche *et al*.^2^ Boc-Arg(Pbf)-OH (1 g, 1.90 mmol) was dissolved in dry THF under N_2_. HATU (2.28 mmol, 1.2 equiv.) and DIPEA (9.5 mmol, 5 equiv.) were added, followed by MeNHOMe (2.28 mmol, 1.2 equiv.). The mixture was stirred at room temperature for 2.5 hours. The mixture was concentrated under *vacuo*, water was added and the product was extracted with EtOAc (3x), the organic layers were washed with H_2_O (2x), Brine (1x), dried over Na_2_SO_4_, and concentrated. The crude material was purified by flash chromatography (20% pentane in ethyl acetate) to yield the Weinreb amide (953 mg, 1.67 mmol, 88%).

**LCMS (ESI)**; RT= 2.63, [M+1H]^1+^: 570.05.

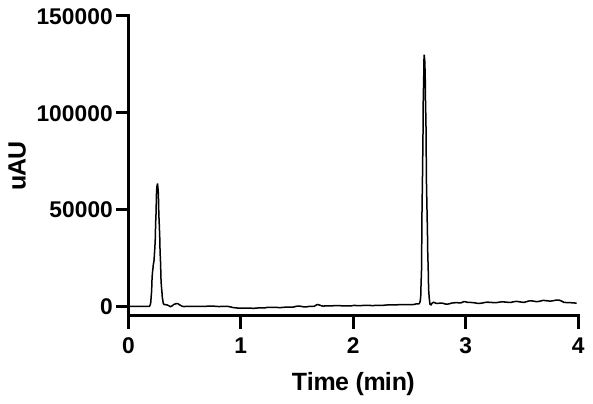

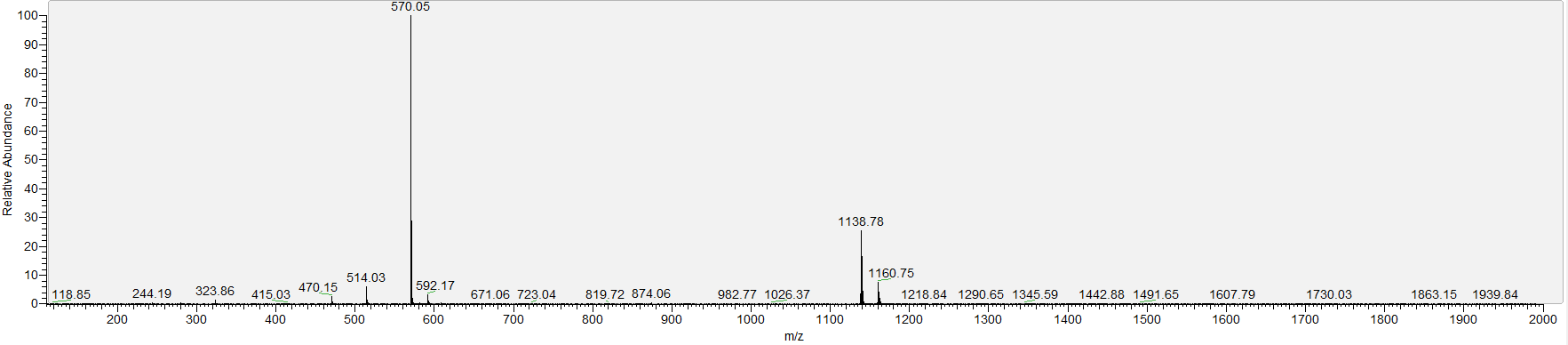

**NMR**; ^1^H NMR (400 MHz, CDCl_3_) δ 6.51 (s, 2H), 5.50 (d, *J* = 8.8 Hz, 1H), 4.63 (s, 1H), 3.74 (s, 3H), 3.42 (d, *J* = 7.6 Hz, 1H), 3.20 (s, 3H), 2.96 (s, 2H), 2.80 (s, 7H), 2.55 (s, 3H), 2.50 (s, 3H), 2.09 (s, 3H), 1.78 – 1.52 (m, 4H), 1.47 (s, 6H), 1.42 (s, 9H).

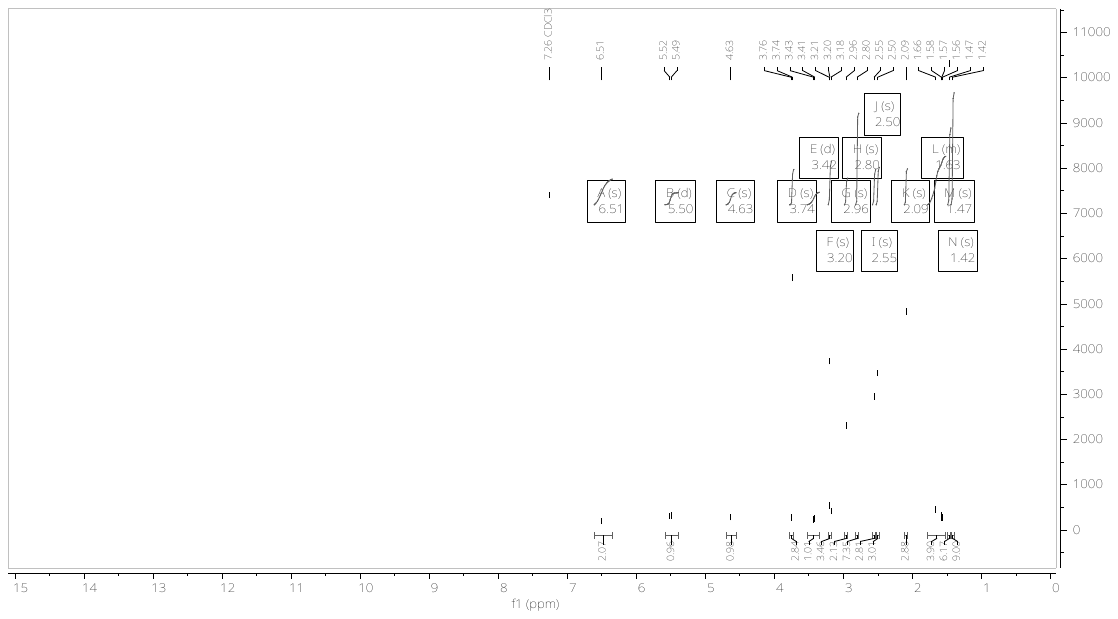

**tert-butyl (S)-(1-(benzo[d]thiazol-2-yl)-1-oxo-5-(3-((2,2,4,6,7-pentamethyl-2,3-dihydrobenzofuran-5-yl)sulfonyl)guanidino)pentan-2-yl)carbamate**

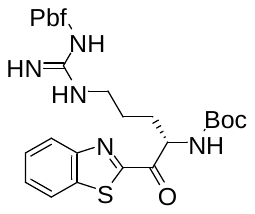

**Chemical Formula**: C_31_H_41_N_5_O_6_S_2_, **Exact Mass**: 643.25, **Molecular Weight**: 643.82

Benzothiazole (6.3 mmol, 18 equiv.) was added to dry THF (21 mL) under N_2_ and cooled to -78 °C. n-BuLi (6.125 mmol, 17.5 equiv.) was added dropwise over 10 minutes and stirred for 30 minutes at -78 °C. A solution of Weinreb amide (0.35 mmol) in THF (11 mL) was added dropwise at -78 °C and stirred for 2 hours. The reaction was quenched with 10 mL of saturated NH_4_Cl and warmed to room temperature. The mixture was extracted with EtOAc (3x), the organic layers were washed with H_2_O (2x), Brine (1x) and dried over Na_2_SO_4_ and concentrated. The crude product was purified by flash chromatography (10 to 100% EtOAc in Pentane) to yield Boc-Arg(Pbf)-Benzothiazole as brown solid (170 mg, 0.26 mmol, 75%).

**LCMS (ESI)**; RT= 3.13, [M+1H]^1+^: 644.06

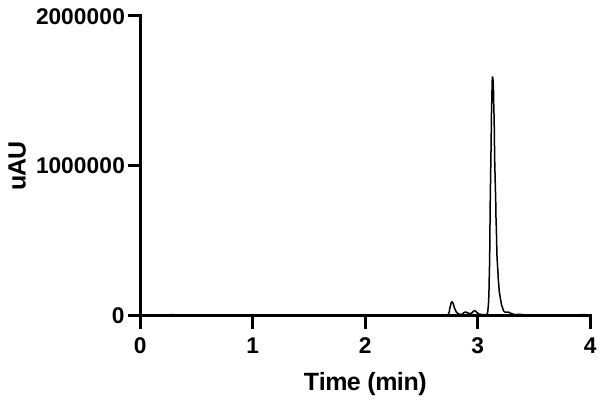

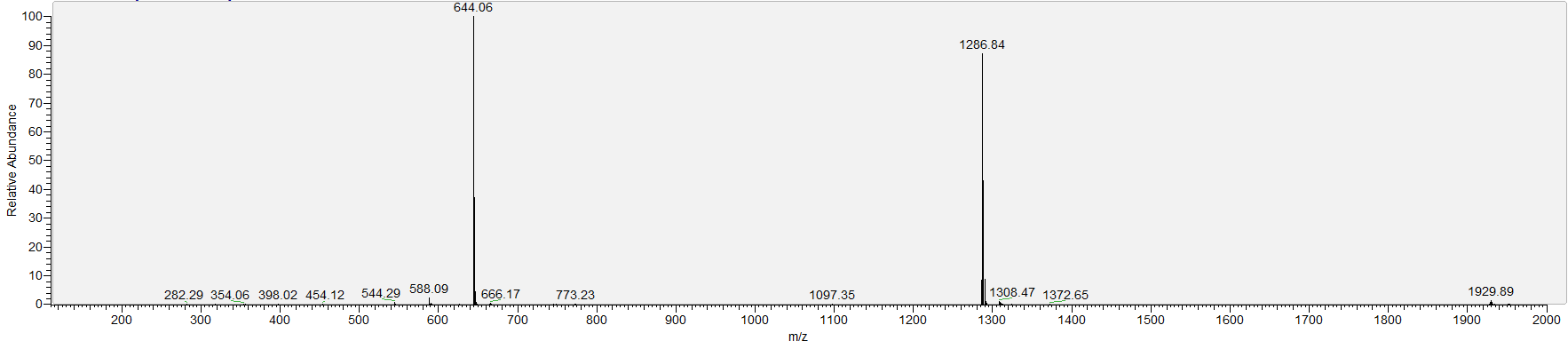

**NMR;** ^1^H NMR (400 MHz, CDCl_3_) δ 8.25 – 8.15 (m, 1H), 8.05 – 7.89 (m, 1H), 7.66 – 7.49 (m, 2H), 6.38 (s, 1H), 6.26 (s, 2H), 5.74 – 5.51 (m, 2H), 3.32 (s, 2H), 2.95 (s, 2H), 2.56 (s, 3H), 2.51 (s, 3H), 2.08 (s, 3H), 1.70 – 1.66 (m, 4H), 1.45 (d, *J* = 9.5 Hz, 15H).

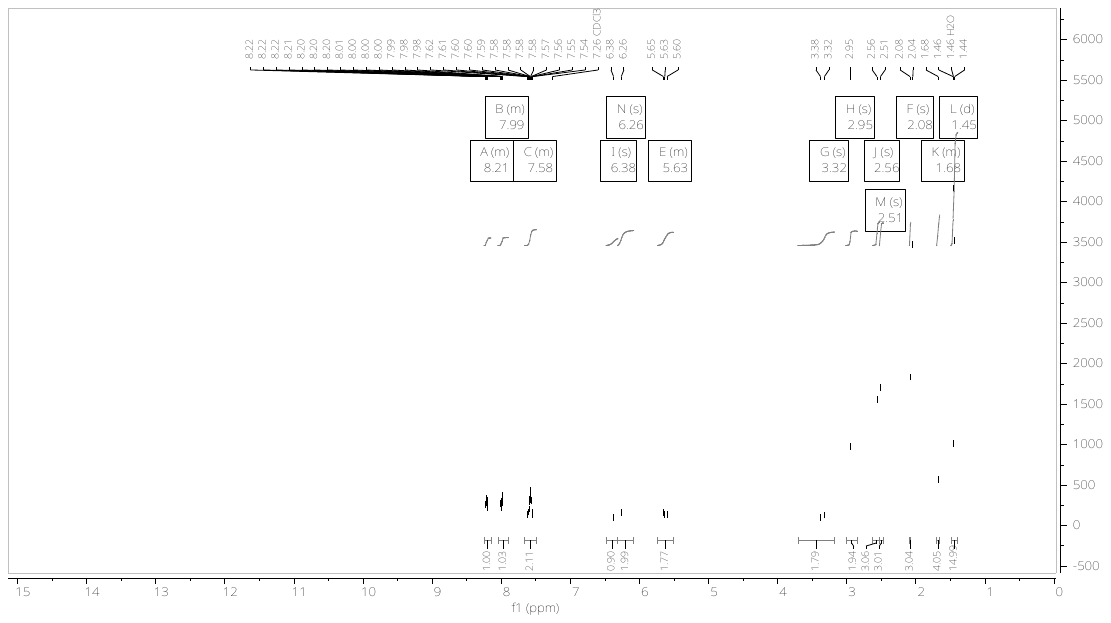

**(S)-N-(N-(4-amino-5-(benzo[d]thiazol-2-yl)-5-oxopentyl)carbamimidoyl)-2,2,4,6,7-pentamethyl-2,3-dihydrobenzofuran-5-sulfonamide**

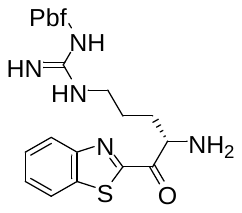

**Chemical Formula**: C_26_H_33_N_5_O_4_S_2_, **Exact Mass**: 543.20, **Molecular Weight**: 543.70.

Boc-Arg(Pbf)-Benzothiazole (74 mg, 0.115 mmol) was dissolved in dioxane and cooled to 0 °C. 4M HCl in dioxane was added and the mixture was stirred for 2 hours at room temperature. The mixture was concentrated under *vacuo* to yield Arg(Pbf)-Benzothiazole as a brown solid (63 mg, quant.). The compound was used without further purification.

**LCMS (ESI)**; RT= 2.22, [M+1H]^1+^: 544.18.

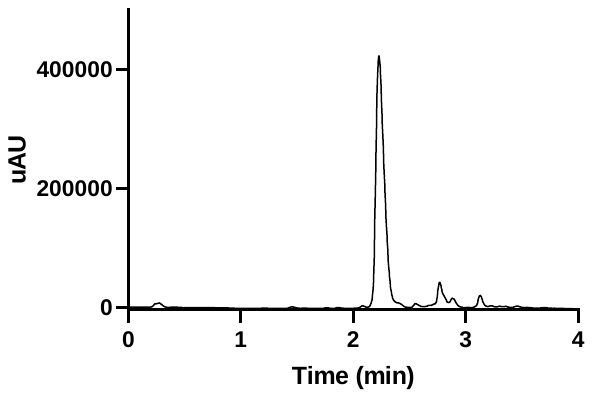

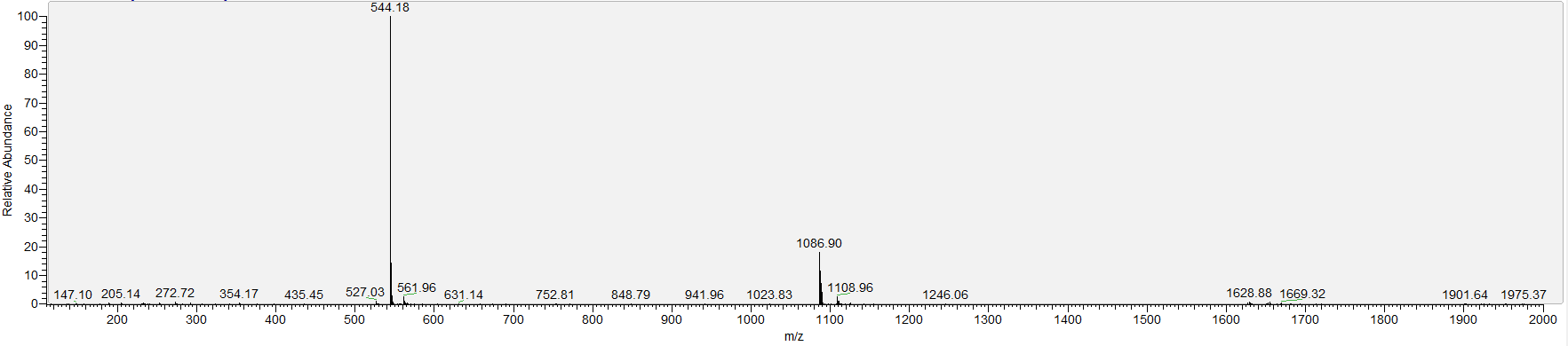

**NMR;** ^1^H NMR (400 MHz, CDCl_3_) δ 8.89 (s, 2H), 8.39 (s, 1H), 8.22 – 8.13 (m, 1H), 7.83 (s, 1H), 7.49 (s, 2H), 5.52 (s, 1H), 3.74 (d, *J* = 7.5 Hz, 1H), 3.62 – 3.35 (m, 2H), 2.90 (s, 2H), 2.54 (d, *J* = 6.3 Hz, 2H), 2.41 (d, *J* = 19.4 Hz, 6H), 2.19 (s, 2H), 1.43 (s, 6H).

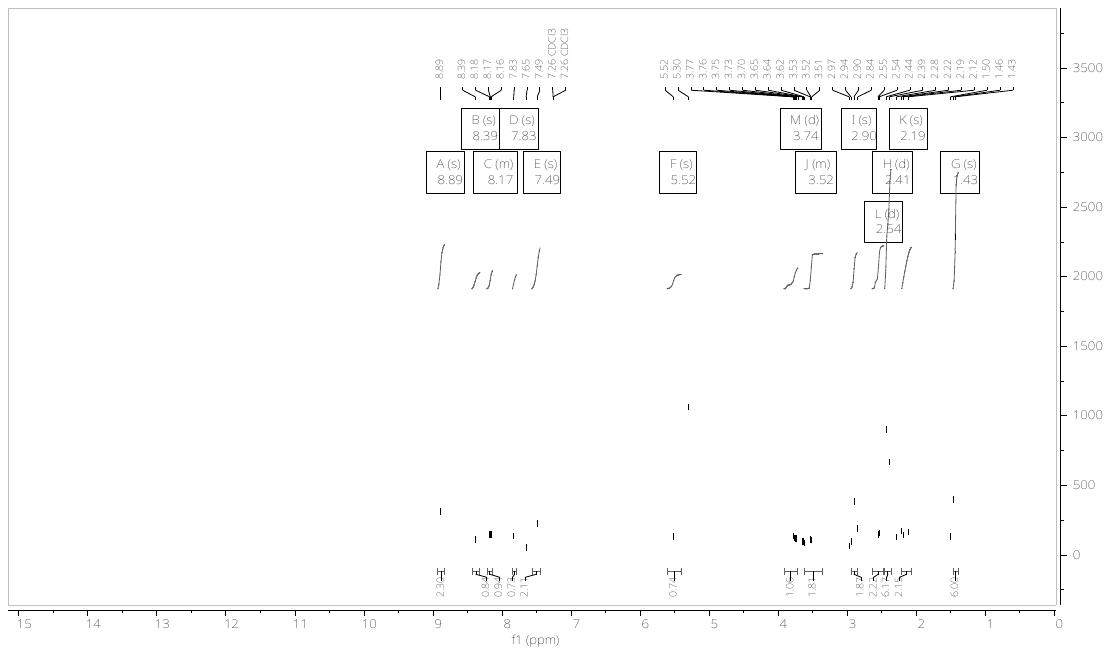

### 5. **Synthesis of phe-Pro-Arg-Coumarin**

**(9H-fluoren-9-yl)methyl (S)-(1-((4-methyl-2-oxo-2H-chromen-7-yl)amino)-1-oxo-5-(3-((2,2,4,6,7-pentamethyl-2,3-dihydrobenzofuran-5-yl)sulfonyl)guanidino)pentan-2-yl)carbamate (Fmoc-Arg(Pbf)-Coumarin)**

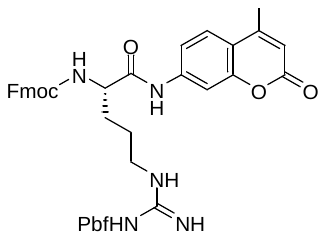

**Chemical Formula**: C_44_H_47_N_5_O_8_S, **Exact Mass**: 805.32, **Molecular Weight**: 805.95.

7-amino-4-methylcoumarin (0.28 mmol) and Fmoc-Arg(Pbf)-OH (0.28 mmol) were dissolved in pyridine (1.6 mL) and cooled to 0 °C. POCl_3_ (0.052 mL) was added dropwise, the reaction was allowed to warm to room temperature and stirred for 1 hour. The reaction mixture was quenched with 10 mL H_2_O and extracted with EtOAc (3x). The organic layers were washed with 2 M HCl (1x), 5% NaHCO_3_ (1x) and brine (1x), dried on sodium sulfate and concentrated under *vacuo*. The crude product was purified by reverse phase chromatography (Biotage, 48g C18 column, 10-100% ACN in H_2_O) to yield Fmoc-Arg(Pbf)-Coumarin (0.112 mmol, 40%).

**LCMS (ESI)**; RT= 3.03, [M+1H]^1+^: 806.13.

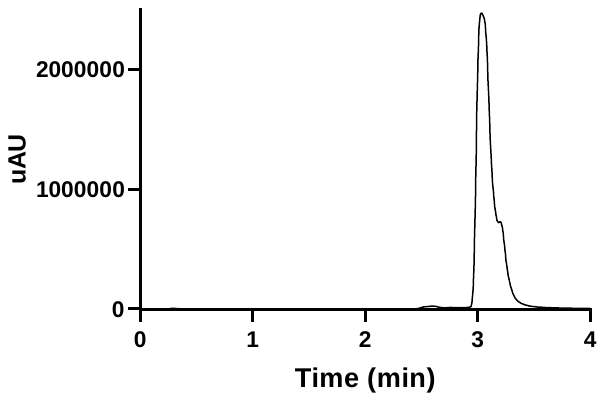

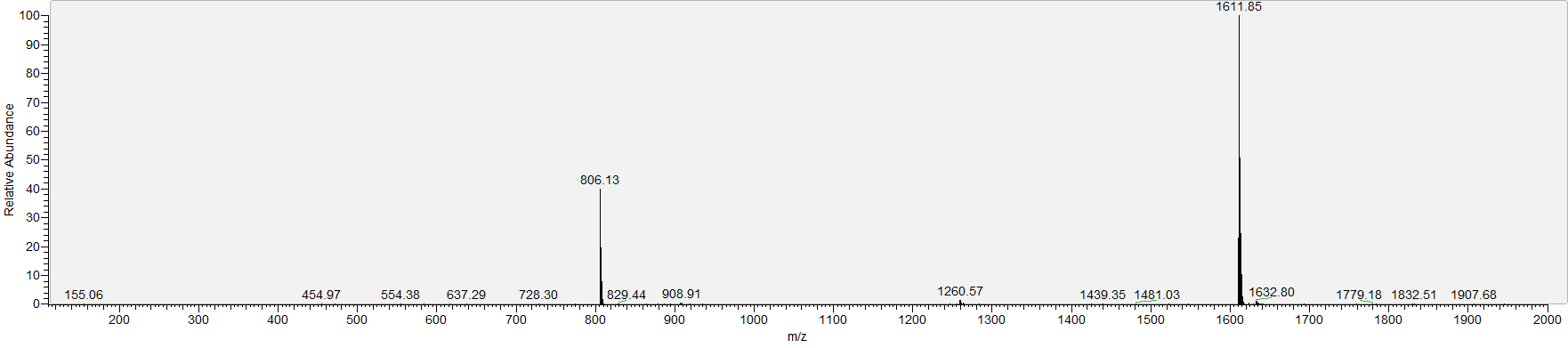

**NMR;** ^1^H NMR (400 MHz, DMSO) δ 10.49 (s, 1H), 7.88 (d, *J* = 7.5 Hz, 2H), 7.80 – 7.69 (m, 5H), 7.49 (dd, *J* = 8.7, 2.1 Hz, 1H), 7.40 (t, *J* = 7.5 Hz, 2H), 7.31 (tt, *J* = 7.4, 1.6 Hz, 2H), 6.27 (d, *J* = 1.4 Hz, 1H), 4.28 (d, *J* = 8.0 Hz, 2H), 4.22 (d, *J* = 6.8 Hz, 1H), 4.15 (td, *J* = 8.4, 5.6 Hz, 1H), 3.06 (h, *J* = 6.5 Hz, 2H), 2.89 (s, 2H), 2.45 (s, 3H), 2.41 – 2.37 (m, 6H), 1.95 (s, 3H), 1.76 – 1.56 (m, 2H), 1.50 (t, *J* = 7.6 Hz, 2H), 1.37 (s, 6H).

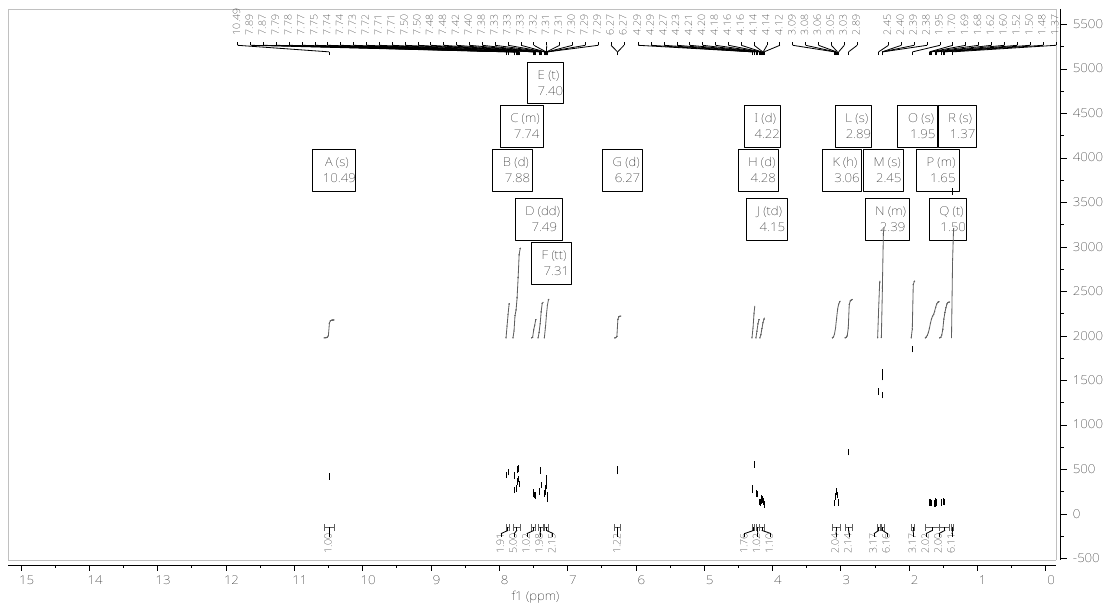

**(9H-fluoren-9-yl)methyl (S)-2-(((S)-1-((4-methyl-2-oxo-2H-chromen-7-yl)amino)-1-oxo-5-(3-((2,2,4,6,7-pentamethyl-2,3-dihydrobenzofuran-5-yl)sulfonyl)guanidino)pentan-2-yl)carbamoyl)pyrrolidine-1-carboxylate**

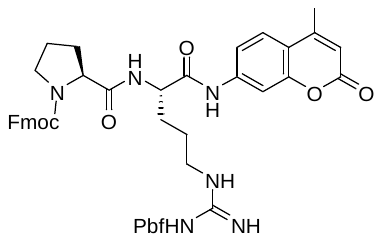

**Chemical Formula**: C_49_H_54_N_6_O_9_S, **Exact Mass**: 902.37, **Molecular Weight**: 903.06.

Fmoc-Arg(Pbf)-Coumarin (0.11 mmol) was treated with DEA (1 mL) in acetonitrile (1 mL) for 30 minutes. The reaction mixture was concentrated under *vacuo*, redissolved in acetonitrile (5 mL) and reconcentrated (2x). The crude Arg(pbf)-coumarin was dissolved in 0.3 mL dry DMF.

Fmoc-Pro-OH (0.12 mmol), HATU (0.11 mmol) and DIPEA (0.33 mmol) were dissolved in dry DMF (0.8 mL) and stirred at 0 °C for 15 minutes. Arg(pbf)-coumarin (0.11 mmol) in 0.3 mL dry DMF was added dropwise. The reaction mixture was allowed to warm to room temperature and stirred for 1 hour. The reaction mixture was quenched with 10 mL H_2_O and extracted with EtOAc (3x). The organic layers were washed with 2 M HCl (1x), 5% NaHCO_3_ (1x) and brine (1x), dried on sodium sulfate and concentrated under *vacuo*. The crude product was purified by reverse phase chromatography (Biotage, 48g C18 column, 10-100% ACN in H_2_O) to yield Fmoc-Pro-Arg(Pbf)-Coumarin (0.070 mmol, 64%).

**LCMS (ESI)**; RT= 2.89, [M+1H]^1+^: 903.17.

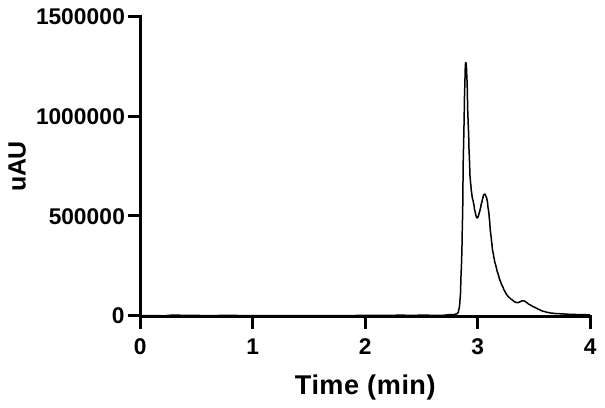

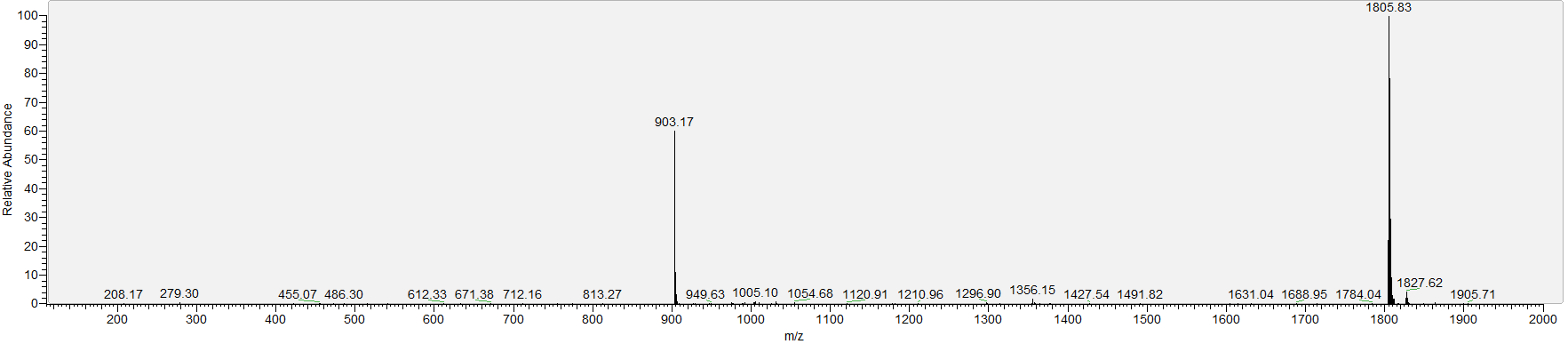

**NMR;** ^1^H NMR (400 MHz, DMSO) δ 7.91 – 7.88 (m, 1H), 7.82 – 7.77 (m, 1H), 7.73 – 7.61 (m, 3H), 7.51 (ddd, *J* = 8.8, 5.4, 3.3 Hz, 1H), 7.44 – 7.26 (m, 5H), 6.29 – 6.22 (m, 1H), 4.47 (dt, *J* = 9.0, 4.5 Hz, 1H), 4.41 – 4.30 (m, 1H), 4.30 – 4.10 (m, 3H), 3.47 (s, 3H), 3.06 (s, 2H), 2.87 (d, *J* = 12.4 Hz, 2H), 2.44 (s, 2H), 2.41 – 2.37 (m, 5H), 2.34 (s, 2H), 1.94 (d, *J* = 11.5 Hz, 3H), 1.88 – 1.67 (m, 4H), 1.54 (d, *J* = 62.5 Hz, 3H), 1.37 (s, 6H).

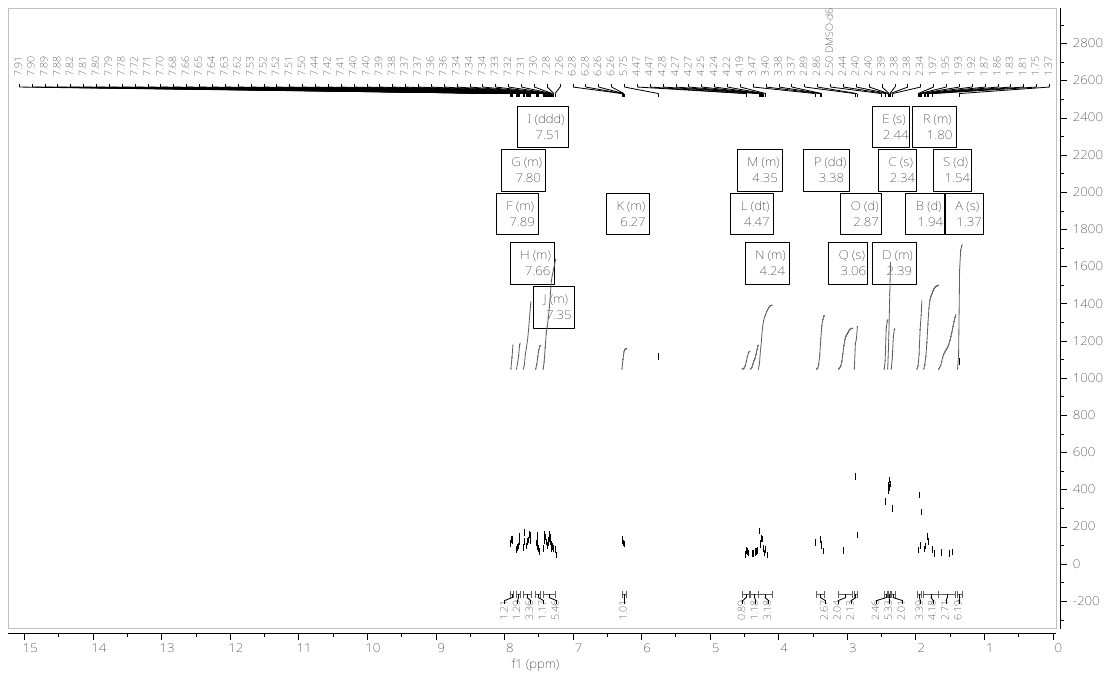

**(S)-1-(D-phenylalanyl)-N-((S)-5-guanidino-1-((4-methyl-2-oxo-2H-chromen-7-yl)amino)-1-oxopentan-2-yl)pyrrolidine-2-carboxamide (phe-Pro-Arg-Coumarin)**

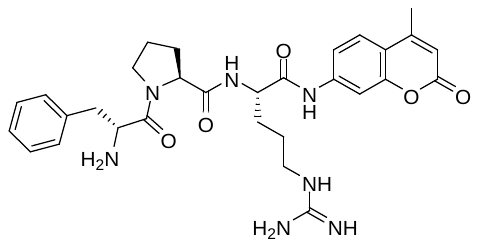

**Chemical Formula**: C_30_H_37_N_7_O_5_, **Exact Mass**: 575.29, **Molecular Weight**: 575.67.

Fmoc-Pro-Arg(Pbf)-Coumarin (0.066 mmol) was treated with DEA (1.5 mL) in acetonitrile (1.5 mL) for 30 minutes. The reaction mixture was concentrated under *vacuo*, redissolved in acetonitrile (5 mL) and reconcentrated (2x). The crude Pro-Arg(pbf)-coumarin was dissolved in 0.2 mL dry DMF.

Fmoc-d-phe-OH (0.073 mmol), HATU (0.066 mmol) and DIPEA (0.198 mmol) were dissolved in dry DMF (0.5 mL) and stirred at 0 °C for 15 minutes. Pro-Arg(pbf)-coumarin (0.066 mmol) in 0.2 mL dry DMF was added dropwise. The reaction mixture was allowed to warm to room temperature and stirred for 1 hour. The reaction mixture was quenched with 10 mL H_2_O and extracted with EtOAc (3x). The organic layers were washed with 2 M HCl (1x), 5% NaHCO_3_ (1x) and brine (1x), dried on sodium sulfate and concentrated under *vacuo*.

The crude protected peptide was dissolved in 50:50 TFA:DCM and stirred at room temperature for 2 hours. The mixture was concentrated under *vacuo* and azeotroped with toluene three times. The peptide was then dissolved in 20% piperidine in DMF and stirred for 30 minutes. The mixture was purified by reverse phase chromatography (Biotage, 48g C18 column, 10-100% ACN in H_2_O) to yield phe-Pro-Arg(Pbf)-Coumarin (0.015 mmol, 23%).

**LCMS (ESI)**; RT= 1.54, [M+1H]^1+^: 576.26.

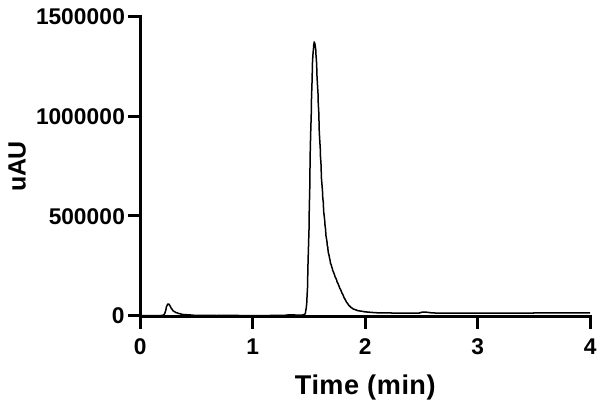

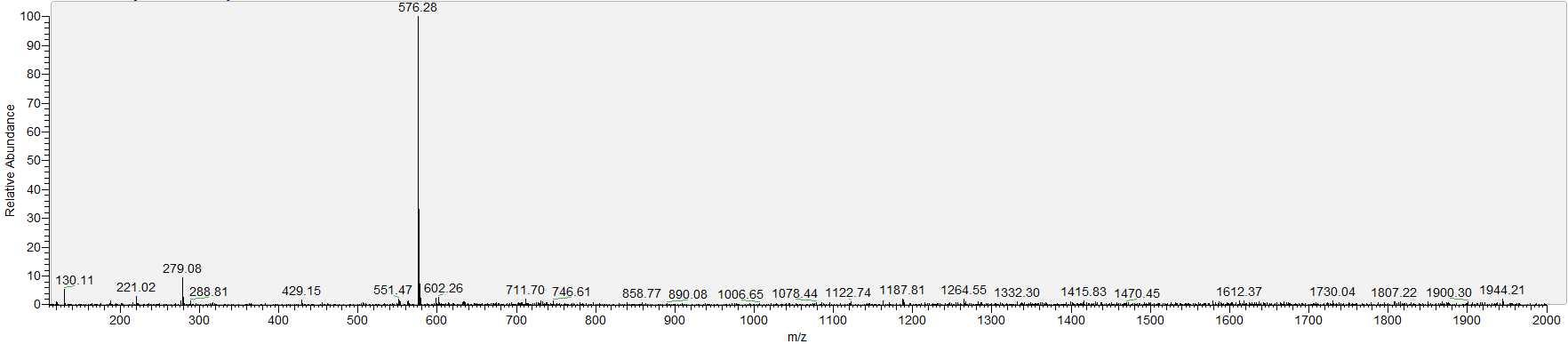

**NMR;** ^1^H NMR (400 MHz, DMSO) δ 10.41 (s, 1H), 8.37 – 8.29 (m, 1H), 8.24 (s, 2H), 7.79 – 7.77 (m, 1H), 7.74 – 7.70 (m, 1H), 7.58 – 7.52 (m, 1H), 7.50 – 7.45 (m, 1H), 7.41 – 7.28 (m, 4H), 7.26 – 7.22 (m, 2H), 6.29 (s, 1H), 4.49 – 4.23 (m, 3H), 3.58 – 3.46 (m, 2H), 3.19 – 3.04 (m, 3H), 3.02 – 2.92 (m, 1H), 2.85 – 2.69 (m, 1H), 2.41 (s, 3H), 1.85 – 1.70 (m, 4H), 1.69 – 1.56 (m, 2H), 1.55 – 1.42 (m, 2H).

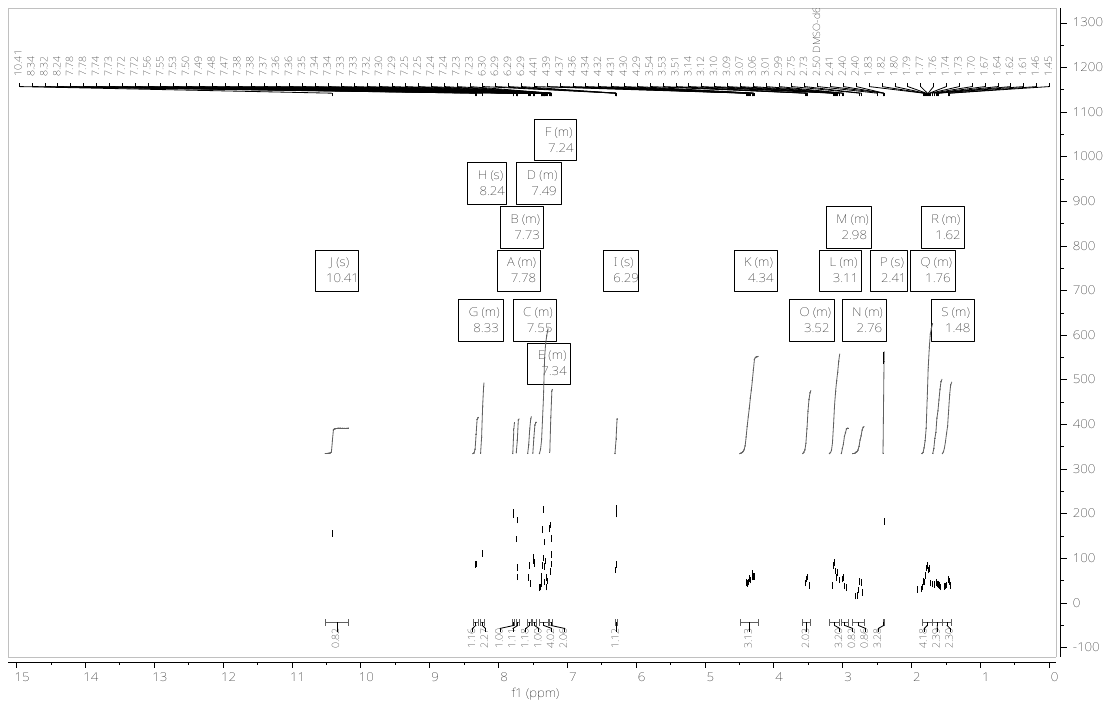

^13^C NMR (126 MHz, DMSO) δ 170.96, 170.77, 166.24, 159.68, 156.31, 153.32, 152.77, 141.67, 134.15, 129.22, 128.26, 127.17, 125.69, 114.98, 114.90, 112.13, 105.43, 59.07, 53.17, 51.61, 46.95, 46.51, 40.12, 39.78, 36.47, 28.99, 28.55, 24.94, 23.48, 17.67.

### 6. **Characterisation of PNA-peptide compounds**

i) **Inhibitors**

**A1**

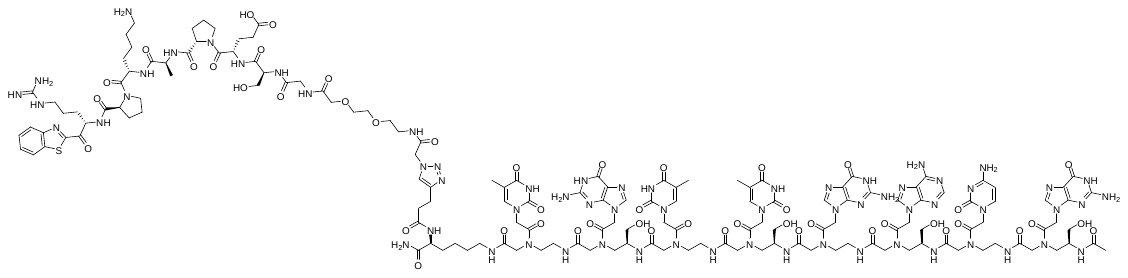

**Chemical Formula**: C_154_H_211_N_65_O_48_S, **Exact Mass**: 3770.58, **Molecular Weight**: 3772.85.

**MALDI-TOF**; m/z found: 3772.40.

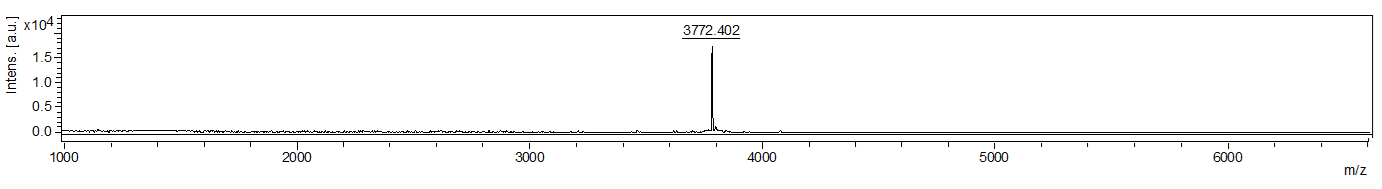

**LCMS (ESI)**; RT= 1.44 min, [M+3H]^3+^: 1258.42, [M+4H]^4+^: 944.25, [M+5H]^5+^: 755.50, [M+6H]^6+^: 629.92.

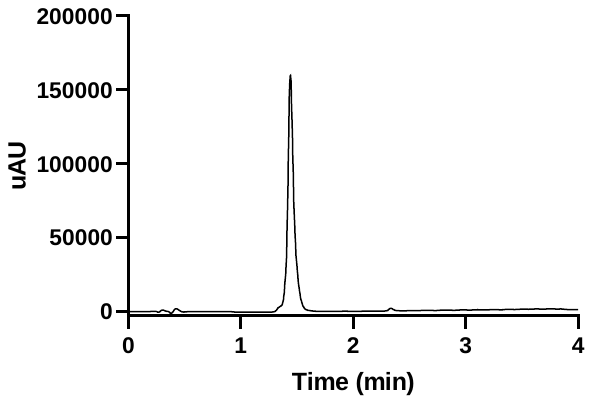

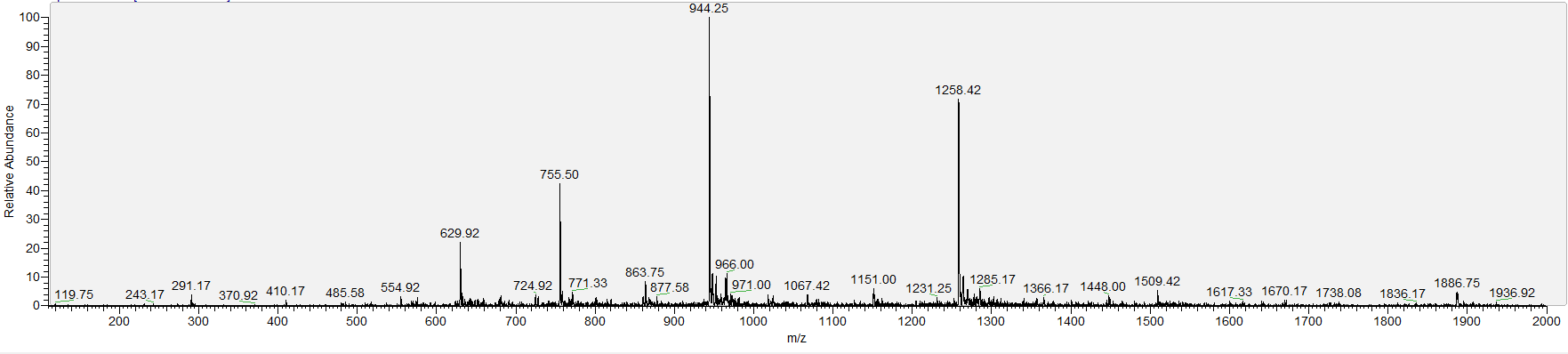

**HRMS;** [M+4H]^4+^: 943.8892, [M+5H]^5+^: 755.5194.

**E1**

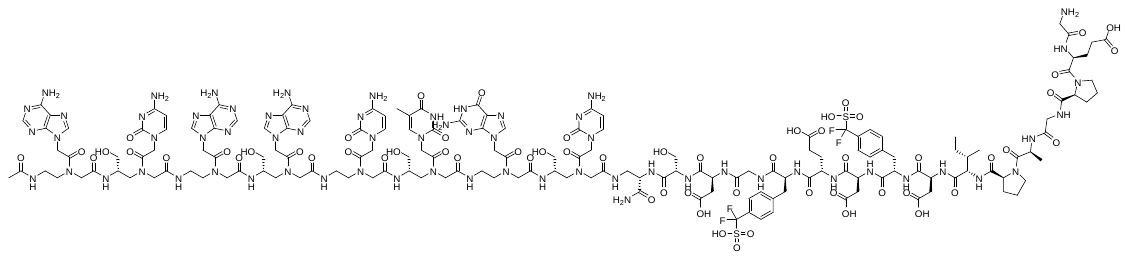

**Chemical Formula**: C_164_H_215_F_4_N_65_O_60_S_2,_ **Exact Mass**: 4194.52, **Molecular Weight**: 4197.03.

**MALDI-TOF**; m/z found: 4197.52.

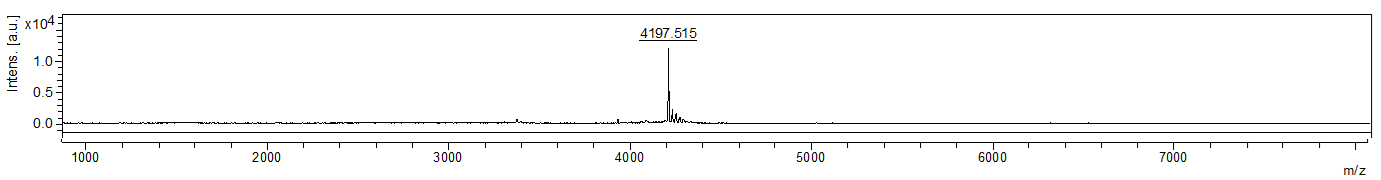

**LCMS (ESI)**; RT= 1.38 min, [M+3H]^3+^: 1399.75, [M+4H]^4+^: 1050.25.

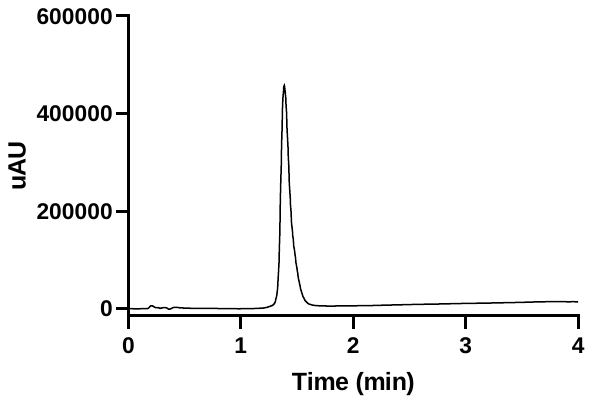

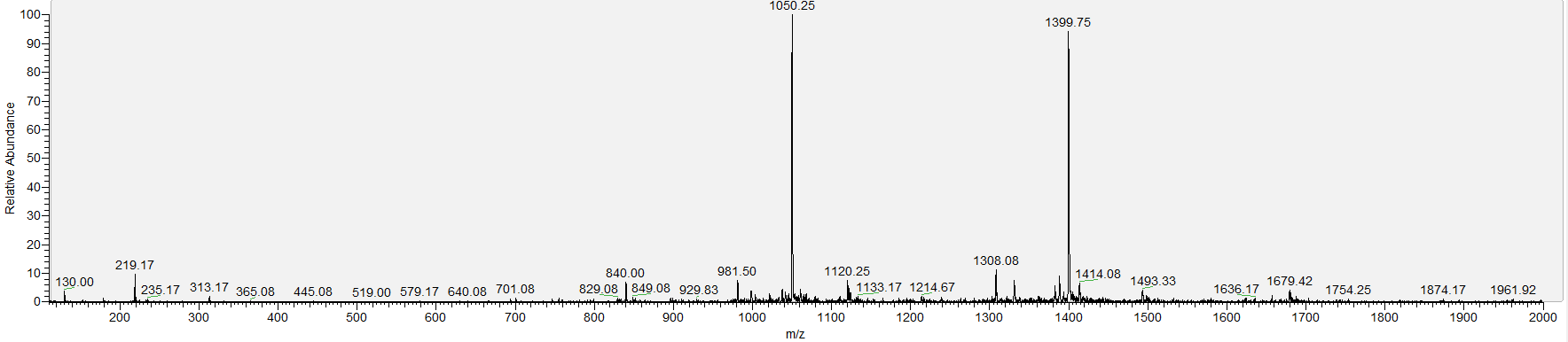

**HRMS;** [M+3H]^3+^: 1399.8491, [M+4H]^4+^: 1050.3862.

ii) **Variation of PNA length**

**E2**

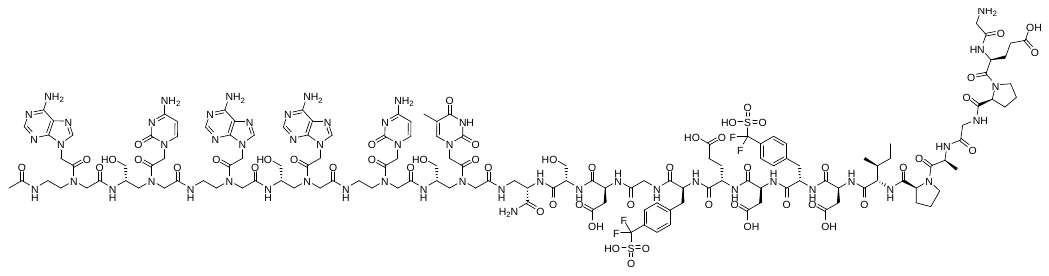

**Chemical Formula**: C_142_H_187_F_4_N_53_O_53_S_2_, **Exact Mass**: 3622.29, **Molecular Weight**: 3624.49

**MALDI-TOF**; m/z found: 3624.26

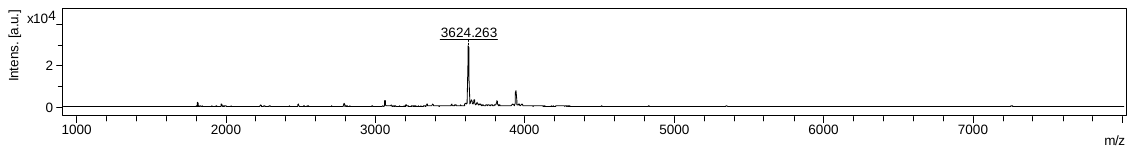

**LCMS (ESI)**; RT= 1.36 min, [M+3H]^3+^: 1208.75, [M+4H]^4+^: 906.83

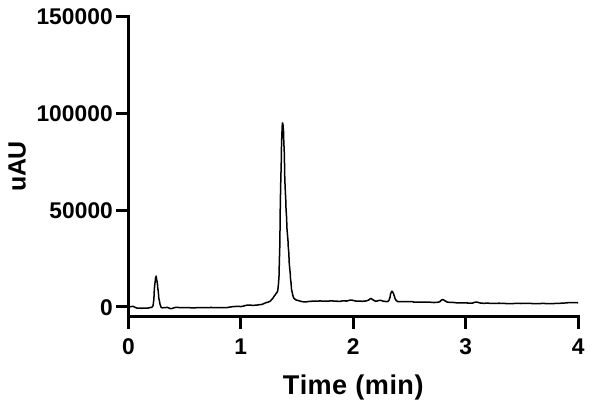

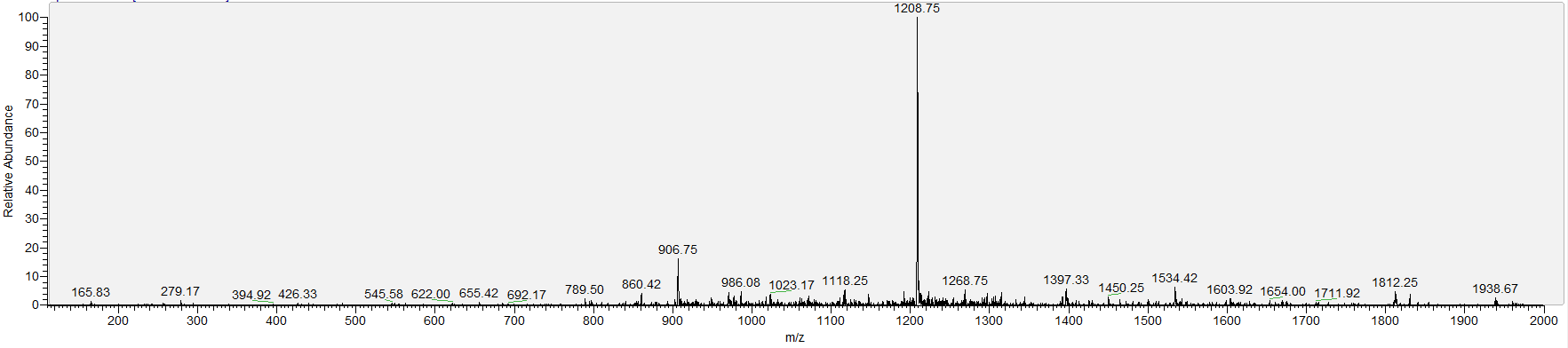

**A2**

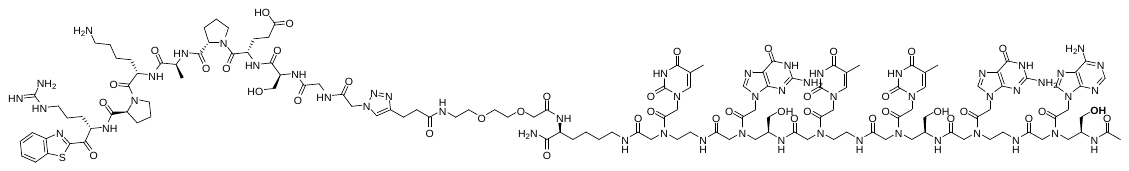

**Chemical Formula**: C_132_H_182_N_52_O_42_S, **Exact Mass**: 3199.34, **Molecular Weight**: 3201.29

**MALDI-TOF**; m/z found: 3201.17.

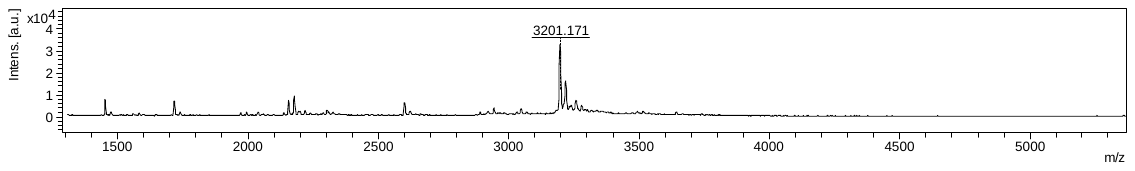

**LCMS (ESI)**; RT= 1.48 min, [M+2H]^2+^: 1600.75, [M+3H]^3+^: 1067.83, [M+4H]^4+^: 801.00, [M+5H]^5+^: 641.17.

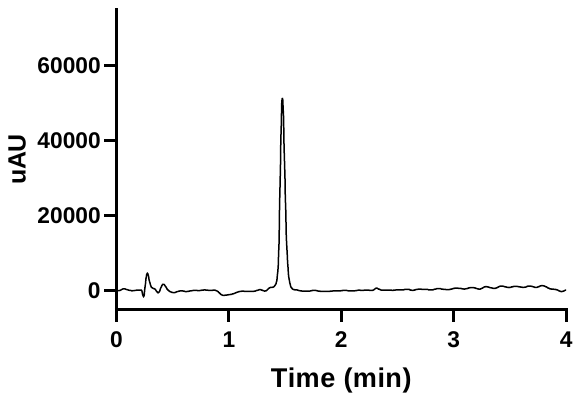

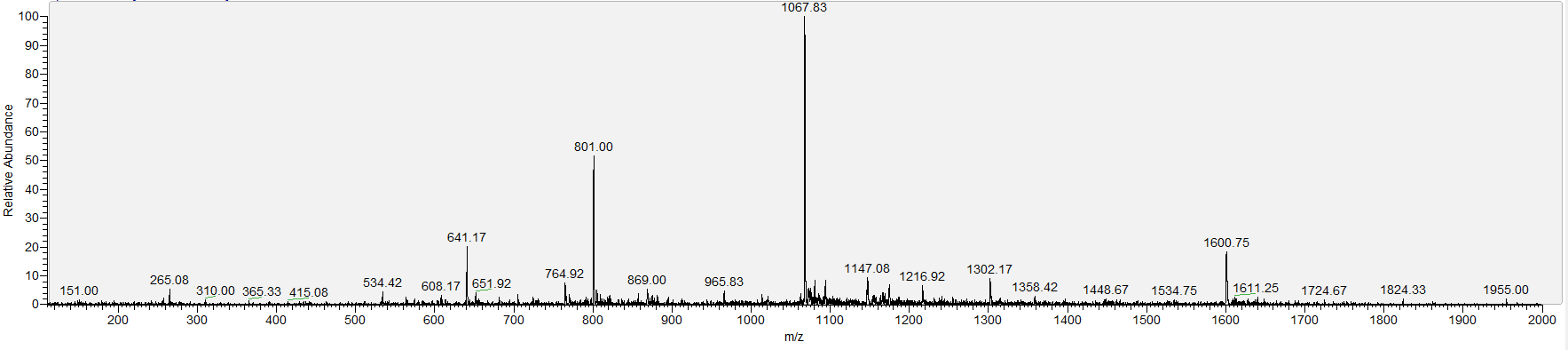

**E3**

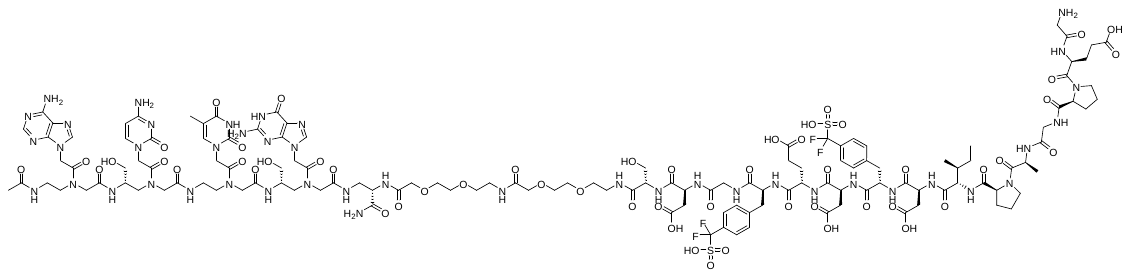

**Chemical Formula**: C_132_H_181_F_4_N_43_O_54_S_2_, **Exact Mass**: 3372.21, **Molecular Weight**: 3374.26.

**MALDI-TOF**; m/z found: 3373.96.

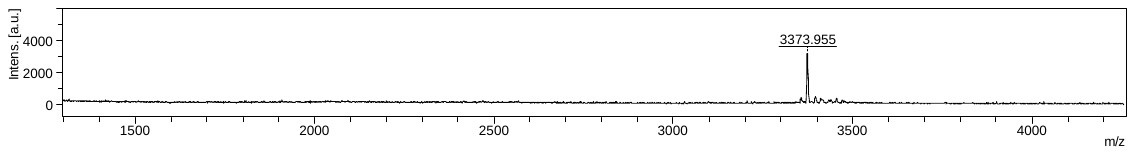

**LCMS (ESI)**; RT= 1.42 min, [M+2H]^2+^: 1687.58, [M+3H]^3+^: 1125.33.

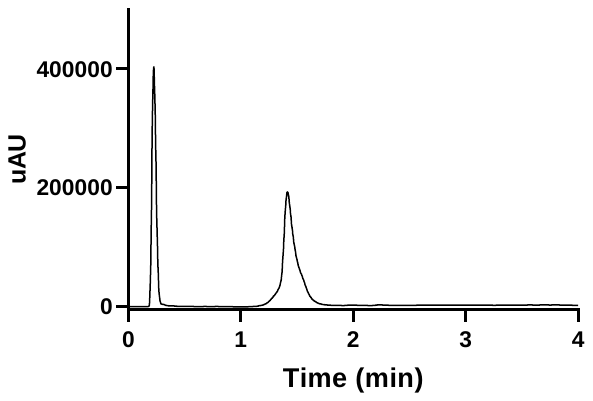

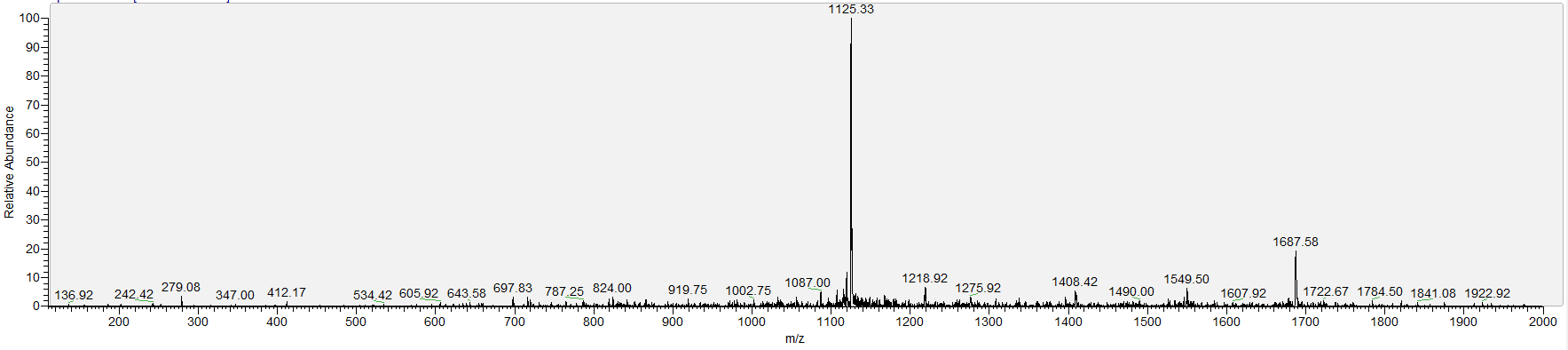

**A3**

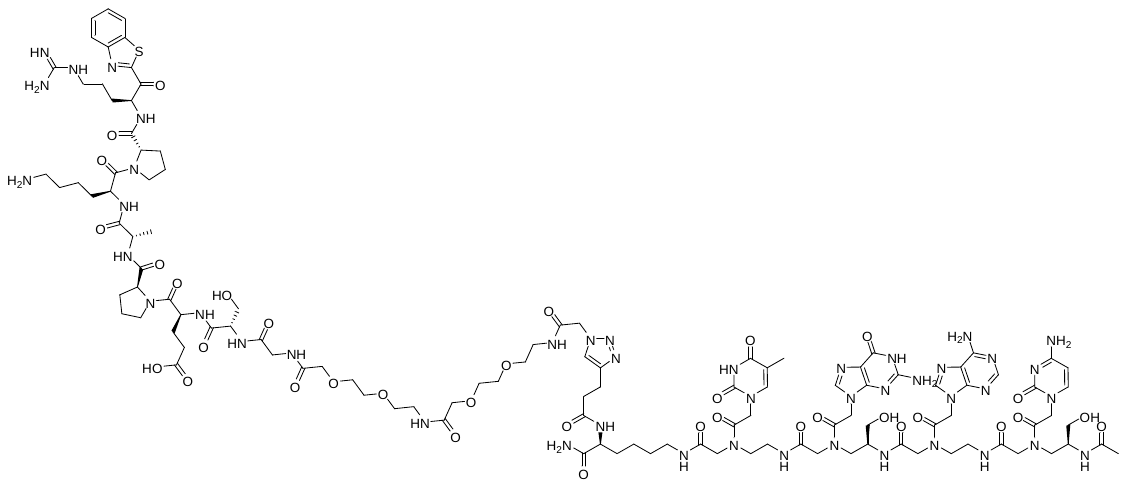

**Chemical Formula**: C_114_H_164_N_44_O_35_S, **Exact Mass**: 2741.21, **Molecular Weight**: 2742.90.

**MALDI-TOF**; m/z found: 2743.43.

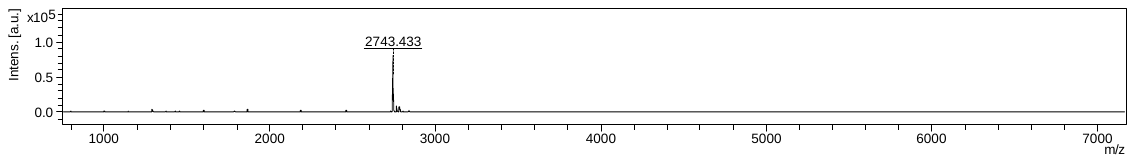

**LCMS (ESI)**; RT= 1.44 min, [M+2H]^2+^: 1372.00, [M+3H]^3+^: 915.17, [M+4H]^4+^: 686.58, [M+5H]^5+^: 549.50.

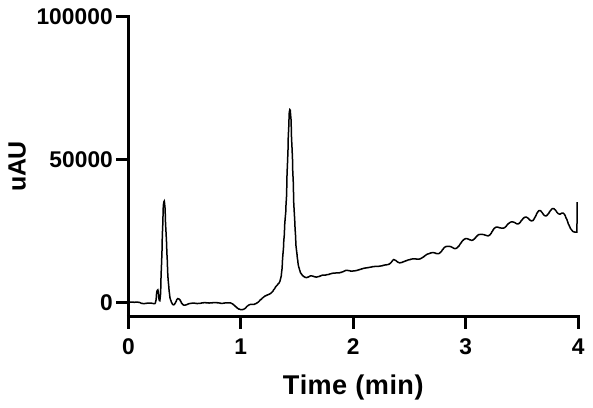

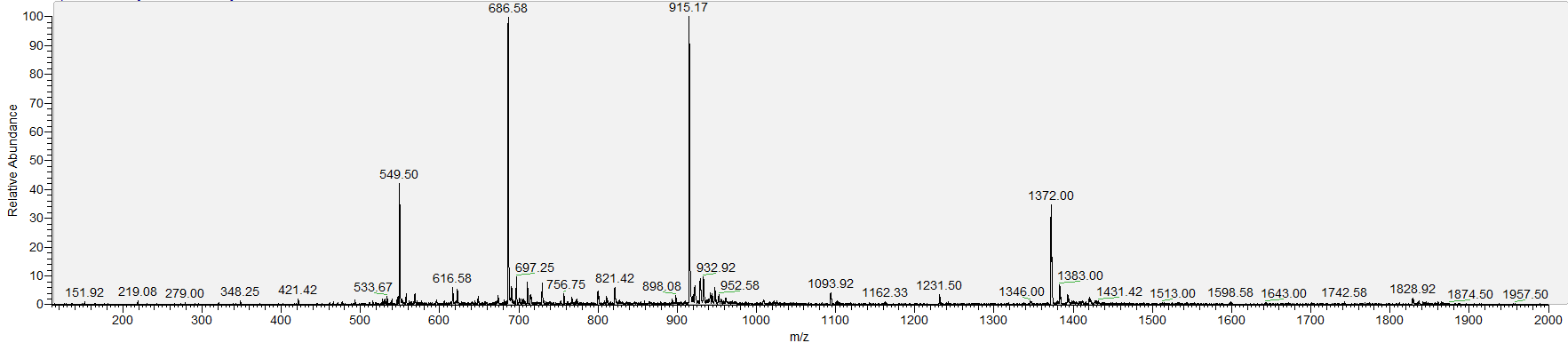

**A8**

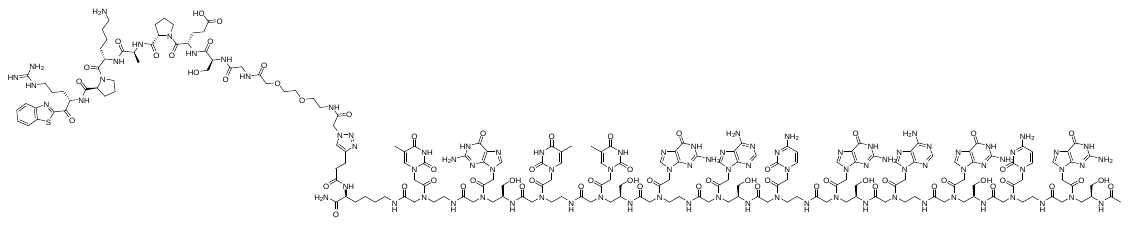

**Chemical Formula**: C_199_H_267_N_91_O_61_S, **Exact Mass**: 4939.03, **Molecular Weight**: 4941.96.

**MALDI-TOF**; m/z found: 4942.01.

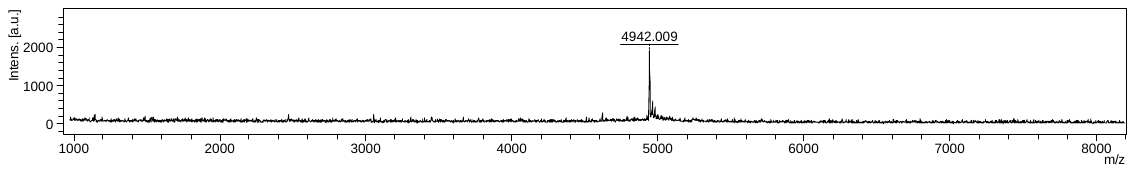

**LCMS (ESI)**; RT= 1.49 min, [M+3H]^3+^: 1647.67, [M+4H]^4+^: 1236.25, [M+5H]^5+^: 969.17, [M+6H]^6+^: 824.50, [M+7H]^7+^:706.67.

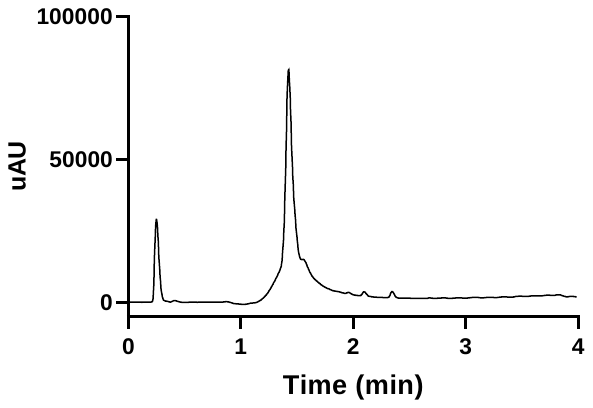

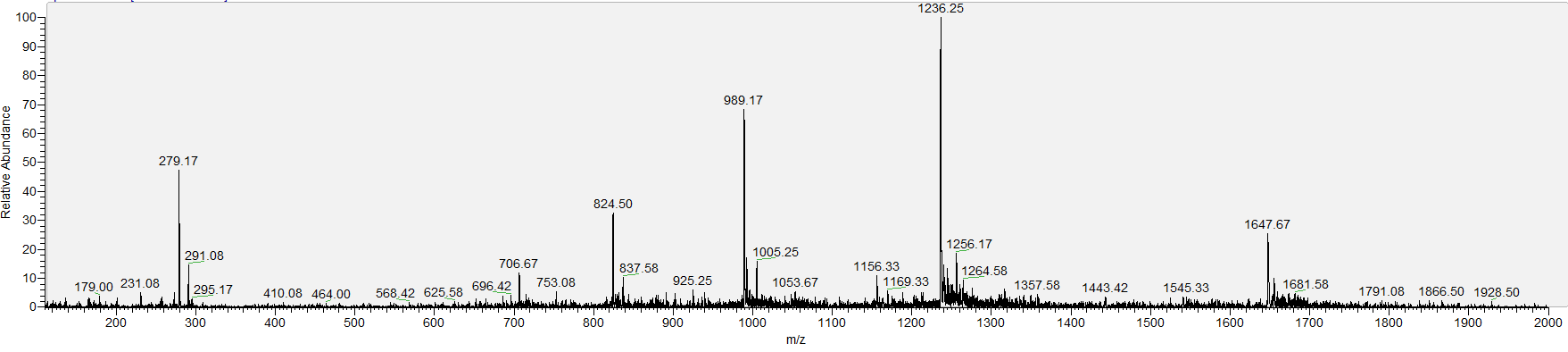

**HRMS;** [M+3H]^3+^: 1648.3586, [M+4H]^4+^: 1236.2682, [M+5H]^5+^: 989.2154, [M+6H]^6+^: 824.5084, [M+7H]^7+^:706.8661.

iii) **Charge modification on exosite II binder**

**E4**

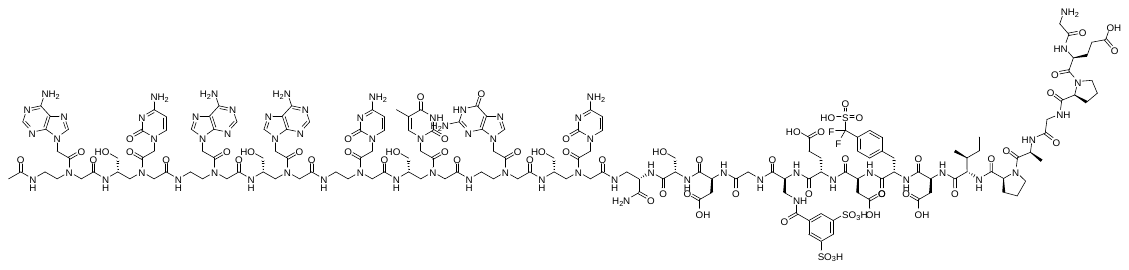

**Chemical Formula**: C_164_H_216_F_2_N_66_O_64_S_3_, **Exact Mass**: 4267.48, **Molecular Weight**: 4270.11.

**LCMS (ESI)**; RT= 1.48 min, [M+3H]^3+^: 1424.00, [M+4H]^4+^: 1068.33.

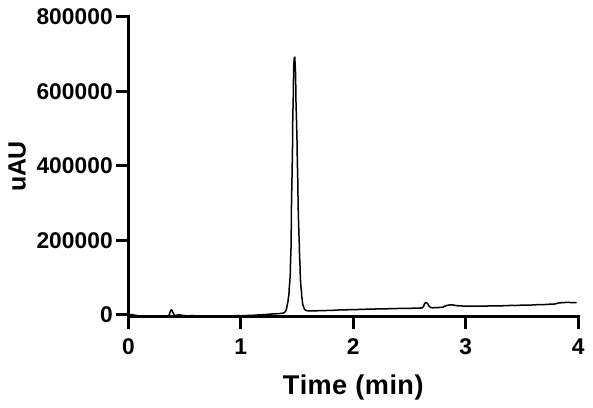

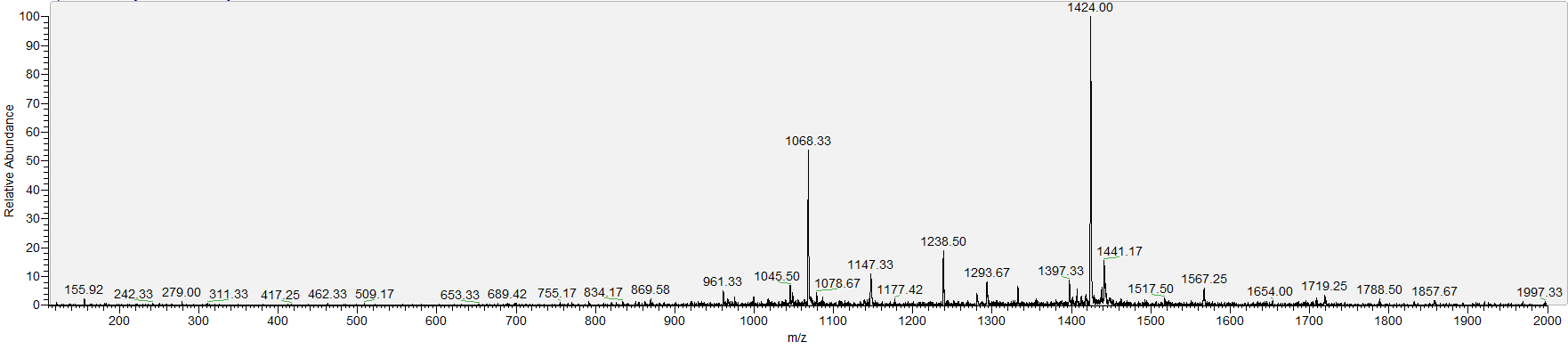

**E5**

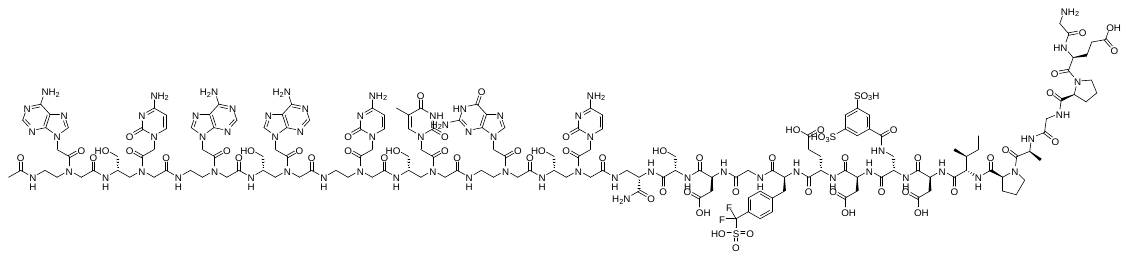

**Chemical Formula**: C_164_H_216_F_2_N_66_O_64_S_3_, **Exact Mass**: 4267.48, **Molecular Weight**: 4270.11.

**LCMS (ESI)**; RT= 1.30 min, [M+3H]^3+^: 1424.08, [M+4H]^4+^: 1068.25.

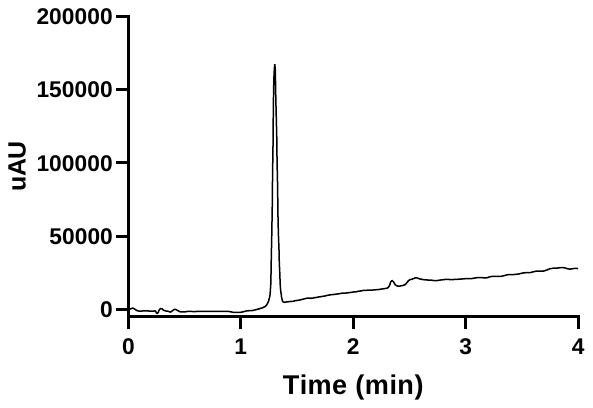

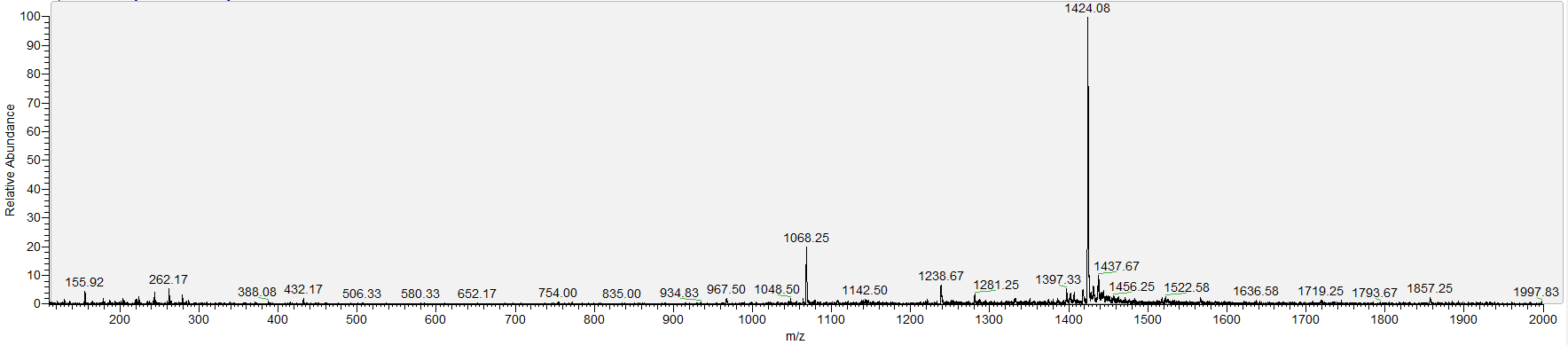

**E6**

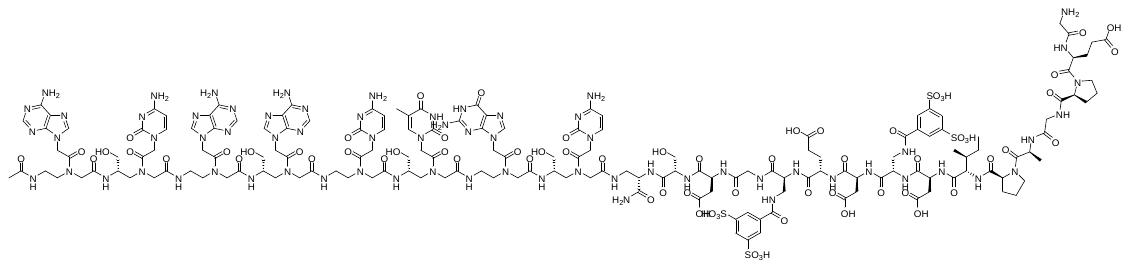

**Chemical Formula**: C_164_H_217_N_67_O_68_S_4_, **Exact Mass**: 4340.45, **Molecular Weight**: 4343.18.

**LCMS (ESI)**; RT= 1.24 min, [M+3H]^3+^: 1446.33, [M+4H]^4+^: 1086.50.

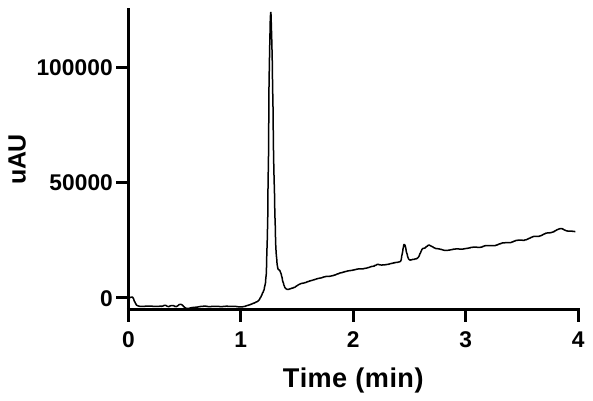

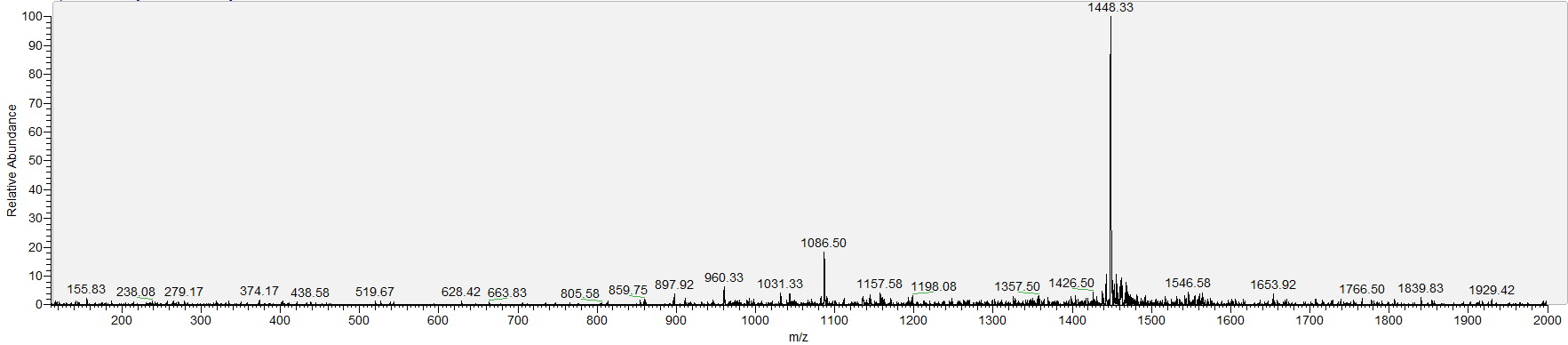

**E7**

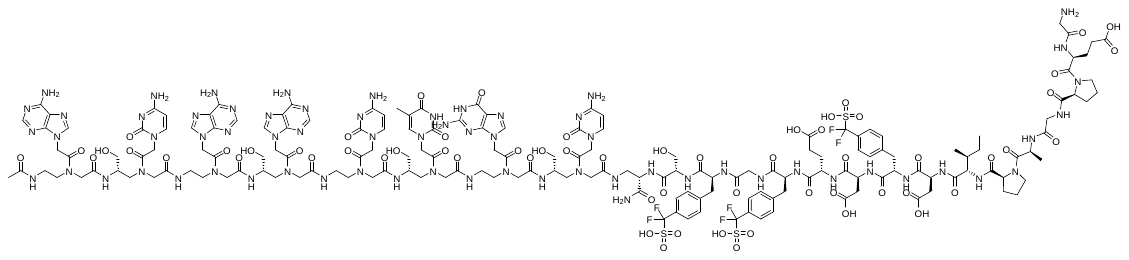

**Chemical Formula**: C_170_H_219_F_6_N_65_O_61_S_3_, **Exact Mass**: 4356.51, **Molecular Weight**: 4359.19

**MALDI-TOF**; m/z found: 4359.75

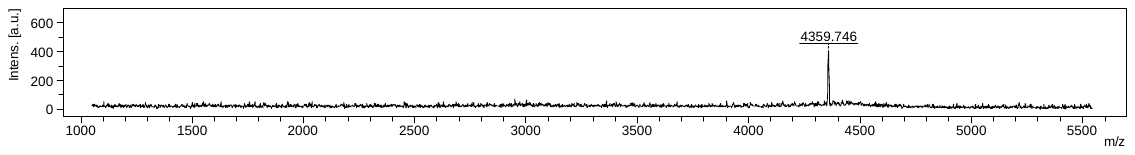

**LCMS (ESI)**; RT= 1.42 min, [M+3H]^3+^: 1453.67, [M+4H]^4+^: 1090.67.

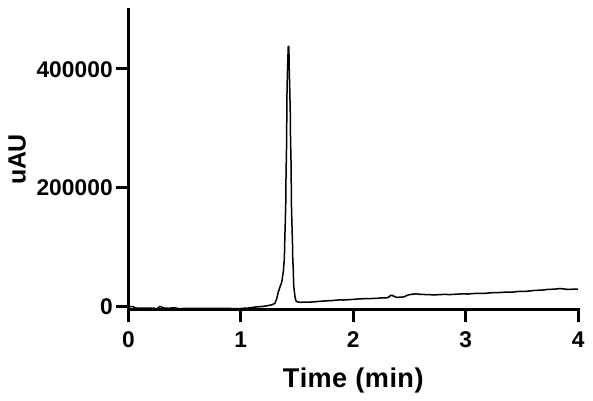

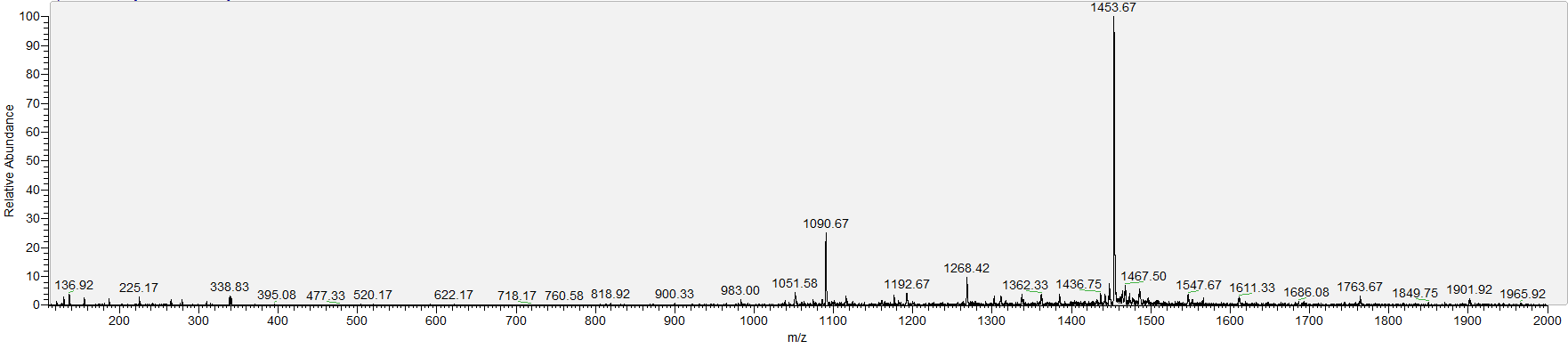

**E8**

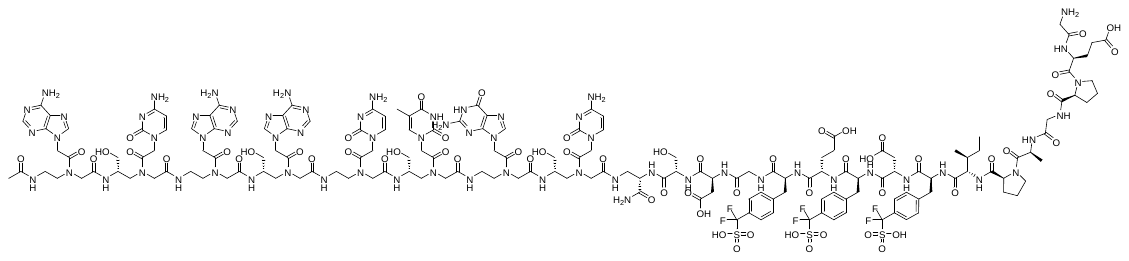

**Chemical Formula**: C_170_H_219_F_6_N_65_O_61_S_3_, **Exact Mass**: 4356.51, **Molecular Weight**: 4359.19

**MALDI-TOF**; m/z found: 4359.70

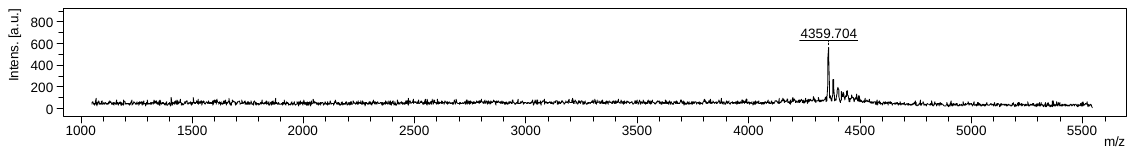

**LCMS (ESI)**; RT= 1.46 min, [M+3H]^3+^: 1453.75, [M+4H]^4+^: 1090.33.

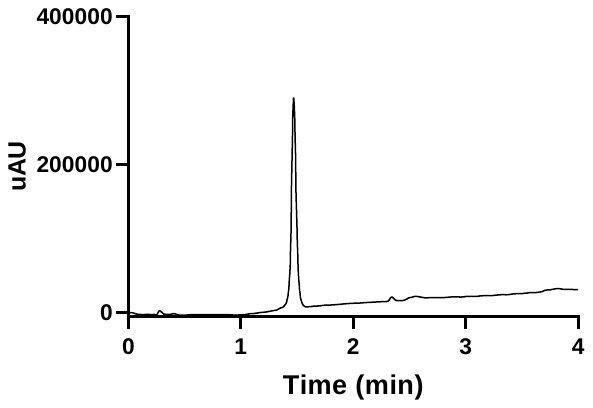

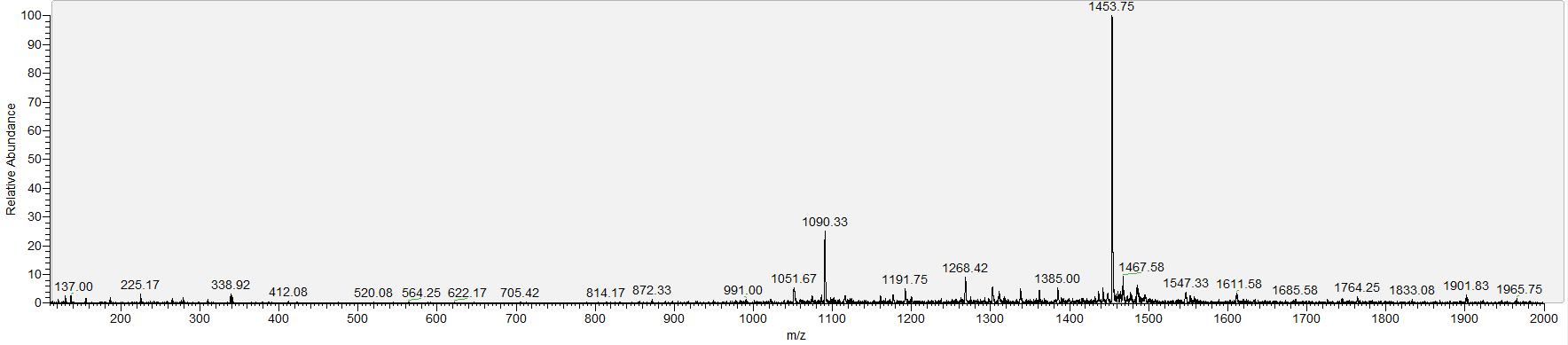

iv) **Hydrophobic residues**

**E9**

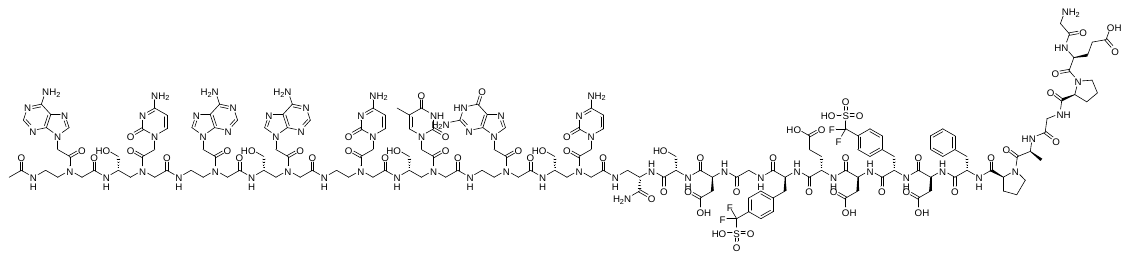

**Chemical Formula**: C_167_H_213_F_4_N_65_O_60_S_2_, **Exact Mass**: 4228.50, **Molecular Weight**: 4231.05.

**MALDI-TOF**; m/z found: 4230.99.

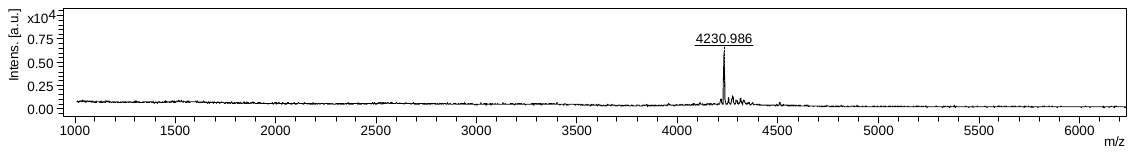

**LCMS (ESI)**; RT= 1.48 min, [M+3H]^3+^: 1411.08, [M+4H]^4+^: 1058.33.

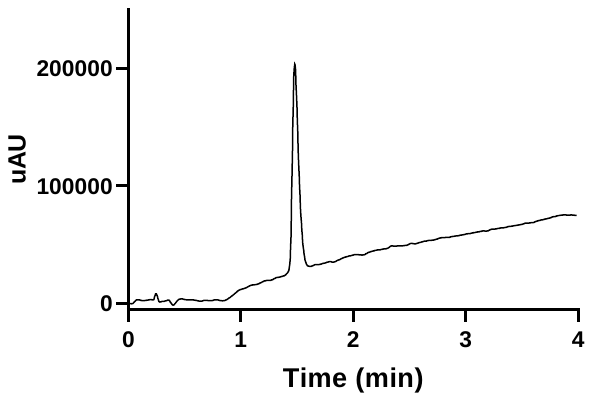

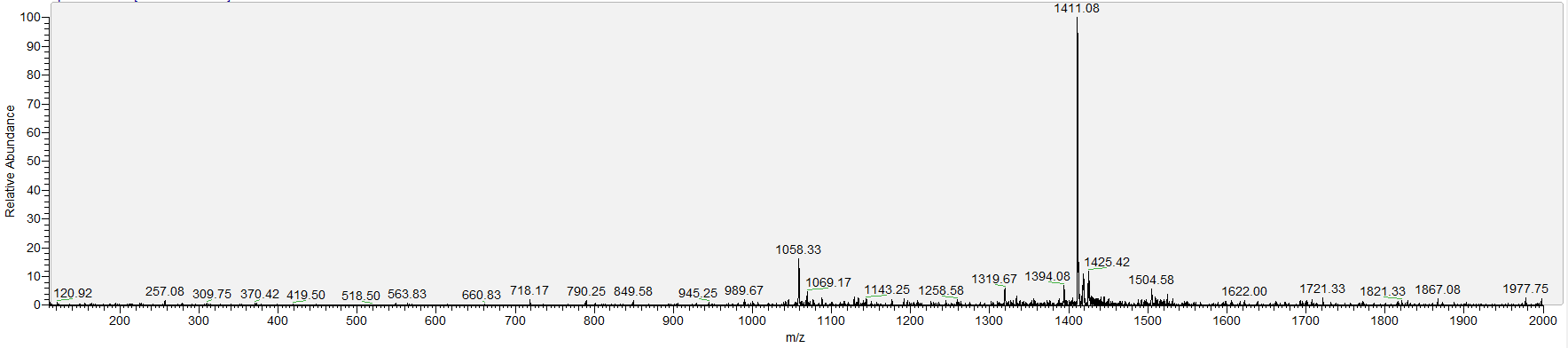

**E10**

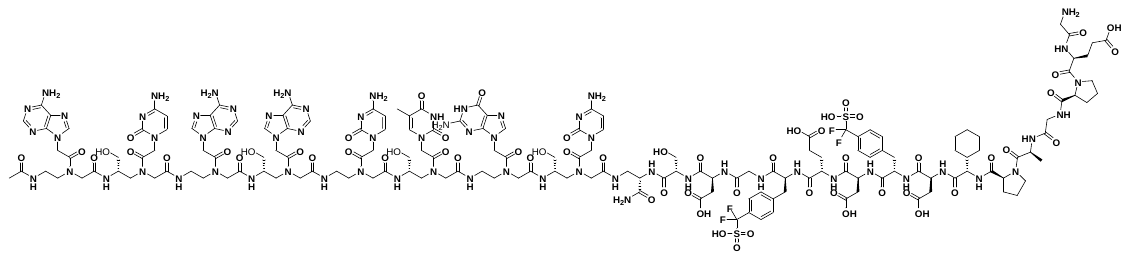

**Chemical Formula**: C_166_H_217_F_4_N_65_O_60_S_2_, **Exact Mass**: 4220.53, **Molecular Weight**: 4223.07.

**MALDI-TOF**; m/z found: 4223.40.

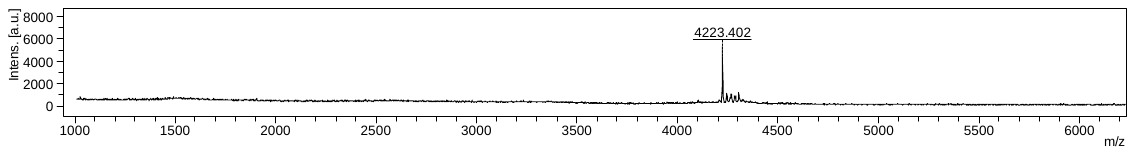

**LCMS (ESI)**; RT= 1.48 min, [M+3H]^3+^: 1408.42, [M+4H]^4+^: 1056.50.

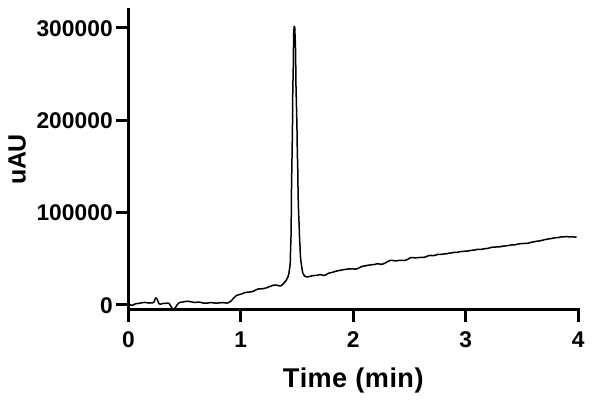

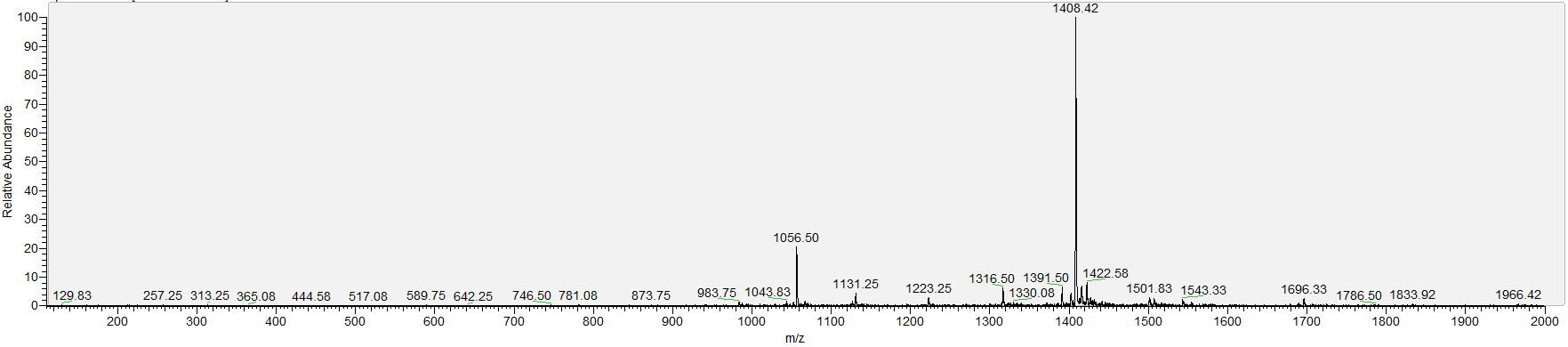

**E11**

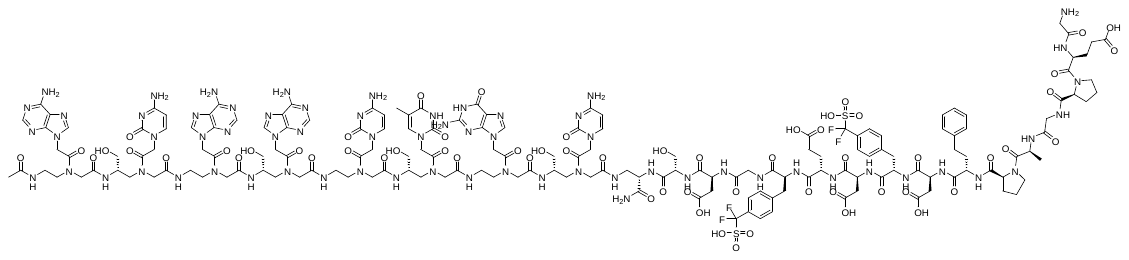

**Chemical Formula**: C_168_H_215_F_4_N_65_O_60_S_2_, **Exact Mass**: 4242.52, **Molecular Weight**: 4245.08.

**MALDI-TOF**; m/z found: 4245.59.

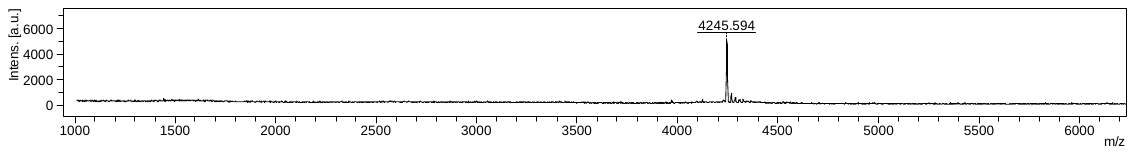

**LCMS (ESI)**; RT= 1.51 min, [M+3H]^3+^: 1415.83, [M+4H]^4+^: 1062.17.

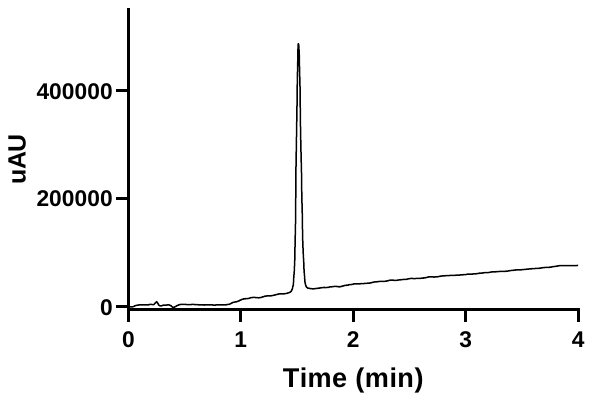

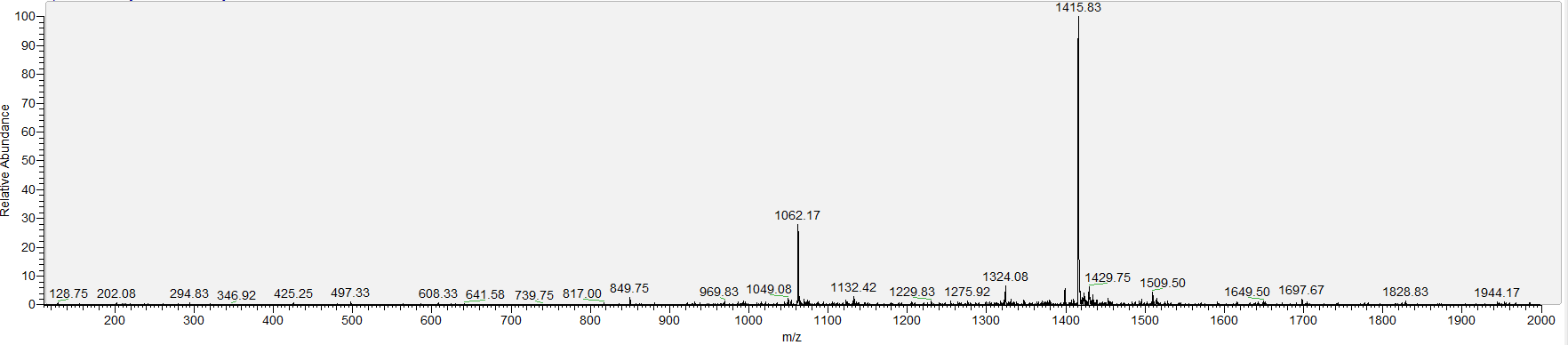

**E12**

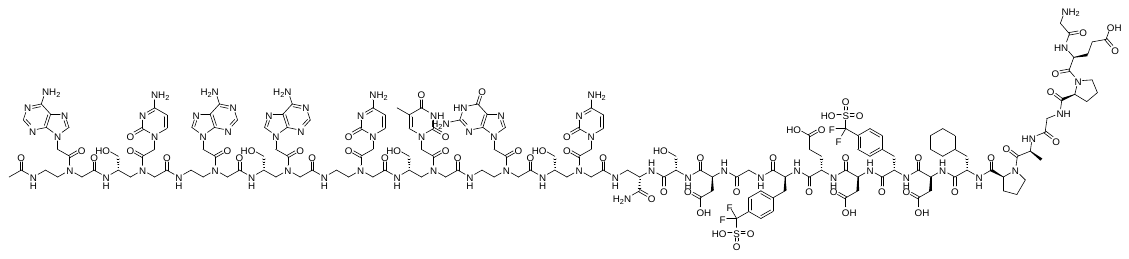

**Chemical Formula**: C_167_H_219_F_4_N_65_O_60_S_2_, **Exact Mass**: 4234.55, **Molecular Weight**: 4237.10

**MALDI-TOF**; m/z found: 4237.21.

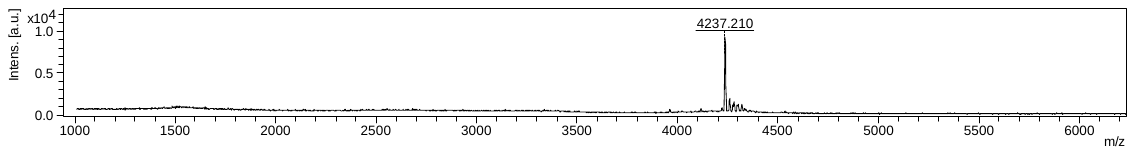

**LCMS (ESI)**; RT= 1.56 min, [M+3H]^3+^: 1413.00, [M+4H]^4+^: 1059.92.

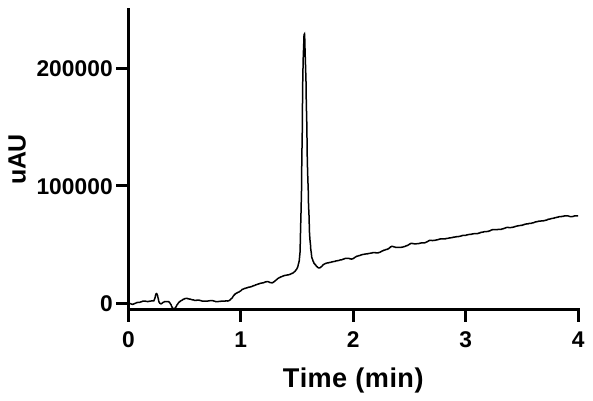

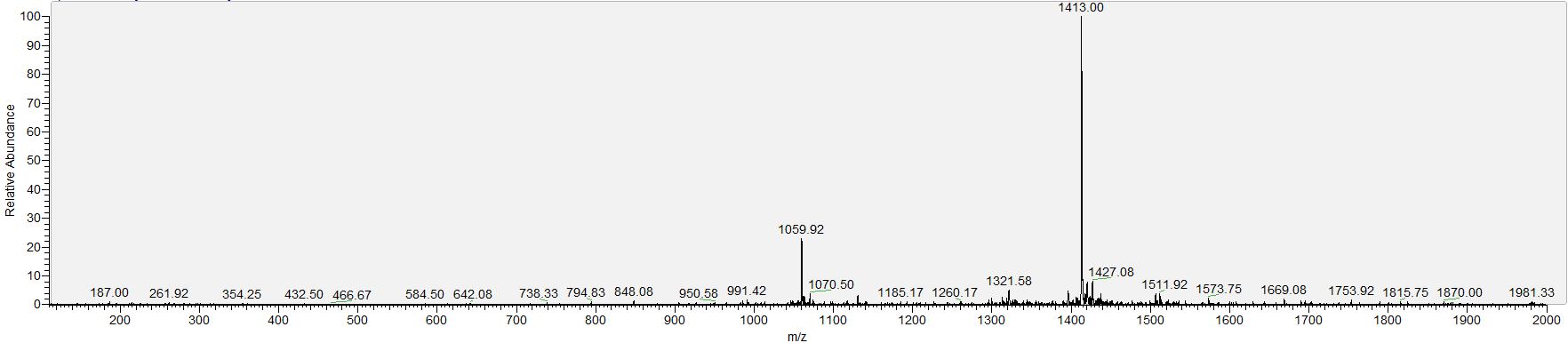

**E13**

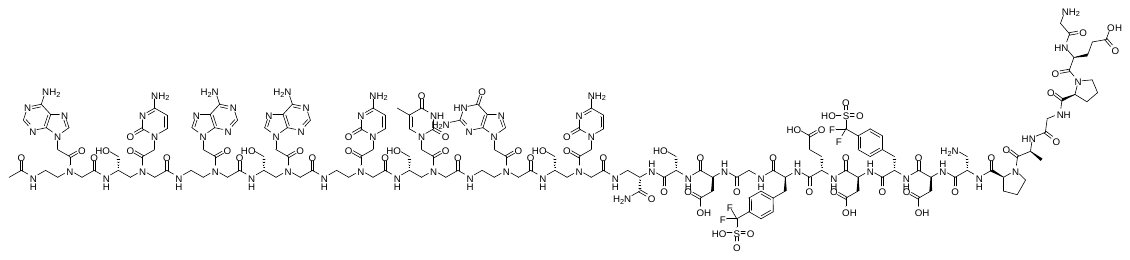

**Chemical Formula**: C_161_H_210_F_4_N_66_O_60_S_2_, **Exact Mass**: 4167.48, **Molecular Weight**: 4169.97.

**MALDI-TOF**; m/z found: 4169.95.

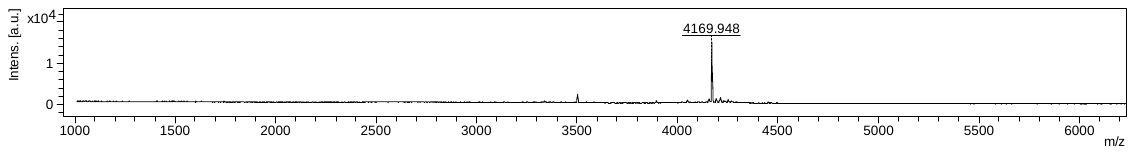

**LCMS (ESI)**; RT= 1.35 min, [M+3H]^3+^: 1390.75, [M+4H]^4+^: 1043.17.

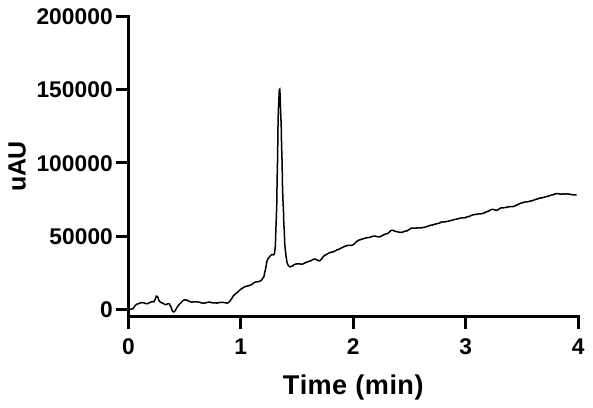

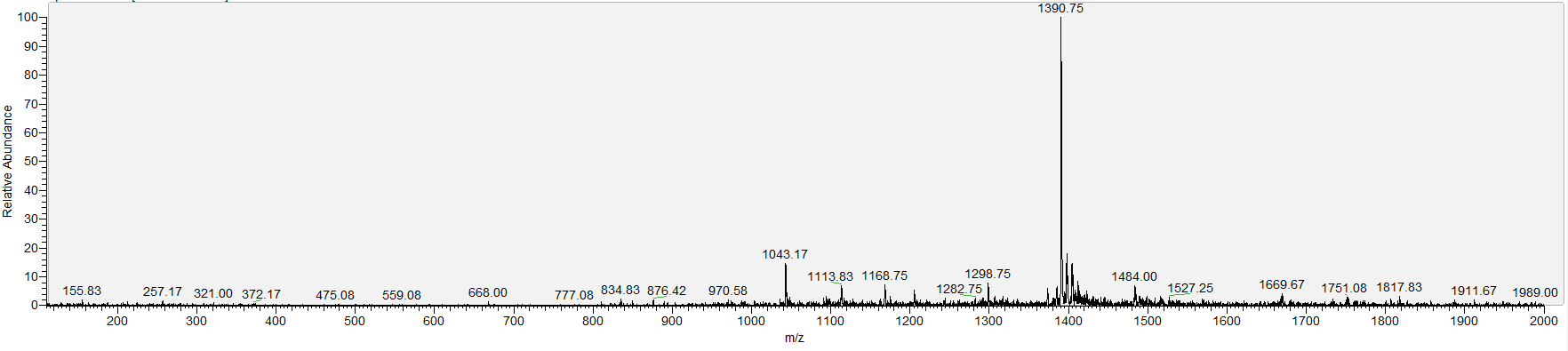

**E14**

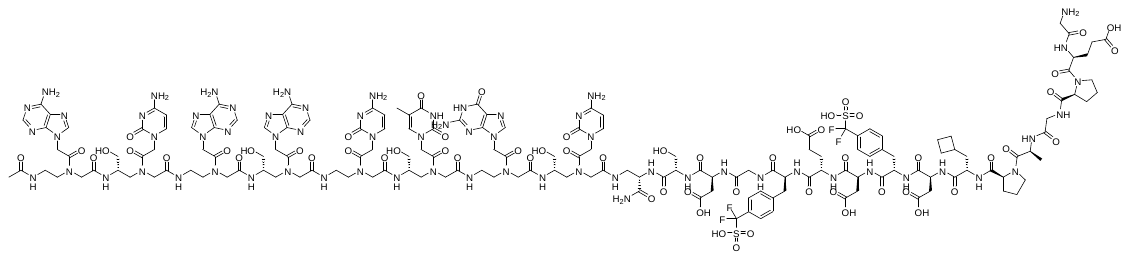

**Chemical Formula**: C_165_H_215_F_4_N_65_O_60_S_2_, **Exact Mass**: 4206.52, **Molecular Weight**: 4209.04.

**MALDI-TOF**; m/z found: 4209.36.

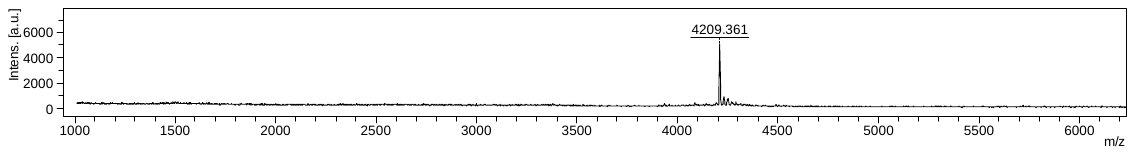

**LCMS (ESI)**; RT= 1.46 min, [M+3H]^3+^: 1403.75, [M+4H]^4+^: 1052.75.

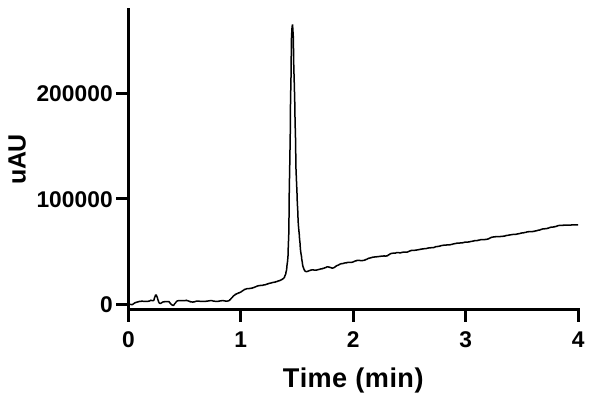

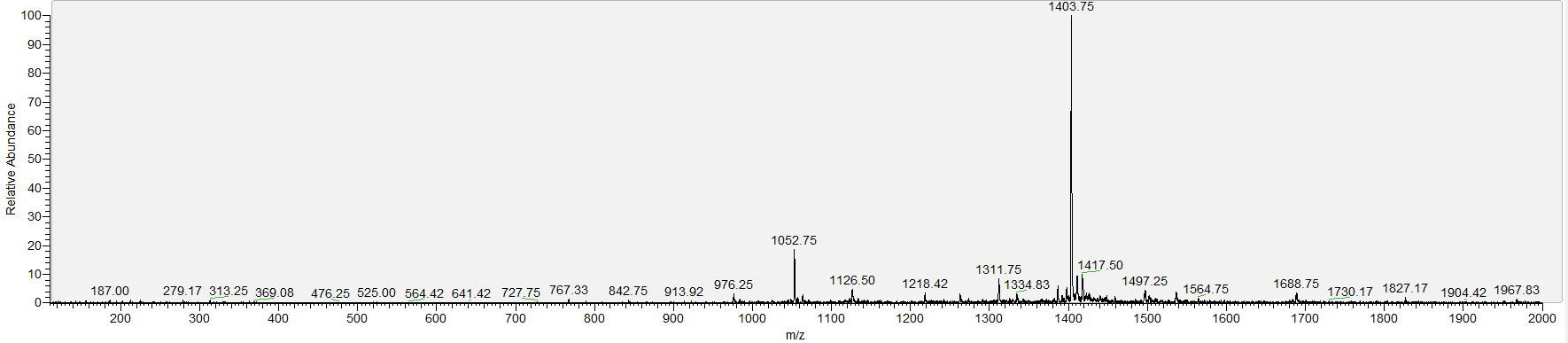

**E15**

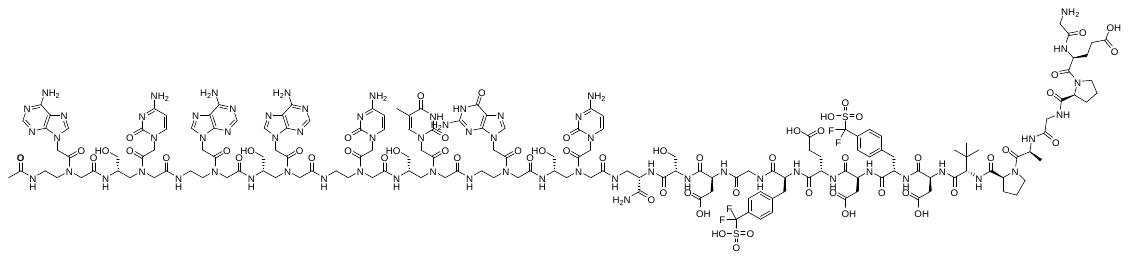

**Chemical Formula**: C164H215F4N65O60S2, **Exact Mass**: 4194.52, **Molecular Weight**: 4197.03.

**MALDI-TOF**; m/z found: 4196.96

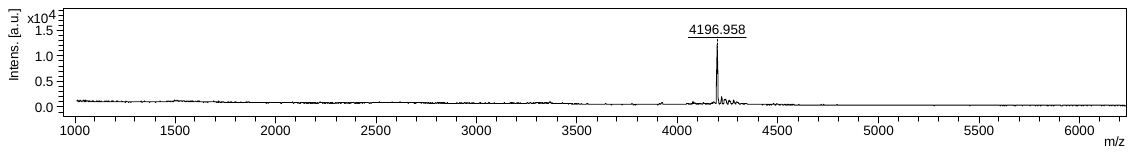

**LCMS (ESI)**; RT= 1.42 min, [M+3H]^3+^: 1399.67, [M+4H]^4+^: 1050.17.

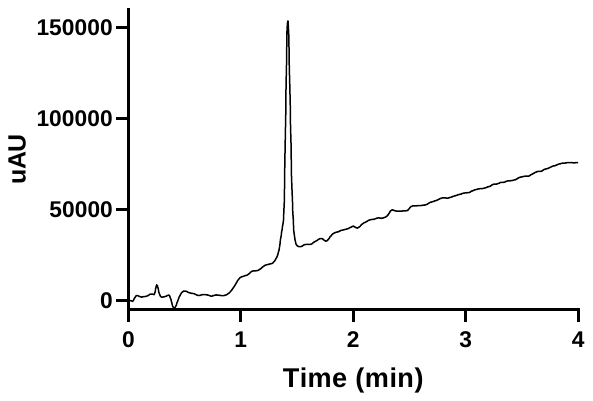

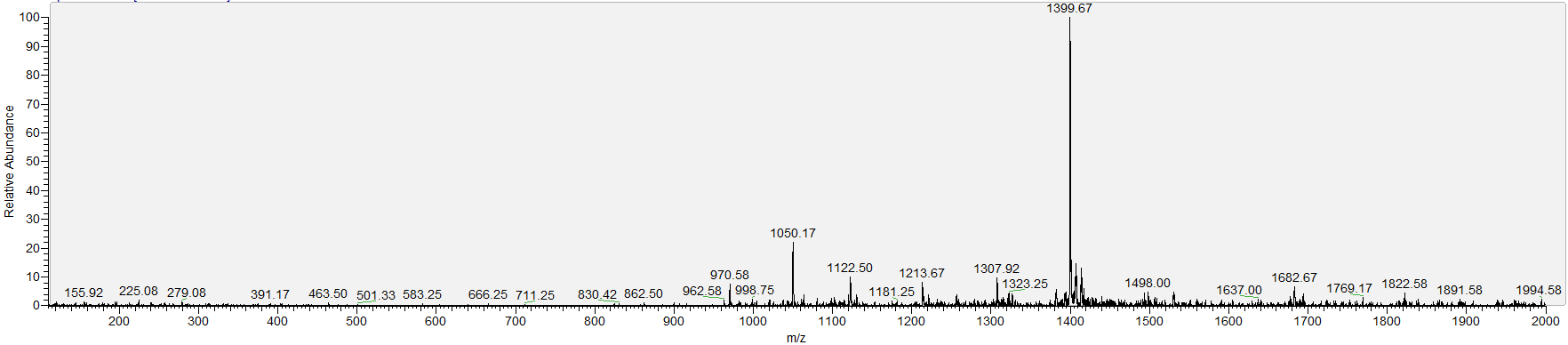

**E16**

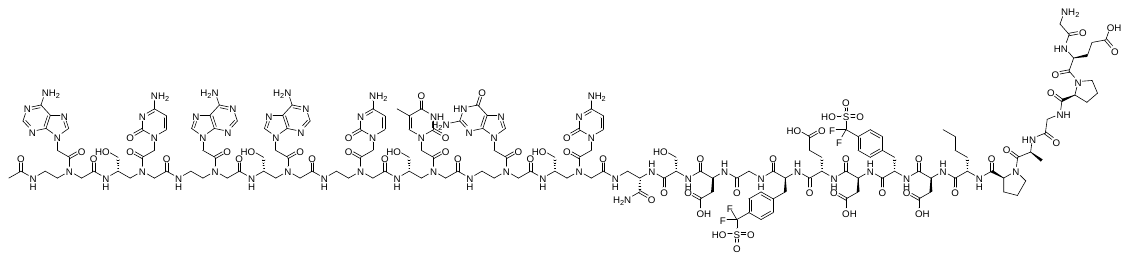

**Chemical Formula**: C_164_H_215_F_4_N_65_O_60_S_2_, **Exact Mas**s: 4194.52, **Molecular Weight**: 4197.03

**MALDI-TOF**; m/z found: 4197.05.

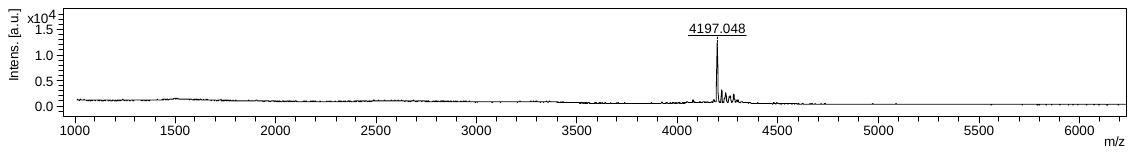

**LCMS (ESI)**; RT= 1.44 min, [M+3H]^3+^: 1399.75, [M+4H]^4+^: 1049.75.

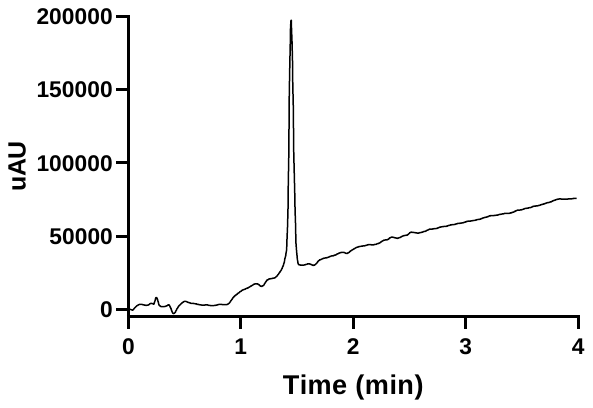

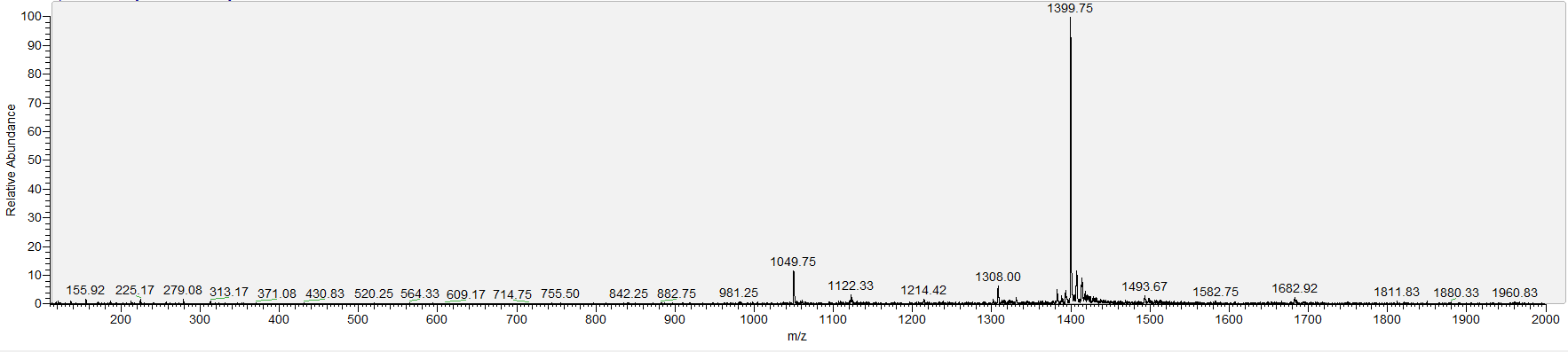

i) **Ala scan**

**E17**

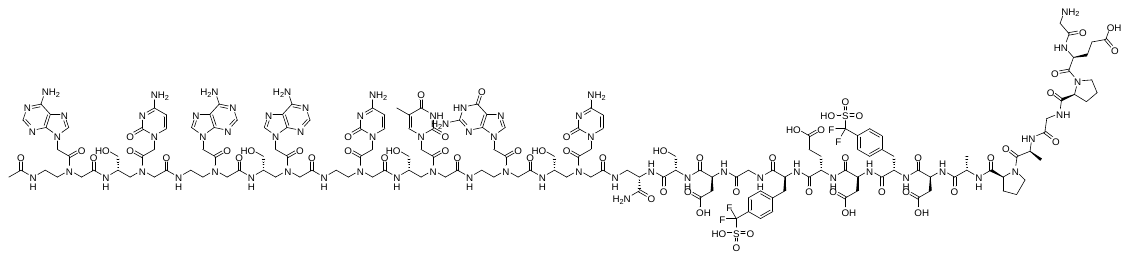

**Chemical Formula**: C_161_H_209_F_4_N_65_O_60_S_2_, **Exact Mass**: 4152.47, **Molecular Weight**: 4154.95

**MALDI-TOF**; m/z found: 4154.51

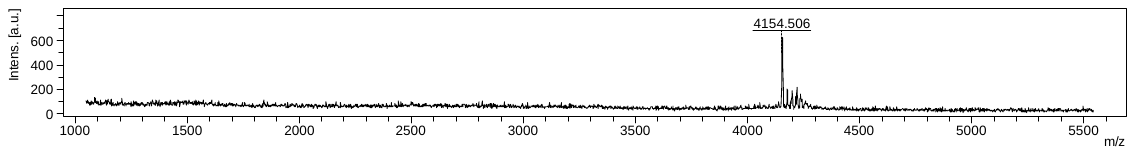

**LCMS (ESI)**; RT= 1.32 min, [M+3H]^3+^: 1385.67, [M+4H]^4+^: 1039.33.

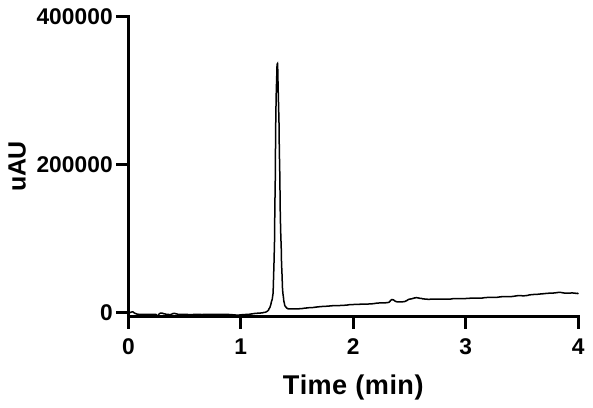

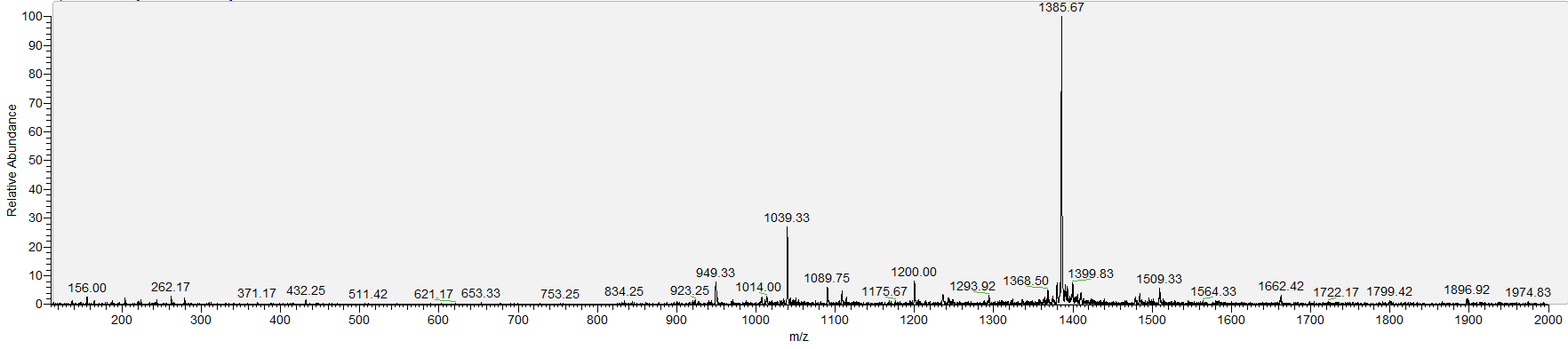

**E18**

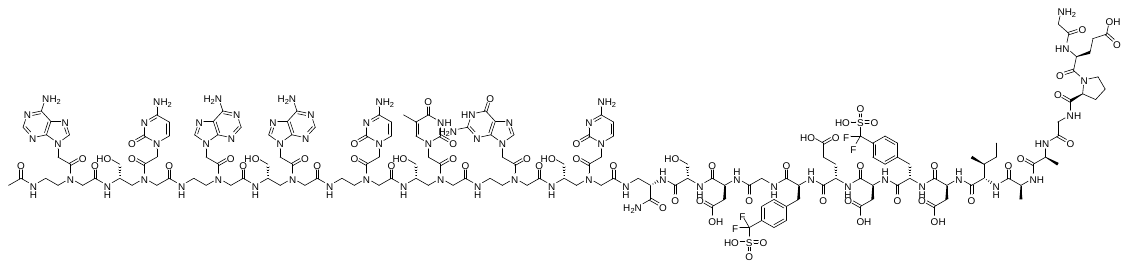

**Chemical Formula**: C_162_H_213_F_4_N_65_O_60_S_2_, **Exact Mass**: 4168.50, **Molecular Weight**: 4171.00

**MALDI-TOF**; m/z found: 4170.71

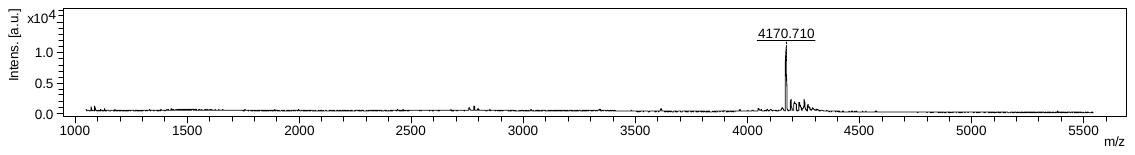

**LCMS (ESI)**; RT= 1.38 min, [M+3H]^3+^: 1391.08, [M+4H]^4+^: 1043.75.

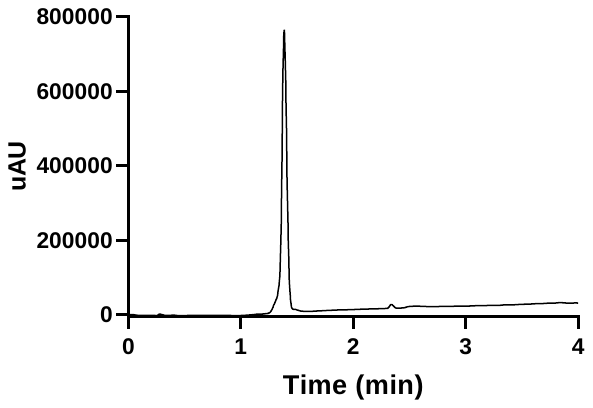

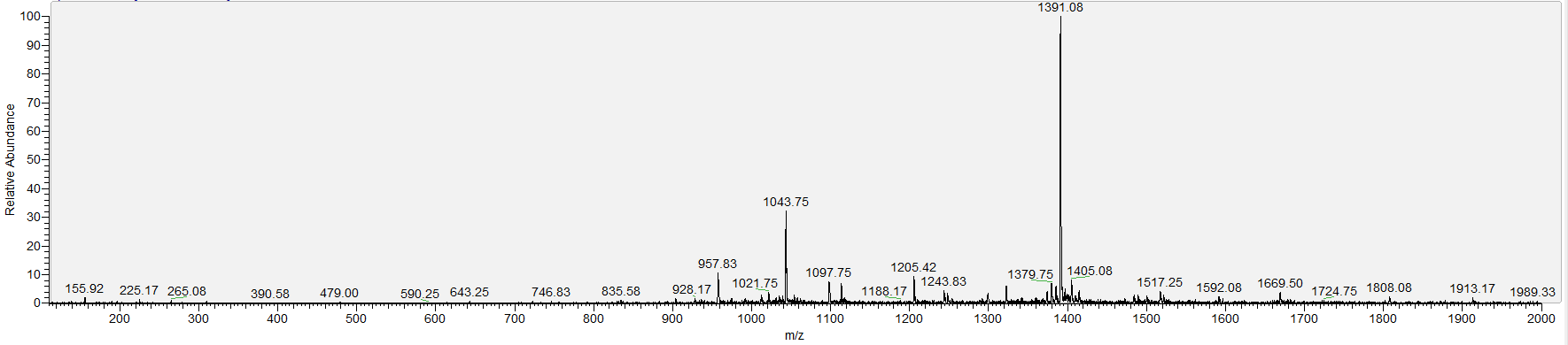

**E19**

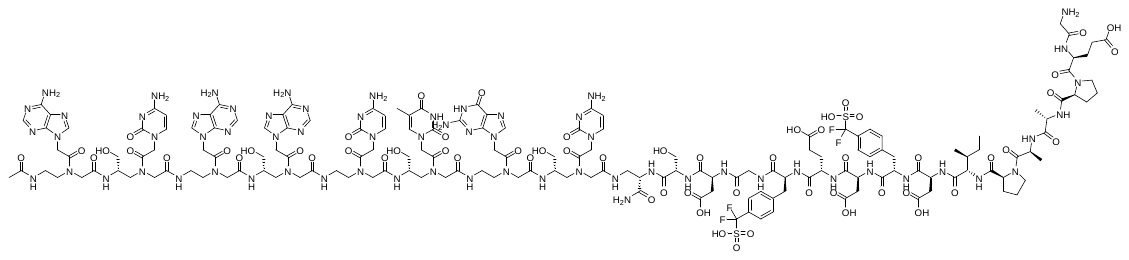

**Chemical Formula**: C_165_H_217_F_4_N_65_O_60_S_2_, **Exact Mass**: 4208.53, **Molecular Weight**: 4211.06

**MALDI-TOF**; m/z found: 4211.61.

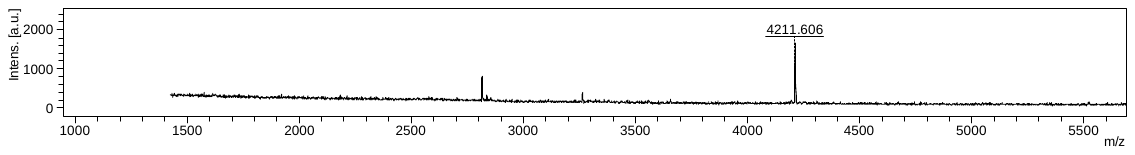

**LCMS (ESI)**; RT= 1.42 min, [M+3H]^3+^: 1404.17, [M+4H]^4+^: 1053.58.

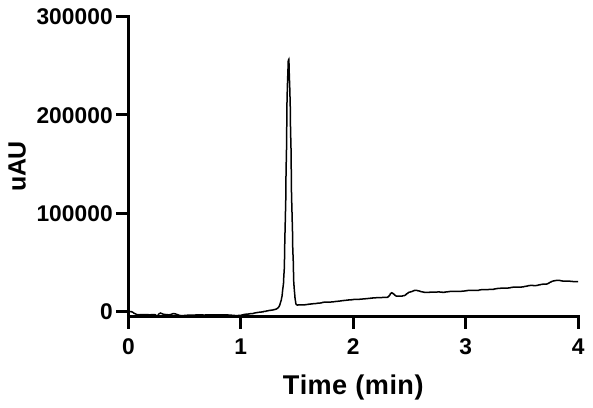

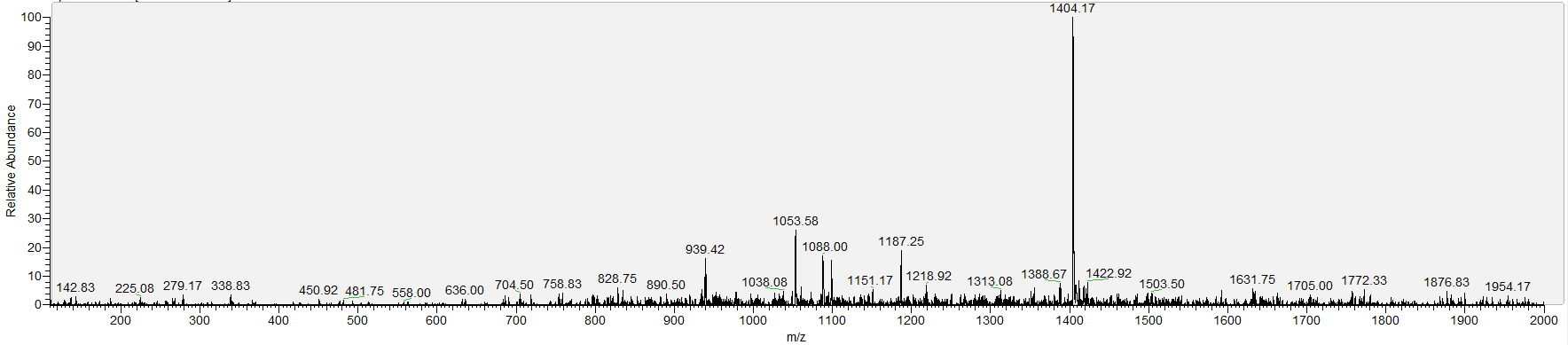

**E20**

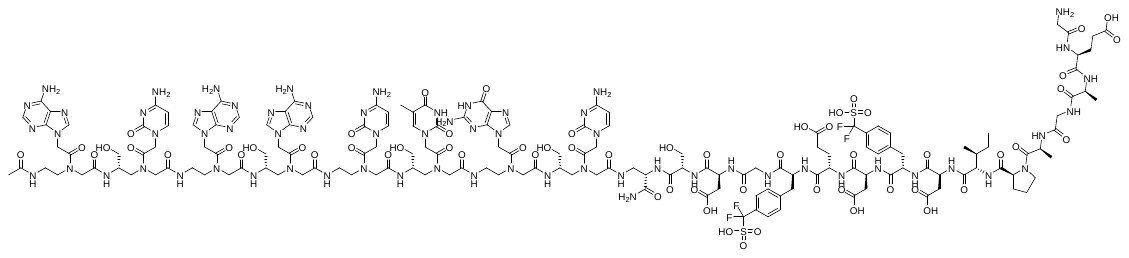

**Chemical Formula**: C_162_H_213_F_4_N_65_O_60_S_2_, **Exact Mass**: 4168.50, Molecular Weight: 4171.00

**MALDI-TOF**; m/z found: 4171.11.

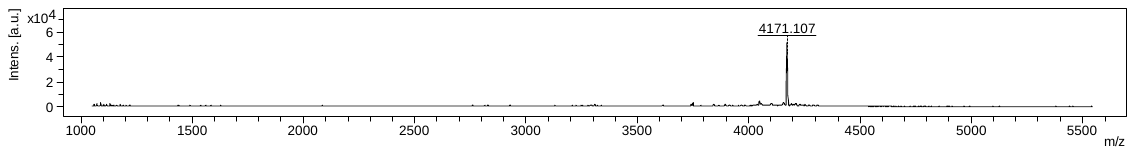

**LCMS (ESI)**; RT= 1.39 min, [M+3H]^3+^: 1391.25, [M+4H]^4+^: 1043.50.

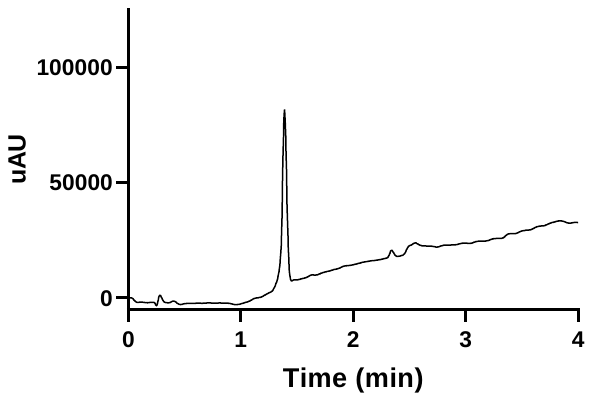

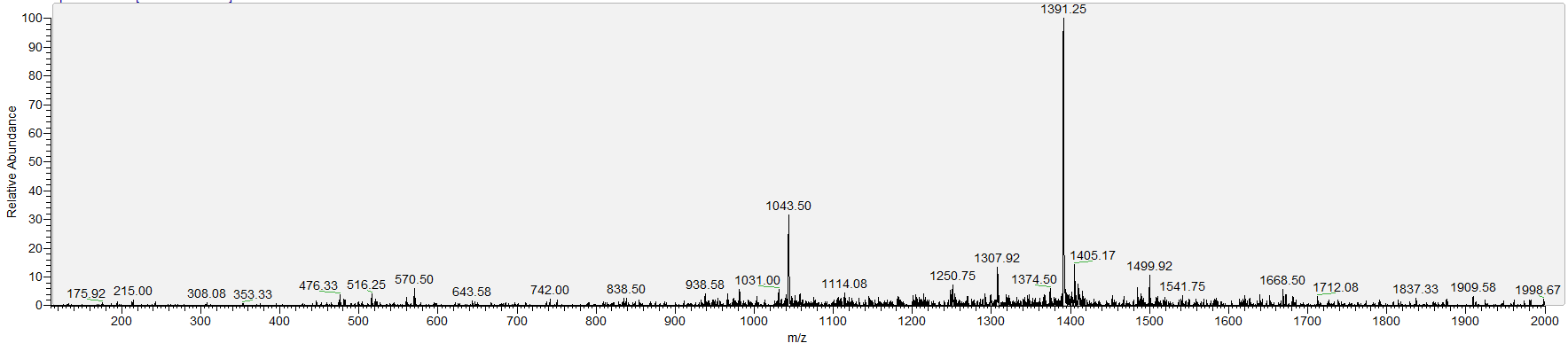

**E21**

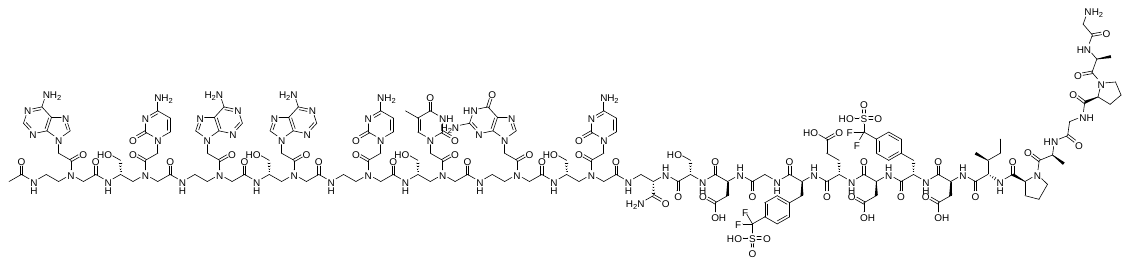

**Chemical Formula**: C_162_H_213_F_4_N_65_O_58_S_2_, **Exact Mass**: 4136.51, **Molecular Weight**: 4139.00.

**MALDI-TOF**; m/z found: 4139.19.

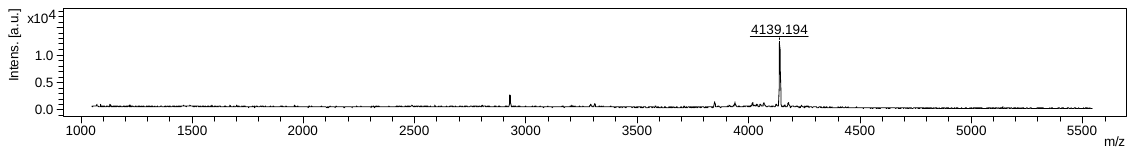

**LCMS (ESI)**; RT= 1.39 min, [M+3H]^3+^: 1380.42, [M+4H]^4+^: 1035.67.

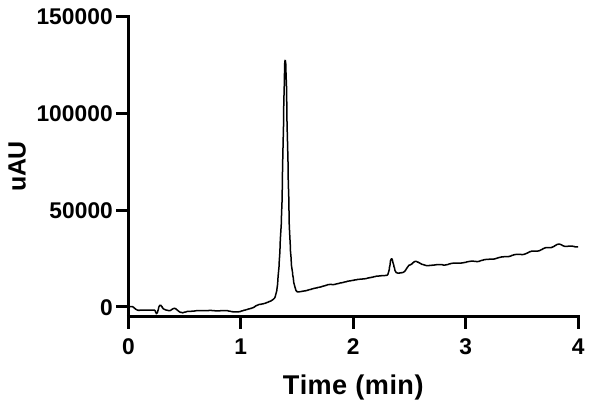

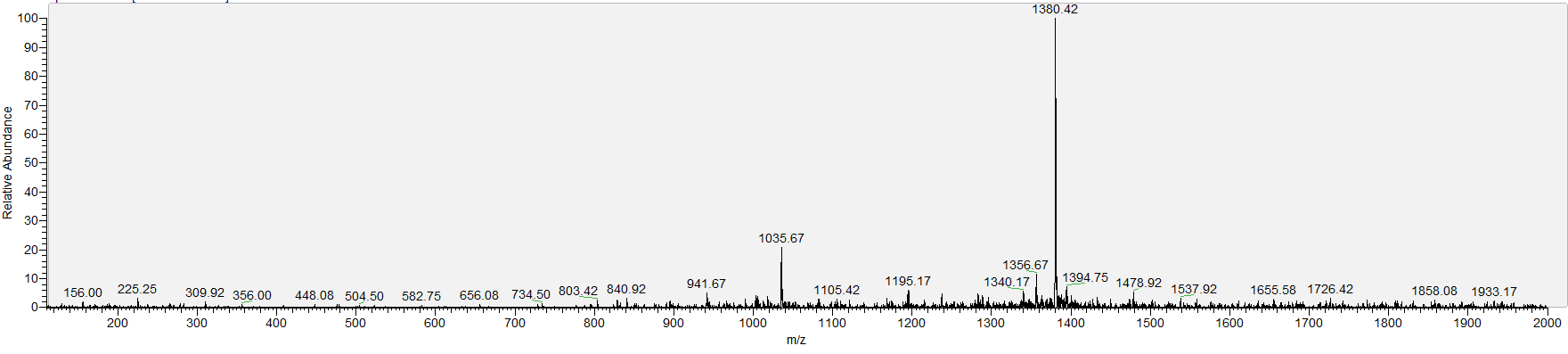

**E22**

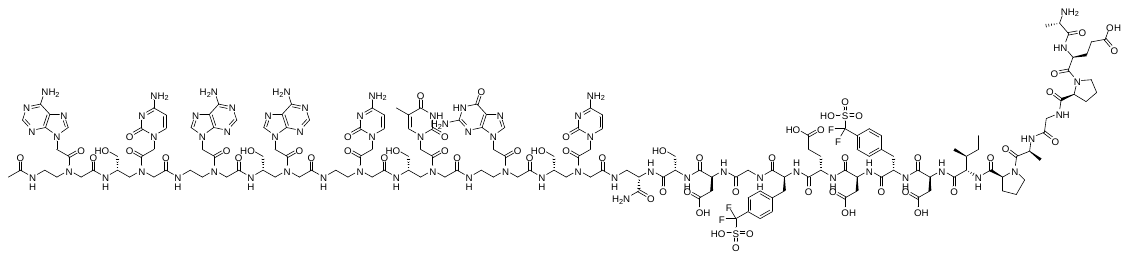

**Chemical Formula:** C_165_H_217_F_4_N_65_O_60_S_2_, **Exact Mass**: 4208.53, **Molecular Weight**: 4211.06.

**MALDI-TOF**; m/z found: 4211.44.

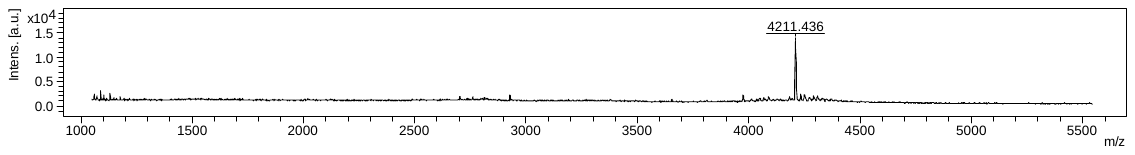

**LCMS (ESI)**; RT= 1.41 min, [M+3H]^3+^: 1404.42, [M+4H]^4+^: 1053.58.

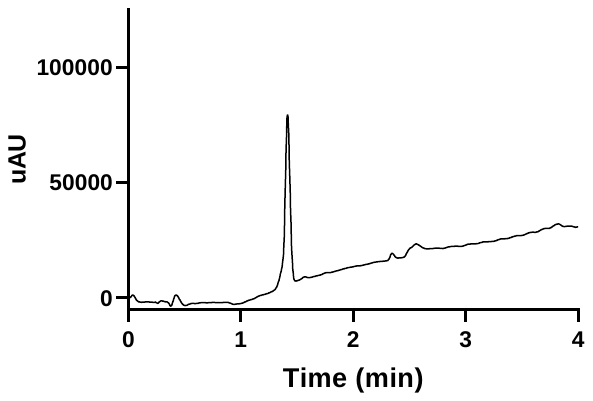

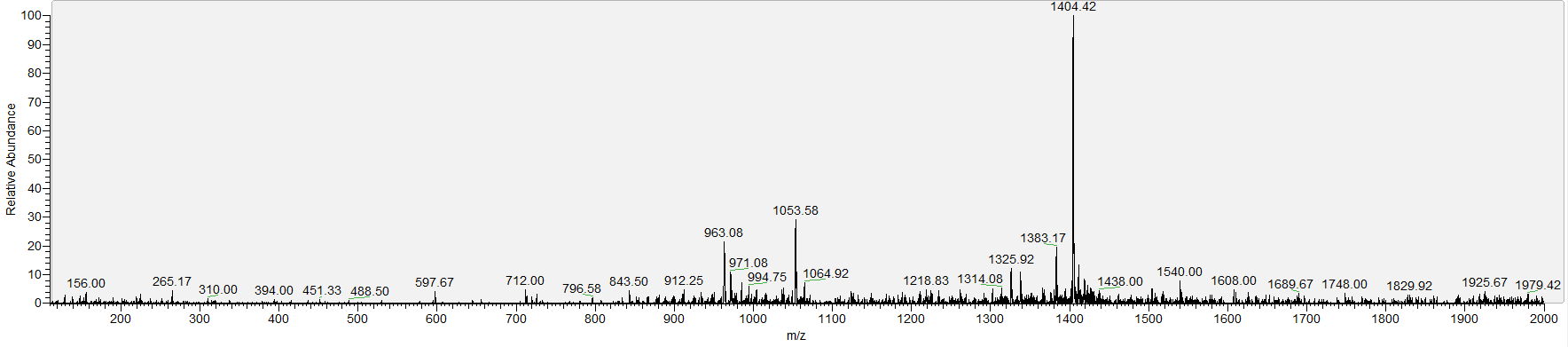

ii) **Exosite from different species**

**E23**

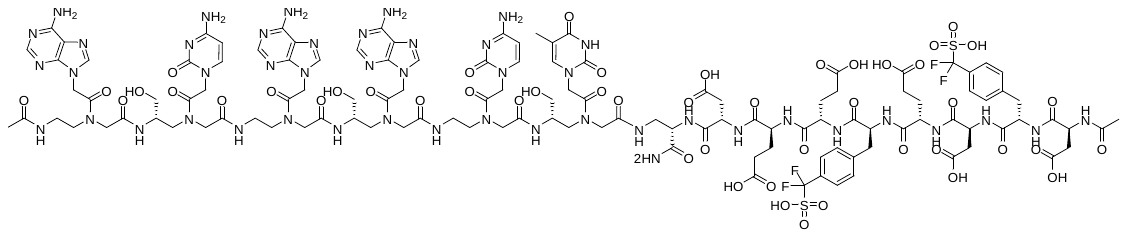

**Chemical Formula:** C_121_H_152_F_4_N_46_O_48_S_2_, **Exact Mass**: 3157.02, **Molecular Weight**: 3158.94.

**MALDI-TOF**; m/z found: 3159.22.

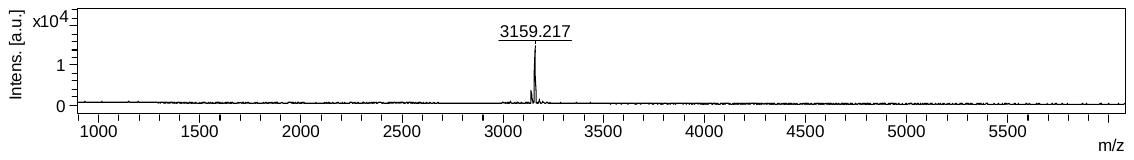

**LCMS (ESI)**; RT= 1.28 min, [M+3H]^3+^: 1053.92.

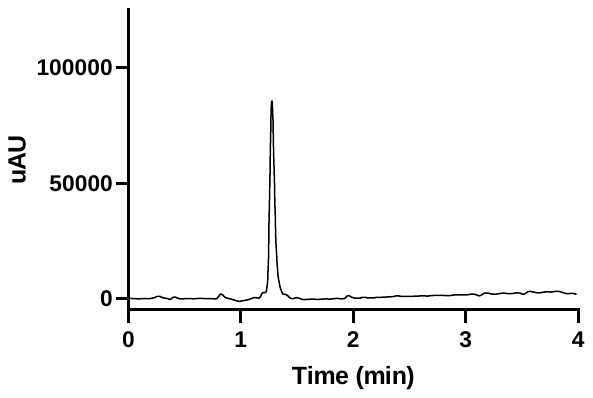

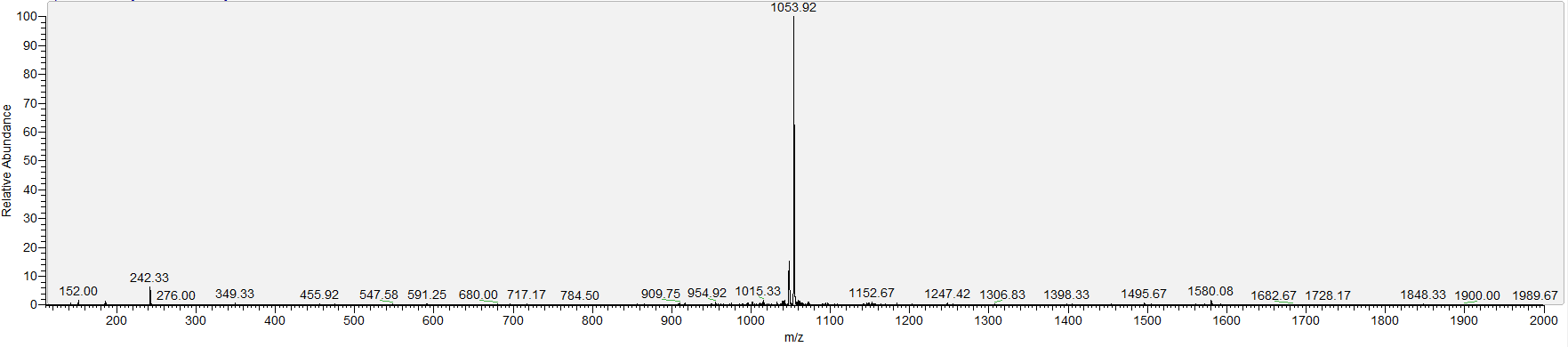

iii) **Active Site P3**

**A4**

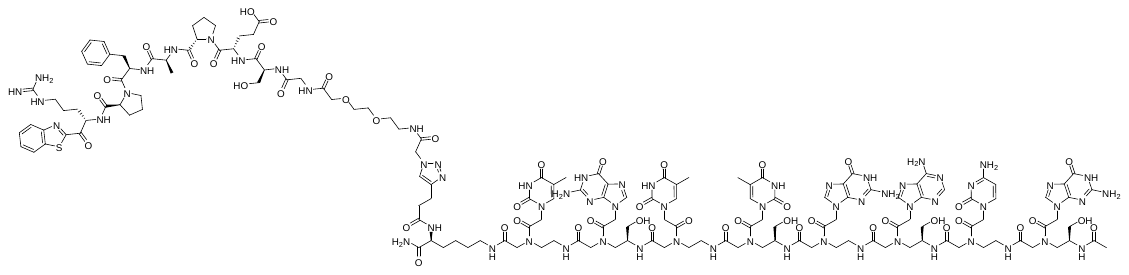

**Chemical Formula**: C_157_H_208_N_64_O_48_S, **Exact Mass**: 3789.55, **Molecular Weight**: 3791.85.

**MALDI-TOF**; m/z found: 3791.02.

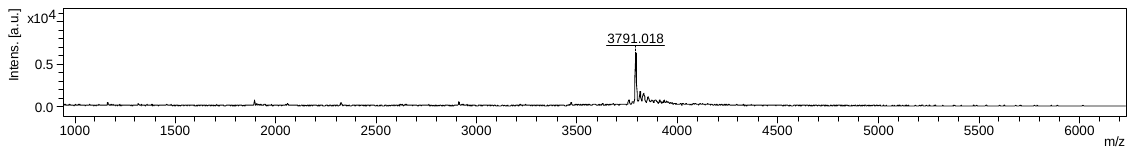

**LCMS (ESI)**; RT= 1.72 min, [M+3H]^3+^: 1264.67, [M+4H]^4+^: 948.92, [M+5H]^5+^: 759.25.

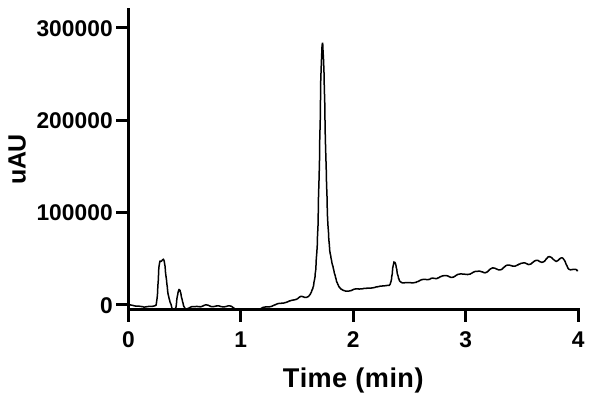

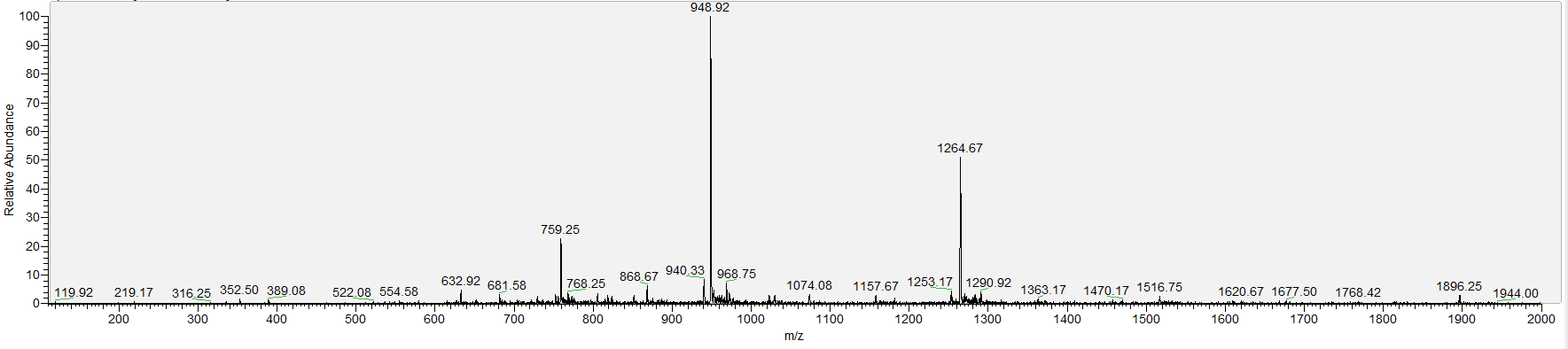

**A5**

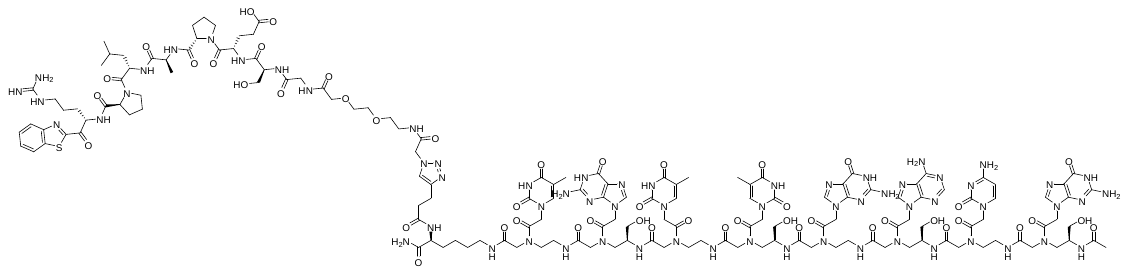

**Chemical Formula**: C_154_H_210_N_64_O_48_S, **Exact Mass**: 3755.57, **Molecular Weight**: 3757.83.

**MALDI-TOF**; m/z found: 3757.59.

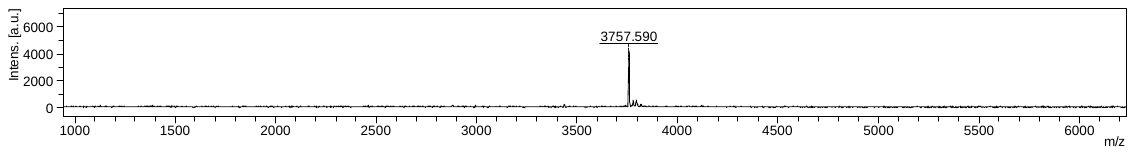

**LCMS (ESI)**; RT= 1.71 min, [M+3H]^3+^: 1253.25, [M+4H]^4+^: 940.33, [M+5H]^5+^: 752.33.

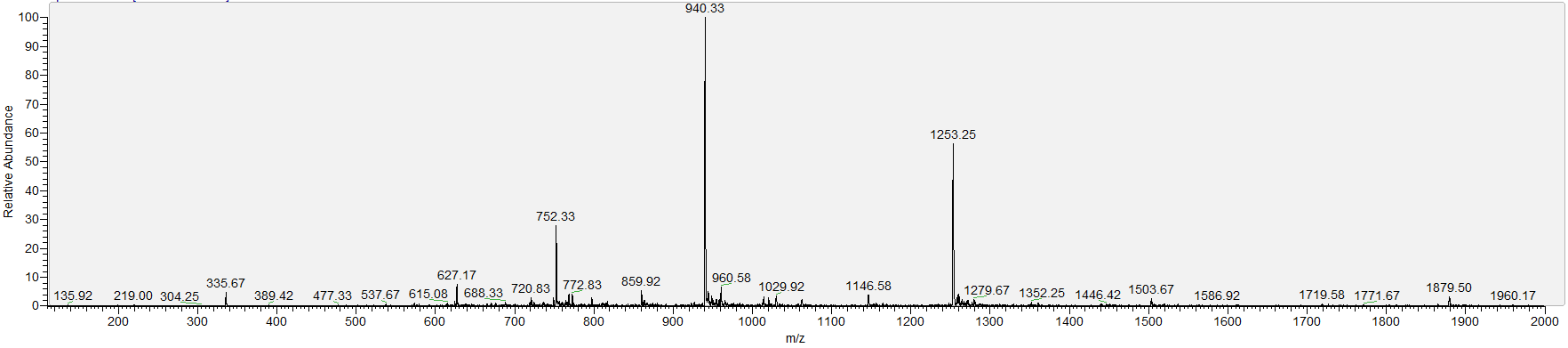

**A6**

**Chemical Formula:** C_152_H_206_N_64_O_49_S, **Exact Mass**: 3743.53, **Molecular Weight**: 3745.78.

**MALDI-TOF**; m/z found: 3745.70.

**LCMS (ESI)**; RT= 1.53 min, [M+3H]^3+^: 1249.25, [M+4H]^4+^: 937.25, [M+5H]^5+^: 750.08.

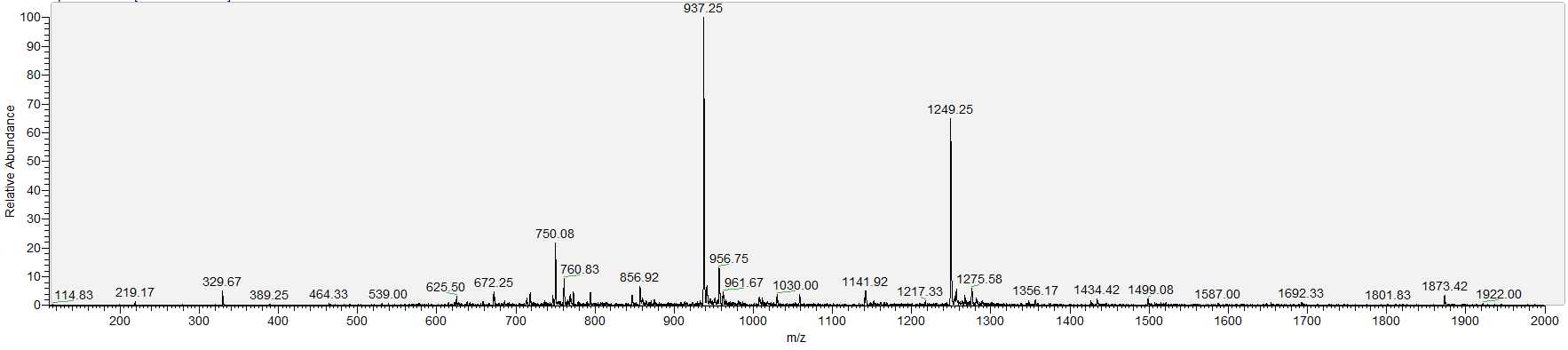

**A7**

**Chemical Formula**: C_150_H_202_N_64_O_48_S, **Exact Mass**: 3699.51, **Molecular Weight**: 3701.73.

**MALDI-TOF**; m/z found: 3701.51.

**LCMS (ESI)**; RT= 1.52 min, [M+3H]^3+^: 1234.58, [M+4H]^4+^: 926.33.

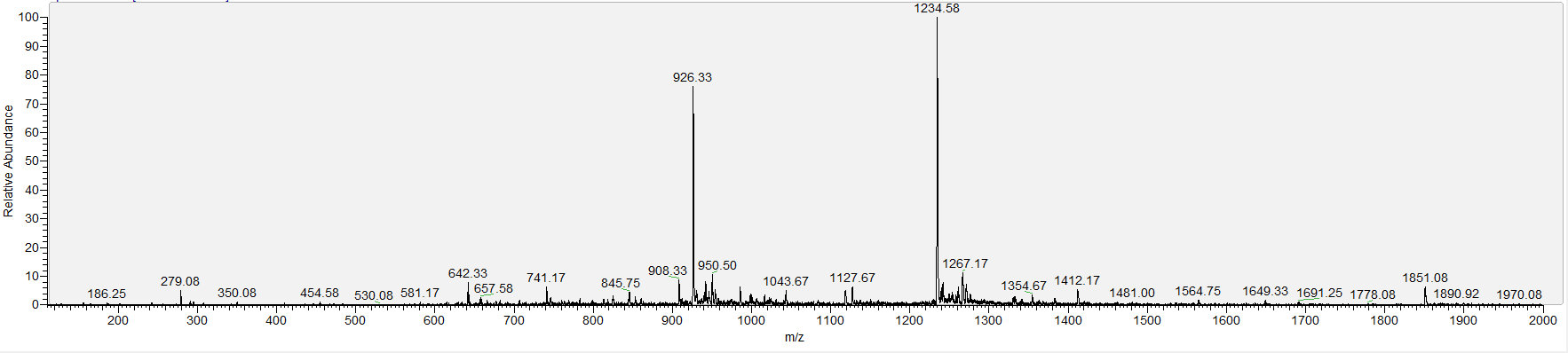

iv) **Antidote PNA**

Fmoc D monomer was prepared according to the procedure reported by Sugiyama *et al*.^3^

**AD1**

**Chemical Formula**: C_90_H_119_N_51_O_26_, **Exact Mass**: 2329.96, **Molecular Weight**: 2331.27.

**MALDI-TOF**; m/z found: 2330.85.

**LCMS (ESI)**; RT= 1.02 min, [M+2H]^2+^: 1166.33, [M+3H]^3+^: 778.00, [M+4H]^4+^: 583.83.

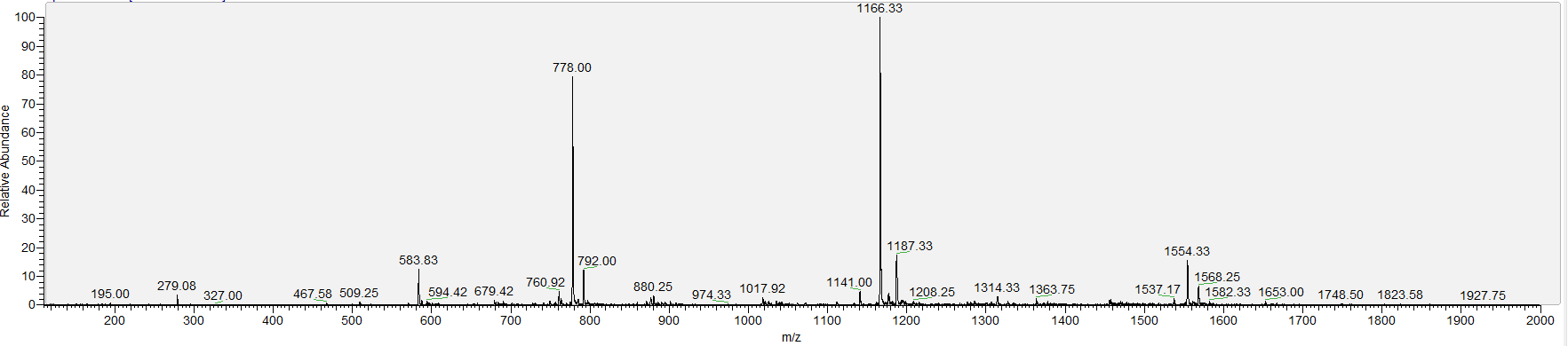

**HRMS**; [M+2H]^2+^: 1166.4845, [M+3H]^3+^: 777.9944, [M+4H]^4+^: 583.7455.

**AD2**

**Chemical Formula**: C_134_H_176_N_72_O_41_, **Exact Mass**: 3449.36, **Molecular Weight**: 3451.35

**MALDI-TOF**; m/z found: 3451.04.

**LCMS (ESI)**; RT= 1.22 min, [M+2H]^2+^: 1725.83, [M+3H]^3+^: 1151.25, [M+4H]^4+^: 863.83.

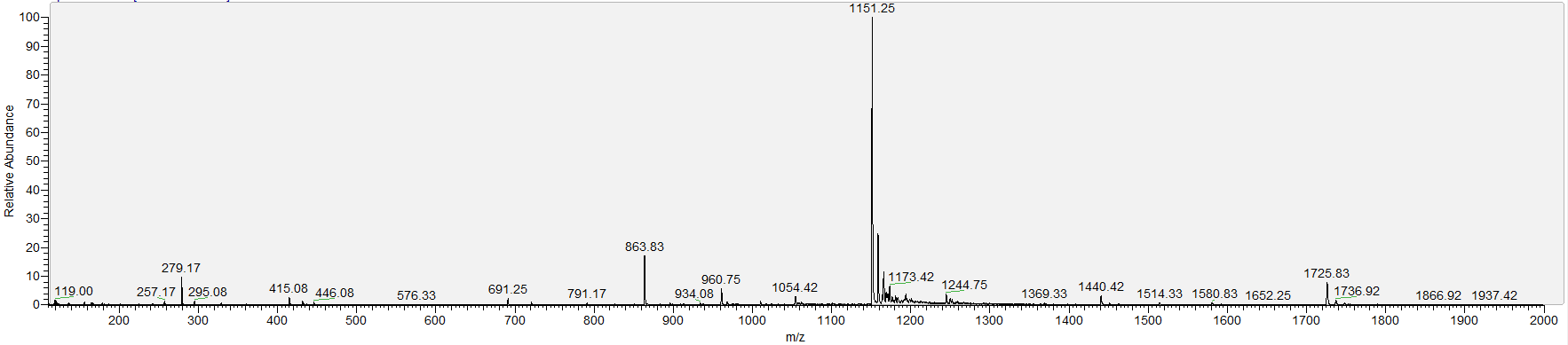

**HRMS**; [M+3H]^3+^: 1151.1405, [M+4H]^4+^: 863.6079, [M+5H]^5+^: 691.0870.

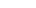

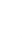

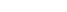

v) **PNA for SPR measurements**

**Biotin-PNA(8mer)**

**Chemical Formula**: C_105_H_141_N_51_O_31_S, **Exact Mass**: 2644.08, **Molecular Weight**: 2645.67.

**MALDI-TOF**; m/z found: 2645.35.

**LCMS (ESI)**; RT= 1.26 min, [M+2H]^2+^: 1323.25, [M+3H]^3+^: 882.67, [M+4H]^4+^: 662.25.

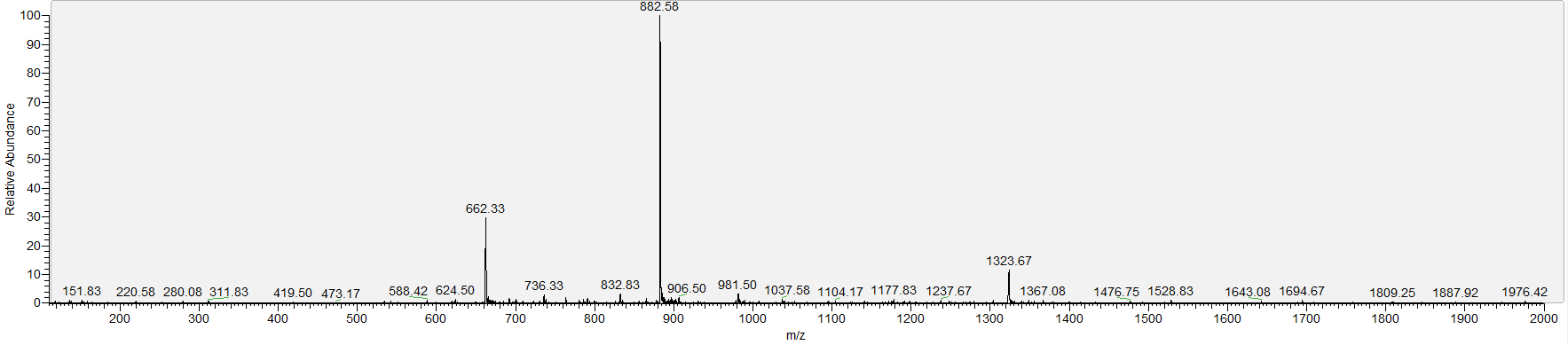

**PNA(8mer)**

**Chemical Formula**: C_99_H_132_N_48_O_32_, **Exact Mass**: 2505.02, **Molecular Weight**: 2506.45.

**MALDI-TOF**; m/z found: 2506.26.

**LCMS (ESI)**; RT= 1.15 min, [M+2H]^2+^: 1254.33, [M+3H]^3+^: 836.25, [M+4H]^4+^: 627.25.

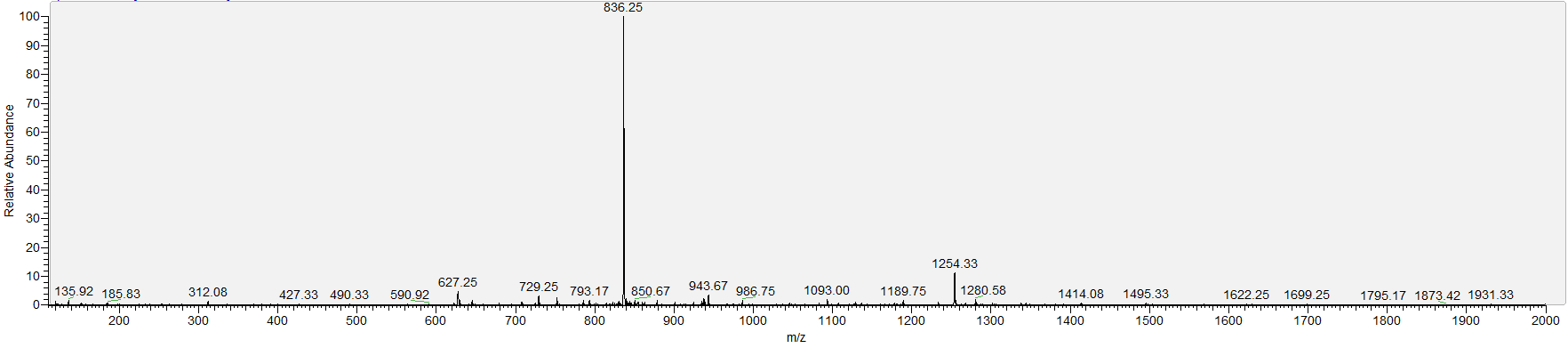

**PNA(6mer)**

**Chemical Formula**: C_71_H_92_N_34_O_24_, **Exact Mass**: 1804.70, **Molecular Weight**: 1805.73.

**MALDI-TOF**; m/z found: 1805.09.

**LCMS (ESI)**; RT= 1.14 min, [M+2H]^2+^: 903.42, [M+3H]^3+^: 602.58.

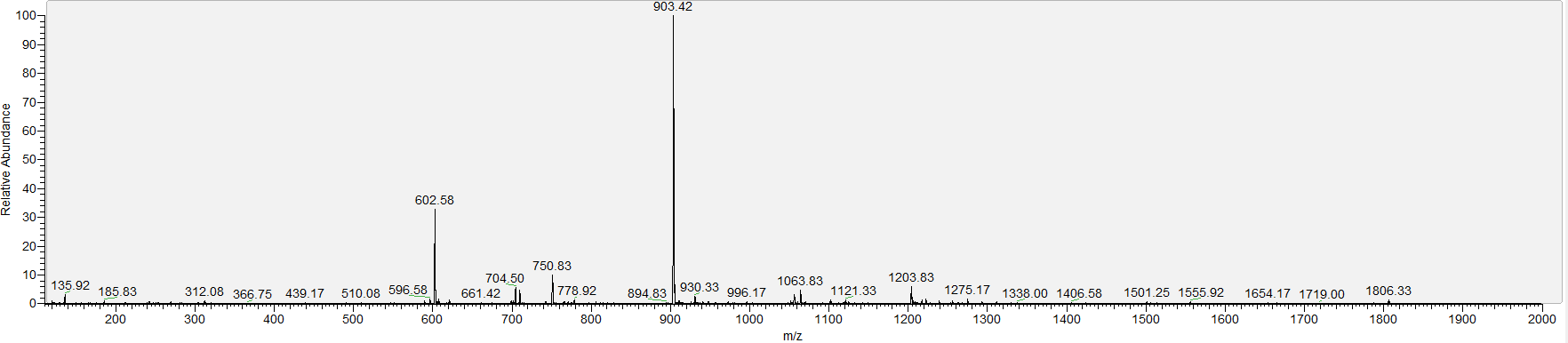

**PNA(4mer)**

**Chemical Formula**: C_47_H_62_N_24_O_15_, **Exact Mass**: 1202.48, **Molecular Weight**: 1203.17.

**MALDI-TOF**; m/z found: 1202.95.

**LCMS (ESI)**; RT= 0.46 min, [M+H]^1+^: 1203.25, [M+3H]^3+^: 602.25.

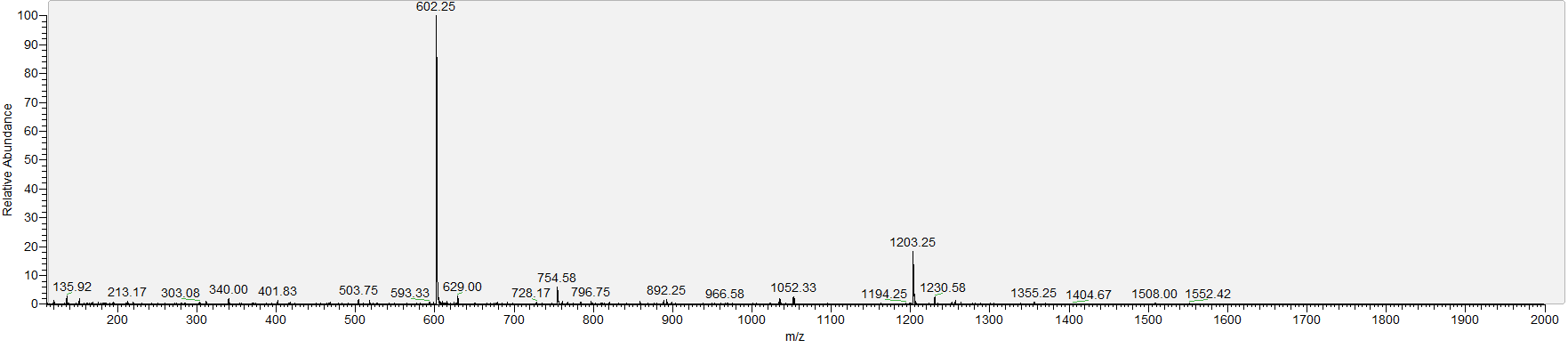

### 7. **Thrombin Inhibition Assay**

In vitro Inhibition of Human α-Thrombin. The inhibition of the activity of human α-thrombin (Haematologic Technologies, HCT0020) was followed spectrophotometrically using phe-Pro-Arg-Coumarin (synthesis described previously) as chromogenic substrate.

Inhibition assays were performed using 0.2 nM enzyme, 20 μM substrate, and increasing concentrations of inhibitor. The concentration of each inhibitor variant was determined using the absorbance of the PNA at 260 nm, measured by nanodrop. All reactions were carried out at 37 °C in 50 mM Tris-HCl pH 8.0, 50 mM NaCl, 1 mg/mL BSA in black 96-well microtiter plates (ref. 267342, ThermoFisher). Reaction progress was monitored by excitation at 339 nm and emission at 439 nm using a SpectraMax or Tecan Spark Plate Reader. Dose–response curves were used to determine the IC50 values using Prism 8.0 (GraphPad Software). For each inhibitor, the reactions were performed in triplicate, together with control reactions in the absence of enzyme. The initial velocity was calculated from the slope of the first 10 minutes of the assay. The curves were normalised to the well without inhibitor, where the initial velocity was set to 100% activity.

To convert IC50 values to Ki values, K_M_ was determined: The assay described above was performed without inhibitor but with variation of [substrate]. The Michaelis Menten plot was obtained, and K_M_ and Vmax were determined to be 2.416 μM and 435.8 respectively (R^2^: 0.98).

IC50 curves for compounds displayed in Figure 2e:

IC50 curves for compounds displayed in Figure 2f:

In the case of the antidote assay, the plate was removed from the plate reader at the desired time of addition (usually 30 minutes), 1 μL of antidote (100x) was added and reading was resumed. The recovery of activity (%) stated in the main text were calculated from the linear regressions of the slope (shown in red on the graph below) at the time stated (30 minutes or 90 minutes after antidote addition) compared to the linear regression of the initial activity of thrombin in the absence of inhibitor.

|  | **No Inhibitor** | **A8-E1+AD2 10 μM** | **A8-E1+AD2 15 nM** |
| --- | --- | --- | --- |
| **Best-fit values** |  |  |  |
| Slope | 573.4 | 212.5 | 101.4 |
| Y-intercept | -1064 | -8763 | -7271 |
| X-intercept | 1.856 | 41.23 | 71.69 |
| **Std. Error** |  |  |  |
| Slope | 2.098 | 1.330 | 0.9244 |
| Y-intercept | 34.69 | 122.0 | 171.5 |
| **Goodness of Fit** |  |  |  |
| R square | 0.9992 | 0.9930 | 0.9798 |
| Equation | Y = 573.4*X - 1064 | Y = 212.5*X - 8763 | Y = 101.4*X - 7271 |

### 8. **Fibrinogen Assay**

Human α-thrombin (Haematologic Technologies, HCT0020, final concentration 2.5 nM) was incubated with compound (final concentration 15 nM) at 37 °C for 30 minutes. Fibrinogen (final concentration 1 mg/mL) was added and absorbance at 288 nm was measured using a SpectraMax Plate Reader. All reactions were carried out at 37 °C in 50 mM Tris-HCl pH 8.0, 50 mM NaCl, 1 mg/mL BSA in clear 96-well microtiter plates (Greiner Bio-One, 650201).

In the case of the antidote assay, the plate was removed from the plate reader at the desired time of addition (usually 30 minutes), 1 μL of antidote (100x) was added and reading was resumed.

### 9. **Selectivity Assays**

The inhibition activity of **A1-E1** was tested against human FIIa, FXIa, and FXa (Haematologic Technologies), αFXIIa and plasma kallikrein (PK) (Enzyme Research Laboratories). Chromogenic assays were followed spectrophotometrically using specific substrates: 100 μM Tos-Gly-Pro-Arg-pNA (Chromozym TH; Roche) for FIIa; 500 µM Pyr-Pro-Arg-pNA (L-2145; Bachem) for FXIa; 500 µM Moc-D-Nle-Gly-Arg-pNA (L-1565; Bachem) for FXa; and 200 µM or 400µM D-Pro-Phe-Arg-pNA (Cayman Chemical) for αFXIIa or PK, respectively. The assay buffers were: 50 mM Tris-HCl pH 8.0, 50 mM NaCl for FIIa (0.2 nM); PBS pH 7.4 for FXIa (0.5 nM); 25 mM Tris-HCl pH 7.5, 100 mM NaCl, 5 mM CaCl_2_ for FXa (0.5 nM); 20 mM HEPES pH 7.6, 150 mM NaCl, 0.1% (w/v) PEG 8000, 0.01% (v/v) Triton X-100 for αFXIIa (4 nM); and 50 mM Tris-HCl pH 8.0, 150 mM NaCl for PK (0.25 nM). Bovine serum albumin (Sigma) was added to all buffers at 1 g/L. All reactions were initiated by the addition of the protease and carried out at 37 °C in 96-well flat-bottom microtiter plates. Reaction progress was monitored at 405 nm for 30 minutes (60 minutes for FXa and αFXIIa), on a multi-mode microplate reader (Synergy2, BioTek) with measurements taken every 5 minutes. All measurements were performed in duplicate. IC_50_ values were determined from the log-dose-response curves with Prism 9 (GraphPad Software).

### 10. **SPR Experiments**

SPR experiments were performed on a Biacore T200 instrument (GE Healthcare) at 25 °C in PBS-P+ buffer (10x stock from Cytiva Life Sciences, 28995084). Biotin-PNA(8mer) was immobilised on a Streptavidin Series S sensor chip (Cytiva Life Sciences, 29104992). Prior to immobilisation, the two channels were conditioned with 1 M NaCl in 50 mM NaOH. After stabilisation, the compound (solution in PBS-P+) was flowed over one of the flow cells of the sensor chip at a concentration of 50 nM at a flow rate of 10 μL min^−1^ with a response unit (RU) target of 500. Biotin-PNA(8mer) reached an RU value of 513.7. The system (not including the flow cells) was washed with 50% isopropanol in 1 M NaCl and 50 mM NaOH after each ligand injection. Kinetic measurements consisted of injections (association 400 seconds, dissociation 450 seconds, flow rate: 30 μL min^−1^) of decreasing concentration of PNA (4, 6 and 8mer) (2-fold cascade dilutions from the starting concentration). The chip was regenerated between cycles by one injection of regeneration solution (50 mM NaOH) for 10 seconds at a flow rate of 20 μL min^−1^, followed by a 10 second stabilisation period. Binding was measured as resonance units over time after blank subtraction, and the data interpreted using the Biacore T200 software, version 3.2. All measurements were performed in duplicate. The K_D_ values were calculated based on steady-state affinity (1:1 binding).

### 11. **Activated Partial Thromboplastin Time in vitro.**

Activated Partial Thromboplastin time (aPTT) measurements were performed on a BFT II benchtop analyser using the manufacturer’s instructions. Dade Actin™ FSL Activated PTT Reagent (Cat. No. 23-044-647) and calcium chloride solution (SMN/catalog number – 10446232 ORHO37) were both sourced from Siemens Healthcare Diagnostics Products, GmbH and lyophilised pooled human reference plasma (Pooled Norm. Cat. No. 00539) was purchased from Diagnostica Stago, Australia and New Zealand Victoria. Pooled human plasma was reconstituted as per manufacturer’s instructions (MQ, 30 minutes, RT). Pooled mouse plasma was prepared by collection of whole blood from 3-4 C57Bl6 mice (ABR, NSW) into sodium citrate (3.8%), with plasma isolated by centrifugation at 5,000x *g* for 15 minutes and stored on ice until required.

Human or mouse plasma was incubated with inhibitors at the indicated concentrations and pre-warmed to 37 °C. 50 μL of each plasma/inhibitor mixture was incubated with Actin™ FSL (50 mL) in a stirred reaction vessel for 3 minutes, prior to addition of 50 mL calcium chloride solution, to initiate coagulation. The time taken for fibrin clot formation was recorded in a semi-automated fashion using the BFT II Analyzer which employs a turbodensitometric detection technique.

### 12. **Ex vivo aPTT**

All procedures involving the use of animals were performed as approved by the University of Sydney Animal Ethics Committee (USyd AEC, protocol 2021/1912). C57Bl6 mice (25-30 g) were anaesthetised using a mixture of ketamine (125 mg/kg) and xylazine (12.5 mg/kg) (intraperitoneal [i.p.] delivery), then administered **A1**-**E1** as a single bolus delivered intravenously via the femoral vein at either 2.5 or 5.0 mg/kg. Blood was drawn from the IVC at the indicated times into citrate anticoagulant (3.8%), plasma isolated as described above for *in vitro* aPTT studies, and aPTT assessed via changes in plasma opacity at 405 nm using a ClarioSTAR plate reader fitted with dual injectors heated to 37 °C, using a modified version of the aPTT protocol described above. Briefly, injectors were primed for Dade Actin™ FSL Activated PTT Reagent (Line A) and calcium chloride solution (line B), and mouse plasma aliquoted in duplicate (25 μL) into wells of a Nunc 368-well polystyrene plate (Cat. No. Z723010, Sigma-Aldrich). Following injection of 25 mL Dade Actin™ FSL, the plate was mixed using the orbital shaking function for 2 seconds (500 rpm) and incubated for 182 seconds at 37 °C. At this time (designated t=0 s) 25 mL of calcium chloride solution was injected, the plate mixed as described above, and absorbance measurements taken at 405 nm for 360 intervals (22 flashes per well, 0.5 sec interval time). Clotting time was denoted by the timing of initial inflection point, denoting transition of plasma from transparent to opaque.

### 13. **Calibrated Automated Thrombogram (CAT)**

Normal lyophilised human pooled plasma (Pool Norm #00539, Diagnostica Stago S.A.S., Asnières-sur-Seine, France) was reconstituted and incubated for 30 mins at 37 °C. Vehicle and various inhibitor concentrations were incubated in plasma post 30-mins incubation. Thrombin assays were performed via a Hemker Calibrated Automated Thrombinoscope (Diagnostica Stago) using a Fluoroskan Ascent® plate reader (Thermo Scientific, MA, United States). All experiments were conducted in triplicate in 96-well microplates for fluorescence-based assays (M33089, Thermo Scientific, MA, United States) and calibrated using untreated plasma and a thrombin calibrator (#86192, Diagnostica Stago S.A.S., Asnières-sur-Seine, France). Thrombinoscope experiments were conducted following patented commercial protocols: in brief, each sample well was filled 20 μL PPP-reagent, containing a mixture of phospholipids and tissue factor (#86193, Diagnostica Stago S.A.S., Asnières-sur-Seine, France). 80 μL plasma (untreated/ treated) was then added to each of these wells, mixed using reverse pipetting, and the well plate was incubated by the plate reader at 37 °C for 10 minutes. Meanwhile, a FluCa-Kit (#86197, Diagnostica Stago S.A.S., Asnières-sur-Seine, France) containing Fluo-Buffer and Fluo-Substrate was warmed to 37 °C. Following incubation, the thrombinoscope dispenser was flushed, emptied, and filled with a FluCa mixture of the Fluo-Buffer and Fluo-Substrate. Twenty μL of the FluCa mixture were dispensed to each well containing plasma samples, commencing the coagulation reaction. Thrombin activity (nM) was measured over 1 hour, with thrombogram parameters including lag time (mins), velocity index (nM/min), time to peak (mins), peak height (nM), endogenous thrombin potential (ETP) (nM*min), and time to tail (min).

### 14. **Needle Injury Thrombosis Model**

C57Bl/6J mice were purchased from Australian BioResources (ABR, NSW, Australia) and housed at the Laboratory Animal Services (LAS) facility (the University of Sydney). All animals were housed in a 12-hour light/dark cycle with access to food and water ad libitum. For in intravital mouse studies, male mice aged between 5-8 weeks old (15g-20g) were used. All studies were approved by the University of Sydney Animal Ethics Committee (Protocol 2021/1912) in accordance with the requirements of the Australian Code of Practice for the Care and Use of Animals for Scientific Purposes.^5^

A clinical preparation of argatroban (Argatra/Exembol®) was purchased from Mitsubishi Tanabe Pharma (Germany) and prepared in sterile saline with 25% (vol/vol) of propylene glycol. Synthesized PNA inhibitors and PNA inhibitors + antidote solutions were prepared in sterile saline at a concentration of 2 mg/mL. Ketamine (150 mg/kg)- and xylazine (15 mg/kg)-anesthetized oxygen-supplemented C57BL/6J male mice (15-25 g) were subjected to an intravital needle-injury model, as previously described.^4^ Systemic injection of a DyLight 647 anti-GPIbβ antibody (X647 Emfret, Germany, 100 µg/kg) and Alexa 546-anti fibrin antibody (0.31 µg/kg) was performed prior to vessel injury, to monitor thrombus formation and fibrin generation, respectively. Argatroban (80ug/kg bolus; 40 ug/kg/min 60-minute infusion) was delivered via jugular catheter using a Harvard apparatus pump (Cat# 704504; Pump Elite 11 I/W Single Syringe Pump, NSW, Australia). Injections of PNA inhibitors or PNA inhibitors + antidote (5mg/kg bolus every 30 minutes) was delivered intravenously. 3-4 successive injuries were created in multiple vessels in each mouse from each treatment group. Following each injury, platelet thrombus formation and fibrin generation were monitored over a 15-minute period using a confocal intravital microscopy platform (Nikon A1R-si; objective: Apo LWD, ×40 magnification, 1.15 numerical aperture, water immersion; sequential excitation: 488-, 561-, and 638-nm lasers; emission: 525/50-, 595/50-, and 700/75-nm filters; using NIS Elements Advanced Research acquisition software). The microscope stage and objective were maintained at 37 °C throughout the experiment via a Peltier heater (OkoLab). Surface renders of confocal stacks representing thrombi from separate groups were generated using Imaris (Ver. 9.8 Bitplane AG, Zurich, Switzerland).

**Quantitative analysis of thrombus volume over time:**

NIS-Elements software (Nikon, Japan) was used to apply a threshold to the DyLight 649–conjugated anti-mouse GP1bβ antibody signal for each *xyz* stack in a time series and was used to calculate the volume for each time point.

**Quantitation of change in fibrin amount over time:**

The signal obtained from DyLight 649–conjugated anti-mouse GP1bβ antibody for each *xyz* stack in a time series was thresholded to create a mask. The total signal (AU) from the Alexa Fluor 546–conjugated anti-fibrin antibody within this mask (i.e., the fibrin signal within the thrombus) for each time point was then quantified using NIS-Elements software (Nikon, Japan).

**Statistical analysis:**

Statistical significance between multiple treatment groups was analyzed using a RM one-way analysis of variance (ANOVA) with Bonferroni post-testing. Statistical significance between 2 treatment groups was analyzed using a paired Student *t* test with 2-tailed *P* values (Prism software; GraphPAD Software for Science, San Diego, CA). Data are presented as means ± SEM where ‘n’ equals the number of independent experiments performed.

**Max intensity projections**

The max intensity projections for the *in vivo* data shown in Fig.3 and Fig.4 are displayed in the table below:

| **No inhibitor** | **Argatroban** | **A1-E1** | **A8-E1** | **A8-E1 + AD2** |
| --- | --- | --- | --- | --- |
| 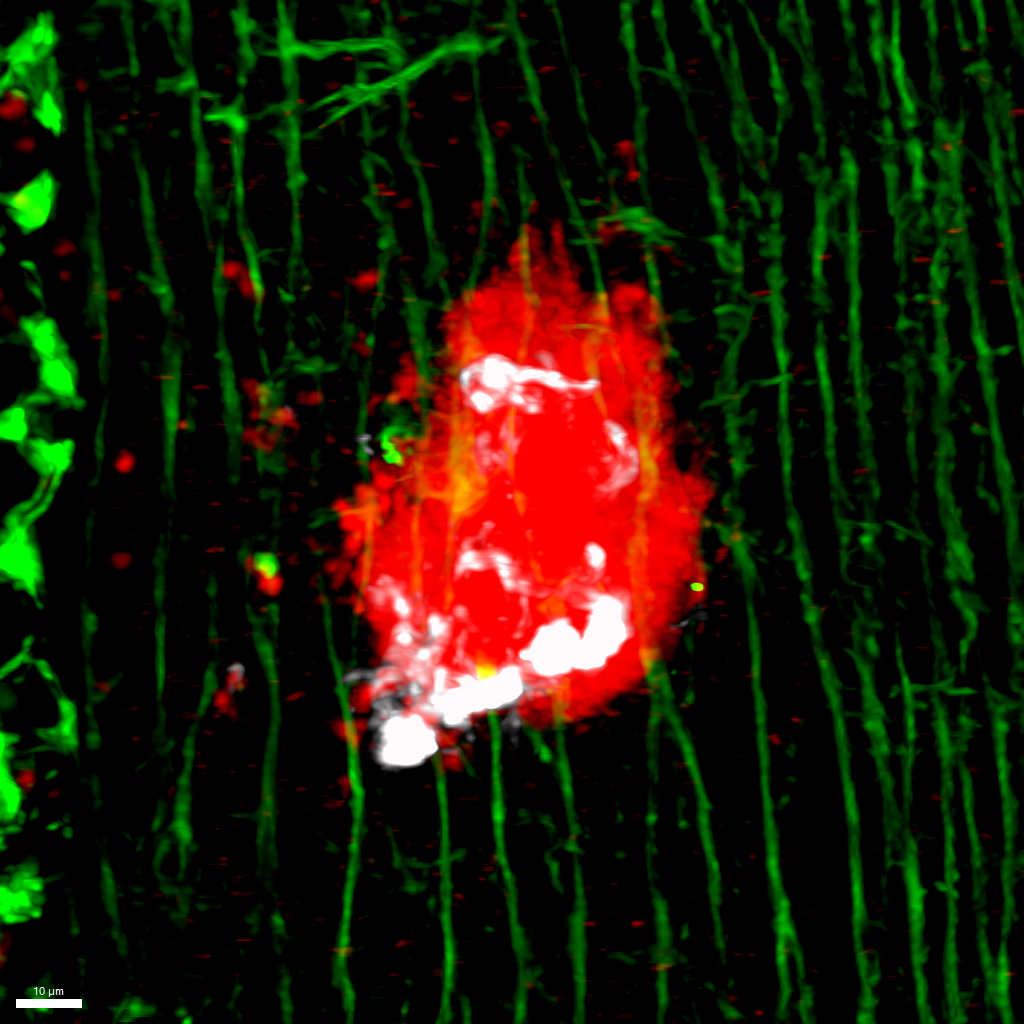 | 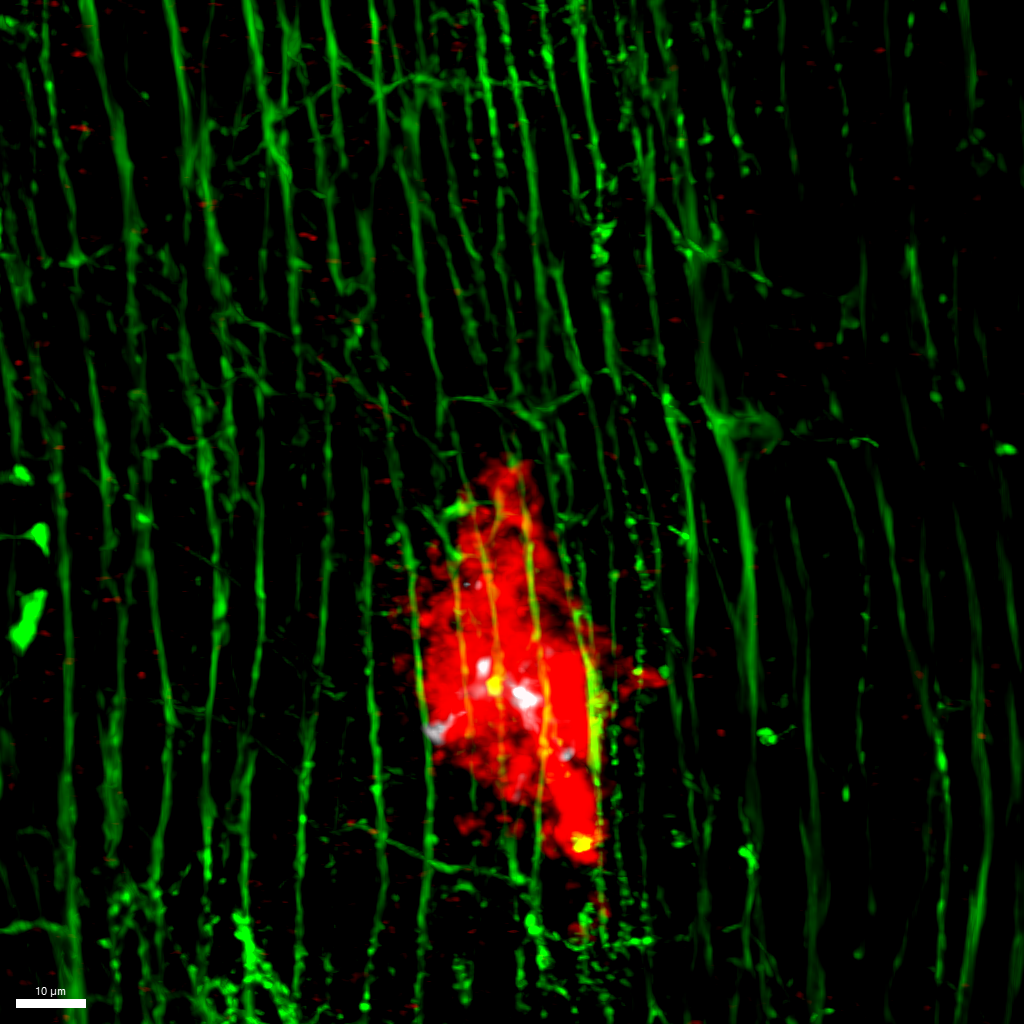 | 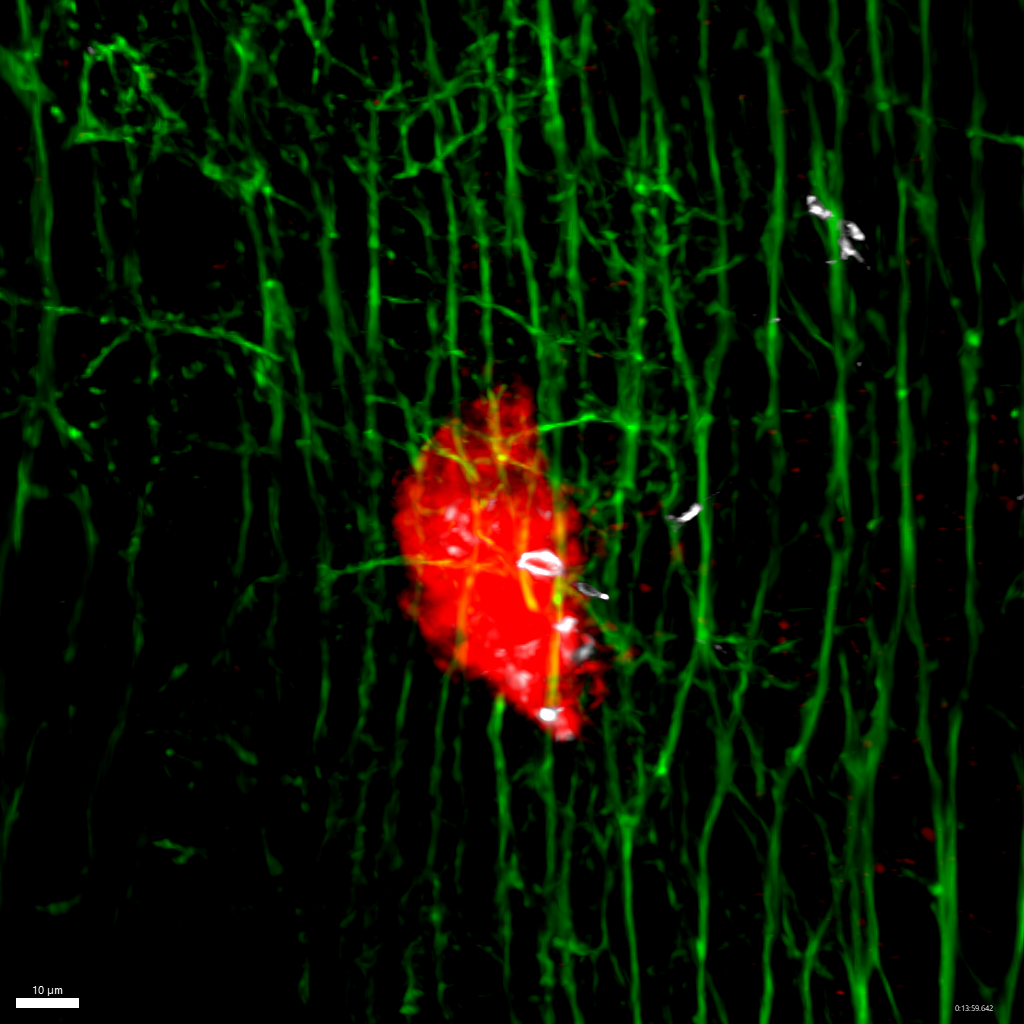 | 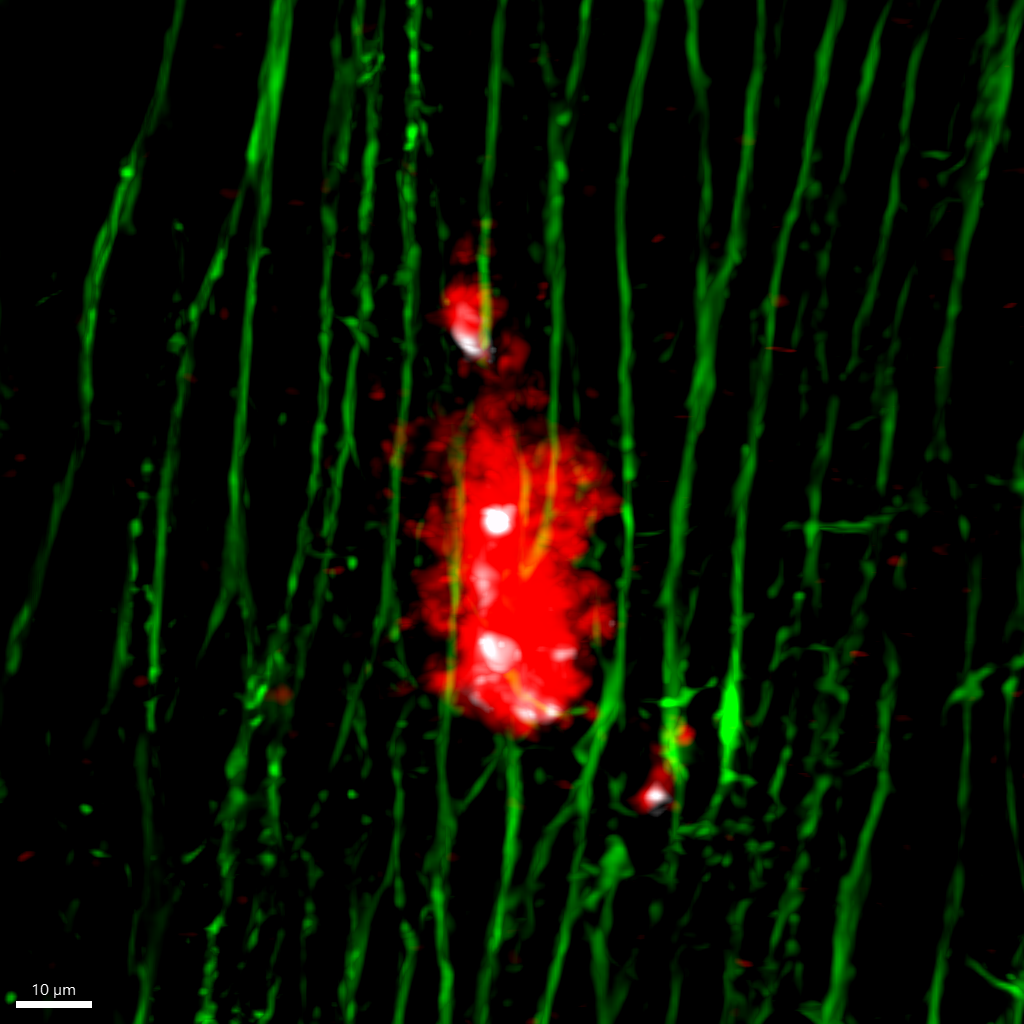 | 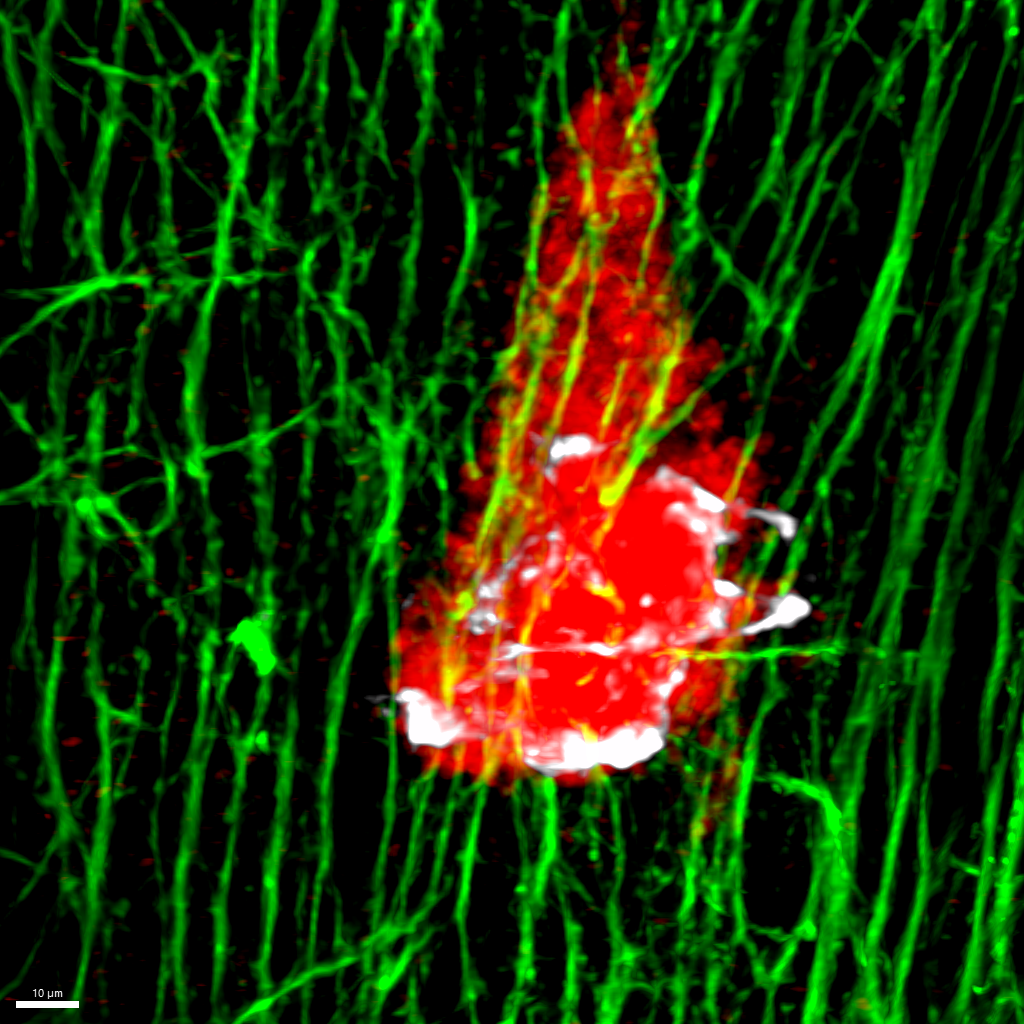 |

### 15. **References**

(1) Dowman, L. J.; Agten, S. M.; Ripoll-Rozada, J.; Calisto, B. M.; Pereira, P. J. B.; Payne, R. J. Synthesis and evaluation of peptidic thrombin inhibitors bearing acid-stable sulfotyrosine analogues. *Chemical Communications* **2021**, *57* (83), 10923-10926, 10.1039/D1CC04742F. DOI: 10.1039/D1CC04742F.

(2) Jakobsche, C. E.; McEnaney, P. J.; Zhang, A. X.; Spiegel, D. A. Reprogramming Urokinase into an Antibody-Recruiting Anticancer Agent. *ACS Chemical Biology* **2012**, *7* (2), 316-321. DOI: 10.1021/cb200374e.

(3) Sugiyama, T.; Hasegawa, G.; Niikura, C.; Kuwata, K.; Imamura, Y.; Demizu, Y.; Kurihara, M.; Kittaka, A. PNA monomers fully compatible with standard Fmoc-based solid-phase synthesis of pseudocomplementary PNA. *Bioorganic & Medicinal Chemistry Letters* **2017**, *27* (15), 3337-3341. DOI: <https://doi.org/10.1016/j.bmcl.2017.06.015>.

(4) Agten, S. M.; Watson, E. E.; Ripoll-Rozada, J.; Dowman, L. J.; Wu, M. C. L.; Alwis, I.; Jackson, S. P.; Pereira, P. J. B.; Payne, R. J. Potent Trivalent Inhibitors of Thrombin through Hybridization of Salivary Sulfopeptides from Hematophagous Arthropods. *Angewandte Chemie International Edition* **2021**, *60* (10), 5348-5356. DOI: <https://doi.org/10.1002/anie.202015127>.

(5) Australian Code for the Care and Use of Animals for Scientific Purposes 8th Edition (2013)
